# MultiNicheNet: a flexible framework for differential cell-cell communication analysis from multi-sample multi-condition single-cell transcriptomics data

**DOI:** 10.1101/2023.06.13.544751

**Authors:** Robin Browaeys, Jeroen Gilis, Chananchida Sang-Aram, Pieter De Bleser, Levi Hoste, Simon Tavernier, Diether Lambrechts, Ruth Seurinck, Yvan Saeys

## Abstract

Dysregulated cell-cell communication is a hallmark of many disease phenotypes. Due to recent advances in single-cell transcriptomics and computational approaches, it is now possible to study intercellular communication on a genome- and tissue-wide scale. However, most current cell-cell communication inference tools have limitations when analyzing data from multiple samples and conditions. Their main limitation is that they do not address inter-sample heterogeneity adequately, which could lead to false inference. This issue is crucial for analyzing human cohort scRNA-seq datasets, complicating the comparison between healthy and diseased subjects.

Therefore, we developed MultiNicheNet (https://github.com/saeyslab/multinichenetr), a novel framework to better analyze cell-cell communication from multi-sample multi-condition single-cell transcriptomics data. The main goals of MultiNicheNet are inferring the differentially expressed and active ligand-receptor pairs between conditions of interest and predicting the putative downstream target genes of these pairs. To achieve this goal, MultiNicheNet applies the principles of state-of-the-art differential expression algorithms for multi-sample scRNA-seq data. As a result, users can analyze differential cell-cell communication while adequately addressing inter-sample heterogeneity, handling complex multifactorial experimental designs, and correcting for batch effects and covariates. Moreover, MultiNicheNet uses NicheNet-v2, our new and substantially improved version of NicheNet’s ligand-receptor network and ligand-target prior knowledge model.

We applied MultiNicheNet to patient cohort data of several diseases (breast cancer, squamous cell carcinoma, multisystem inflammatory syndrome in children, and lung fibrosis). For these diseases, MultiNicheNet uncovered known and novel aberrant cell-cell signaling processes. We also demonstrated MultiNicheNet’s potential to perform non-trivial analysis tasks, such as studying between- and within-group differences in cell-cell communication dynamics in response to therapy. As a final example, we used MulitNicheNet to elucidate dysregulated intercellular signaling in idiopathic pulmonary fibrosis while correcting batch effects in integrated atlas data.

Given the anticipated increase in multi-sample scRNA-seq datasets due to technological advancements and extensive atlas-building integration efforts, we expect that MultiNicheNet will be a valuable tool to uncover differences in cell-cell communication between healthy and diseased states.

## Introduction

Tight coordination between different cells is required to maintain homeostasis in multicellular organisms. Consequently, dysregulation of cell-cell communication can lead to disease phenotypes. Studying cell-cell interactions is thus essential for an improved understanding of fundamental tissue biology and disease pathophysiology. Advances in single-cell and spatial transcriptomics are now providing opportunities to address this need through their ability to generate molecular profiles of cells within a tissue^1^. However, deciphering cell-cell communication from these profiles requires dedicated analysis approaches. As a result, several computational tools have been developed for this task^2, 3^.

Most tools infer cell-cell communication by predicting protein ligand-receptor interactions between pairs of cell clusters^4–7^ (typically annotated as cell types) or cells^8, 9^. They first estimate which ligands are expressed by one cell type (the sender) and which receptors by another (the receiver). They then link the expressed ligands to the expressed receptors if an interaction between a ligand and receptor is documented in a database of cognate ligand-receptor pairs. Differences between these ligand-receptor inference tools involve the choice of the used ligand-receptor database and the statistical procedure by which they infer the interaction likelihood based on the gene expression levels of the ligand and receptor^10^. For example, CellPhoneDB^4^ and CellChat^7^ use a permutation approach to prioritize ligand-receptor interactions based on cell-type specificity. Whereas these tools provide a comprehensive overview of expressed ligand-receptor pairs, they typically return an extensive list of interactions. As a consequence, it can be hard to decide which interactions are the most vital ones in the system of interest. Another limitation is that RNA-level co-expression of ligand and receptor does not necessarily guarantee that the ligand and receptor interact physically. Finally, they might overlook essential interactions that are weakly expressed or not cell-type specific.

Some other tools approach the cell-cell communication inference problem differently by incorporating downstream signaling of ligand-receptor interactions^11, 12^. We previously published NicheNet^11^, which predicts downstream affected target genes of expressed ligand-receptor pairs by combining the expression data of interacting cells with a model of ligand-target regulatory potential. This prior knowledge model is calculated through network-based data integration of signaling and gene regulatory networks. Contrary to the tools that prioritize ligand-receptor pairs based on cell-type specificity, NicheNet prioritizes expressed pairs according to how strongly their predicted targets are enriched in the receiver cell type (their so-called ligand activity). A high ligand activity might thus suggest that the ligand and receptor are not only expressed but also functionally interacting, making that ligand-receptor interaction an attractive candidate as a critical interaction in the biological process of interest. An additional benefit of NicheNet is that it generates hypotheses about the effects of ligand-receptor interactions on the receiver cell type (i.e., which are the specific targets they induce). But, the quality of the predictions critically depends on the accuracy of the prior knowledge used to link the ligand-receptor pairs to their target genes. We can expect false positive predictions if the signaling effects of ligand-receptor pairs were determined through experiments performed in different cell types and contexts than the ones of interest for the user’s analysis. False negative predictions are likely to occur for ligand-receptor pairs that are not well studied and for which there is thus a lack of knowledge on the downstream effects. Nevertheless, our previous benchmark^11^ demonstrates that NicheNet can accurately predict active ligands given an observed gene expression signature after *in vitro* ligand stimulation. The accuracy of ligand activity predictions is harder to systematically assess *in vivo*, but several studies reported experimental *in vivo* validation of some of the top predictions^13–17^.

Although both types of approaches have limitations, they have been applied successfully to study both communication in steady state and differences in communication between conditions^7, 11, 18^. Similarly to their application to steady-state data, the output of both approaches on the differential cell-cell communication inference task should be interpreted differently. The first class of methods returns ligand-receptor pairs of which one or both members are differentially expressed (DE) between the conditions. But, as mentioned above, differential RNA expression does not imply that an interaction is differentially active between conditions. In contrast, NicheNet predicts “differentially active” ligand-receptor interactions for which prior knowledge supports that they could function upstream of the DE genes in a receiver cell type of interest. This means that ligand-receptor interactions should not be DE themselves, increasing the chance of improperly prioritizing non-DE interactions. Improper prioritization is then likely to happen when these interactions would share signaling effects with a DE ligand-receptor pair that is truly pivotal in the process of interest. Even though ligand-receptor pairs should not necessarily be DE to be differentially active, including DE information might thus help in reducing false positives and identifying a more limited number of interactions for further experimental validation. Inferring differential cell-cell communication by prioritizing both differentially active and expressed ligand-receptor pairs is thus an exciting analysis strategy. However, Scriabin^19^ and our recent ad-hoc extension of NicheNet^14^ are currently the only tools to our knowledge with software support for such an analysis.

Moreover, current cell-cell communication tools suffer from additional limitations when applied to infer differential cell-cell communication from multi-sample scRNA-seq data. Biological replication in the form of multiple subjects (research animals or humans) is undoubtedly necessary to draw more robust conclusions. However, analysis strategies should consider the sample-to-sample biological and technical variability to avoid biased inferences. Several studies within the field of DE analysis from multi-sample scRNA-seq data highlighted that ignoring this variation can lead to false discoveries^20–22^. Running the classical cell-cell communication tools in their default mode on multi-sample data generates results after pooling all cells across samples^23–28^. This approach is thus statistically inadequate because it ignores sample-to-sample variation, and results might be skewed towards sample-specific interactions of samples with more cells^19^. Furthermore, one should ideally correct for confounding batch effects and other covariates (e.g., sex and age) if applicable to the data. Noteworthy, this pooling procedure is also suboptimal from a biological perspective because it ignores that cell-cell communication occurs within one sample. Lastly, visualizations showing inter-sample heterogeneity in cell-cell communication patterns are missing from the default output of the current tools. These issues are especially crucial for analyzing scRNA-seq data from clinical cohorts to investigate the role of cell-cell communication in human disease pathophysiology and therapy response.

Fortunately, several studies have already acknowledged these issues and performed a more appropriate ad-hoc analysis^18, 29–32^. Here, cell-cell communication is first analyzed per sample, followed by a statistical comparison of these cell-cell communication outputs between the different conditions of interest. However, dedicated stand-alone tools with a solid statistical basis and broad applicability are lacking. This lack of differential cell-cell communication tools for multi-sample scRNA-seq data is an important issue because of the expected rise in multi-sample datasets due to technological advances, for example, in sample multiplexing^33–38^. Sample multiplexing improves the throughput, decreases reagent costs, and minimizes the risk of introducing batch effects. In parallel to this evolution, more and more datasets are added to existing atlases in projects like the Human Cell Atlas^39, 40^, facilitated by effective integration algorithms^41, 42^. These atlases consist of several healthy and diseased samples of several tissues from multiple individuals. Deciphering the role of cell-cell communication in the pathogenesis of these diseases requires tools that can correct for the source of origin of the data and relevant clinical covariates.

Ideally, these tools should also be able to exploit the wealth of these multi-sample multi-condition datasets and tackle more complex questions than just pairwise comparisons. An exciting analysis would be to elucidate differential dynamics of cell-cell communication, such as comparing therapy response or disease progression between disease subtypes or diseases of the same tissue. Moreover, the multi-sample nature of these data also opens up opportunities to investigate the across-sample covariance structure between genes in different cell types. A few elegant novel methods like DIALOGUE^43^ and scITD^44^ started exploring this concept and aim to uncover so-called “multicellular gene expression programs”. These are gene sets from multiple cell types with co-varying expression across different samples. Tensor-cell2cell^45^ also extracts co-varying expression patterns, but it focuses explicitly on retrieving ligand-receptor interactions between sender-receiver cell-type pairs.

In summary, a few tools exist for expression- and activity-based differential communication analysis or for extracting intercellular communication patterns from multi-sample data. However, methods that connect both are lacking. This is an important issue because both aspects are intricately linked since robust differential analyses require multiple samples. Thus, there is a clear need for dedicated differential cell-cell communication tools that consider both the expression and activity of ligand-receptor pairs and can address the challenges and opportunities of multi-sample scRNA-seq data.

Therefore, we developed MultiNicheNet, a novel tool for differential cell-cell communication analysis from multi-sample multi-condition scRNA-seq data. MultiNicheNet builds upon the principles of state-of-the-art approaches for DE analysis of multi-sample scRNA-seq data. As a result, the algorithm considers inter-sample heterogeneity, can correct for batch effects and covariates, and can cope with complex experimental designs to address more challenging questions than pairwise comparisons. MultiNicheNet uses this DE output to combine the principles of NicheNet and ligand-receptor inference tools into one flexible framework. This enables the prioritization of ligand-receptor interactions based on differential expression, cell-type specific expression, and NicheNet’s ligand activity. Because the trustworthiness of the prioritization strongly depends on the quality of the used ligand-receptor database and ligand-target regulatory potential model, we updated the original NicheNet networks to obtain an improved version of both (“NicheNet-v2”).

We applied MultiNicheNet to scRNA-seq data of several tissues and diseases. On data of patients undergoing immunotherapy against breast cancer^23^, MultiNicheNet uncovered between- and within-group differences in cell-cell communication dynamics in response to therapy. Furthermore, MultiNicheNet’s predictions on data from patients with cutaneous squamous cell carcinoma^24^ or MIS-C (“multisystem inflammatory syndrome in children”)^18^ were corroborated by spatial co-localization or serum protein analyses. Finally, we applied MultiNicheNet to integrated lung atlas data to elucidate dysregulated cellular crosstalk in idiopathic pulmonary fibrosis while correcting batch effects^42, 46–49^. These applications demonstrate altogether that MultiNicheNet both retrieves known biology and generates novel hypotheses, including the possible identification of previously undescribed subgroups of patients, thus offering a novel way for patient stratification in cohort studies. In conclusion, MultiNicheNet is a flexible and broadly applicable differential cell-cell communication method that can reliably perform non-trivial analyses from multi-sample multi-condition single-cell transcriptomics datasets.

## Results

### MultiNicheNet prioritizes differential ligand-receptor-target communication patterns from multi-sample multi-condition scRNA-seq data

MultiNicheNet infers and prioritizes condition-specific ligand-receptor pairs and their target genes from multi-sample multi-condition single-cell transcriptomics data (**Figure 1a**). The main idea behind the prioritization strategy is to uncover essential interactions by considering several complementary aspects informative for cell-cell communication inference. We hypothesize that prioritization based on multiple relevant factors can increase the likelihood of inferring interactions vital for the communication process of interest. This contrasts with existing approaches that only use one aspect (e.g., expression or activity) for prioritization. As ideal ligand-receptor pairs, we consider pairs that are more strongly expressed in the condition of interest, are cell-type specific, are present in most samples of the condition of interest, and for which predicted target genes are enriched in the receiver cell type. MultiNicheNet uses the following criteria to achieve this prioritization: differential expression of the ligand in the sender cell type and its receptor(s) in the receiver cell type; cell-type and condition-specific expression of the ligand and its receptor(s); the fraction of samples in the condition of interest with sufficient expression of both ligand and receptor; and differential ligand activity in the receiver cell type (**Methods**). In the rest of this section, we describe how these criteria are calculated and used to obtain a final ranking.

**Figure 1.**
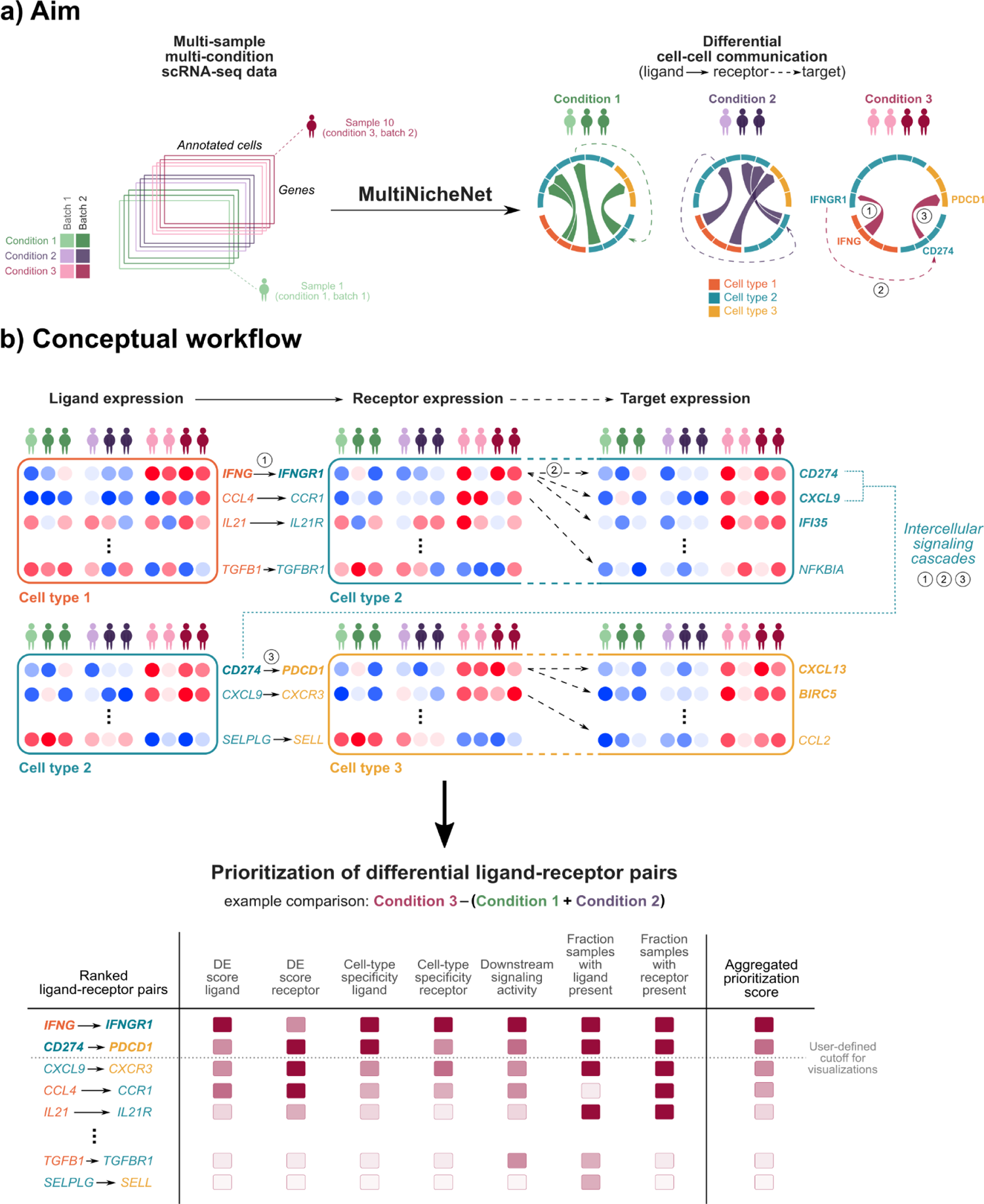
MultiNicheNet performs differential cell-cell communication analysis from multi-sample multi-condition single-cell transcriptomics data. a) MultiNicheNet starts from the raw counts of an annotated scRNA-seq dataset (each sample is here represented by one data rectangle). It infers differential cell-cell communication patterns that include ligand-receptor pairs (thick solid arrows) and predicted downstream target genes (dashed arrows). b) MultiNicheNet performs a ranking-based prioritization of differential cell-cell communication patterns based on multiple criteria, such as differential expression (DE) and downstream signaling activity (“ligand activity”). These criteria are summarized in an aggregated score, which can be used to define the most differential interactions for downstream visualizations (e.g., chord diagrams as shown in a)). Differential expression of ligands, receptors, and target genes is visualized through the blue-red color scale (blue: low, red: high). Ligand activity is here represented by the number of target genes predicted for each ligand-receptor pair (dashed arrows show receptor-target links). Some target genes encode for ligands that might be involved in interactions with other cell types, potentially forming “intercellular signaling cascades”.

MultiNicheNet starts from multi-sample multi-condition scRNA-seq data of putatively interacting cells and combines this with a ligand-receptor network and a corresponding NicheNet ligand-target regulatory potential model. As input, users have to provide the raw count matrix of the scRNA-seq dataset with a corresponding metadata table that indicates the following labels for each cell: the cell type, sample, and condition. If applicable, users can add batch and covariate information to correct for these factors. By default, MultiNicheNet will infer cell-cell communication between each pair of cell types, but users can restrict the analysis to pairs of interest, depending on the biological question or information on spatial co-localization. Finally, the user should indicate the comparison(s) they want to assess with MultiNicheNet (e.g., *condition 3* versus *conditions 1* and *2*). MultiNicheNet uses all this information to score each ligand-receptor pair between each sender-receiver cell type combination for all the prioritization criteria described above.

First, DE analysis for the requested comparison(s) is performed for each cell type. The DE analysis output is then used to define the strength of differential expression of ligands and receptors, and to determine the entire set of DE genes in a cell type. This set of DE genes is used to predict ligand activities for all ligands (as described in the original NicheNet paper^11^). To perform the DE analysis, MultiNicheNet uses pseudobulk aggregation followed by edgeR analysis, a state-of-the-art approach for DE analysis from multi-sample scRNA-seq data^21, 22, 50^ (**Methods**). In this approach, counts for all cells of a given cell type are first aggregated per sample into a pseudobulk count matrix. Next, each cell type’s pseudobulk matrix is used individually as input for DE tools designed for bulk RNA-seq analysis, such as edgeR^51^. As a result, we can compare the different conditions by using the sample as the experimental unit (in contrast to the cell, as frequently (mis)used in scRNA-seq DE analyses^21^). This approach thus handles inter-sample variation correctly and overcomes the problems associated with pooling all cells from all samples per condition. Moreover, the edgeR framework enables efficient handling of complex experimental designs and correcting for batch effects and covariates. In brief, state-of-the-art DE analysis is conducted to score each ligand-receptor interaction according to differential expression and ligand activity.

Secondly, the pseudobulk expression matrices are also used to calculate a metric that indicates a combination of cell-type and condition-specific expression of the ligand or receptor. This metric is the average normalized pseudobulk expression value per cell-type-condition combination (**Methods**). We introduced this criterion to avoid prioritizing non-cell-type-specific ligand-receptor pairs with slightly stronger differential expression over more cell-type-specific ligand-receptor pairs with slightly weaker differential expression. Subsequently, MultiNicheNet calculates per condition the fraction of samples in which the ligand-receptor pair is sufficiently expressed. We added this final criterion to reduce the chance of strongly prioritizing ligand-receptor interactions that are DE but are only expressed in a small subgroup of samples.

Finally, we scale each of these metrics for each ligand-receptor interaction across all sender-receiver cell type combinations and aggregate these scaled scores to obtain a final ranking of ligand-receptor interactions according to condition-specificity (**Figure 1b**) (**Methods**). We want to stress that this strategy results in a ranking of interactions, not in a defined set of interactions. We prefer this approach because it incorporates all the criteria mentioned above, but it will not eliminate interactions that score poorly on one of the criteria (which would occur in strategies that use thresholding on individual criterion scores to obtain a final set of interactions). We also want to emphasize the flexibility of the framework. Users can perform the aggregation in a weighted fashion to modify the influence of certain criteria during the prioritization (according to their insight into the data they are working on).

The MultiNicheNet software package does not only provide this prioritization framework, but it also provides possibilities for further downstream analyses and the generation of several intuitive visualizations. One downstream analysis is the prediction of specific target genes downstream of the prioritized differential ligand-receptor interactions with high ligand activity. In addition to inferring ligand-target links solely based on prior knowledge as described in the original NicheNet study, we can now exploit the multi-sample nature of the data to keep links only when there is expression correlation between the ligand-receptor pair and the target gene. Furthermore, we introduce a novel type of downstream analysis that we can only perform because of MultiNicheNet’s ability to predict target genes downstream of ligand-receptor interactions. Some target genes induced by certain ligand-receptor interactions encode for ligands or receptors themselves. Therefore, these might be involved in communication with other cell types, resulting in potential intercellular signaling cascades and feedback chains (**Figure 1b**). The MultiNicheNet software enables users to explore these putative “intercellular regulatory networks”.

As mentioned above, MultiNicheNet requires a ligand-receptor network and a corresponding NicheNet ligand-target model as input. Both have to provide accurate prior knowledge to ensure proper ligand-receptor prioritization and prediction of downstream target genes. The next section describes how we updated the ligand-receptor network and ligand-target model from the original NicheNet study.

### MultiNicheNet uses NicheNet-v2: an update of NicheNet’s prior knowledge that contains an up-to-date comprehensive ligand-receptor database and a more accurate model of ligand-target regulation

To improve ligand-receptor prioritization and target gene predictions, we substantially updated the NicheNet ligand-receptor network and prior knowledge model of ligand-target regulatory potential. In the rest of this manuscript, we will refer to the updated version of the prior knowledge networks as “NicheNet-v2” and to the original version as “NicheNet-v1”.

A major limitation of the NicheNet-v1 ligand-receptor network is that it contains many non-curated links predicted based on protein-protein interactions (PPIs) and gene annotation^11^. A second limitation is that it lacks relatively recently described ligand-receptor interactions that are documented in databases of more recently published cell-cell communication tools. To overcome these limitations, we constructed a novel ligand-receptor network for NicheNet-v2 that mainly comprises ligand-receptor interactions from the Omnipath intercellular communication database^52^ (see Methods). Omnipath is a comprehensive database that includes the different ligand-receptor databases used in multiple ligand-receptor inference methods, like the databases used in CellChat^7^ and CellPhoneDB^4^. In addition to Omnipath, we also incorporated additional ligand-receptor interactions from Verschueren et al^53^. Verschueren et al. described a set of novel ligand-receptor interactions of which at least one member is part of the immunoglobulin superfamily. We included these interactions because of their clinical relevance, certainly in tumor immunology, which is one of the research fields in which we expect MultiNicheNet to be used the most.

In addition to the ligand-receptor network, we also reworked the ligand-target regulatory potential model (see Methods, **Supplementary Note 1**, and **Supplementary Table 1**). We updated the existing data sources for which updates were available and included some novel data sources relevant to signaling pathways and gene regulation (e.g., KnockTF^54^). However, the main improvement of NicheNet-v2’s ligand-target model is the addition of *in vitro* experimentally determined target genes for 119 ligands. For this, we used the CytoSig^55^ database and the ligand treatment datasets described in the original NicheNet paper^11^ (see Methods). To assess whether the update results in a better prior model of ligand-target regulatory potential, we benchmarked the NicheNet-v2 model against the NicheNet-v1 model. As benchmark, we performed the same evaluation procedure as in the original NicheNet study^11^. In short, we determined a gold standard of ligand-target links as follows: we analyzed public transcriptomics datasets of cells before and after being treated by a ligand (*in vitro*), and we determined DE genes to obtain a set of true target genes of the ligand. Using this gold standard, we calculated two evaluation measures: target gene prediction and ligand activity prediction performance. To evaluate target gene prediction, we calculated how accurate the model predicts which genes are DE after ligand stimulation. To assess ligand activity prediction, we estimated how well the model predicts by which ligand the cells were stimulated. In addition to the ligand treatment datasets used in the NicheNet study, we also used the cytokine signatures provided by the CytoSig database^55^ as a second gold standard dataset. Because we also included the NicheNet and CytoSig ligand-treatment inferred ligand-target links to the model itself (see the previous paragraph), we had to design an evaluation strategy to avoid data leakage between model construction and model evaluation. In essence, we removed all CytoSig-based ligand-target links from the model before evaluating on the CytoSig gold standard and all NicheNet-based ligand-target links before evaluating on the NicheNet gold standard (see Methods). Using this procedure, we compared the predictive performance of NicheNet-v2 against NicheNet-v1 for both evaluation measures and gold standard datasets. We can conclude that NicheNet-v2 is substantially more accurate in target gene prediction than NicheNet-v1, whereas ligand activity accuracy is similar (**Supplementary Figure 1** and **Supplementary Table 2c-f**). These conclusions are identical based on both the NicheNet and CytoSig gold standard.

In summary, NicheNet-v2 provides a comprehensive and up-to-date ligand-receptor network and a more accurate ligand-target regulatory potential model than NicheNet-v1. We expect both improvements to reduce false positive and false negative predictions of ligand-receptor and ligand-target predictions in case studies. NicheNet-v2 provides both a human and mouse version of the ligand-receptor network and ligand-target model. In the rest of this paper, we will describe several MultiNicheNet case study analyses that we performed with this NicheNet-v2 model. However, users can now apply this new NicheNet-v2 model for regular NicheNet analyses as well.

### MultiNicheNet highlights critical pre-therapy cell-cell signaling patterns linked to therapy response in breast cancer patients

As a first case study, we applied MultiNicheNet to publicly available scRNA-seq data from breast cancer biopsies of patients receiving anti-PD1 immune-checkpoint blockade therapy^23^. Bassez et al. collected from each patient one tumor biopsy before anti-PD1 therapy (“pre-treatment”) and one during subsequent surgery (“on-treatment” - collected +/- nine days after the anti-PD1 treatment)^23^. Because of the importance of clonal expansion of T cells in defining the treatment response to immune checkpoint blockade therapy, Bassez et al. performed both single-cell transcriptomics and T cell receptor sequencing (scTCR-seq). Based on the scTCR-seq results, they identified one group of patients with clonotype expansion as response to the therapy (“E”) and one group with only limited or no clonotype expansion (“NE”).

Given this information, we applied MultiNicheNet to compare *pre-treatment* cell-cell communication between the E and NE patients. First, we focus on the interactions between macrophages and T cells (**Figure 2a** and **Supplementary Figures 2 & 3**). The two interactions with the highest prioritization scores were between PDL1 (*CD274*) from macrophages and PD1 (*PDCD1*) from CD4 T and CD8 T cells in E patients (**Supplementary Table 3a**). These interactions are characterized by both strongly differential expression and high ligand activity (see bubble heatmap of **Figure 2a**). In this case study, PDL1-PD1 signaling can be considered as a true positive prediction because we expect that PD1 signaling should be present in the tumor microenvironment of patients responding to anti-PD1 therapy^56^. In addition to PD1 signaling, MultiNicheNet inferred many other immune checkpoint interactions in the top 50 E-specific interactions between macrophages and T cells (**Supplementary Figure 2)**^56^. The most strongly E-specific checkpoint interactions from macrophages towards CD4 T cells are PDCD1LG2-PDCD1, NECTIN2-TIGIT, TNFRSF14-BTLA, LGALS3-LAG3, and CD86-CTLA4.

**Figure 2.**
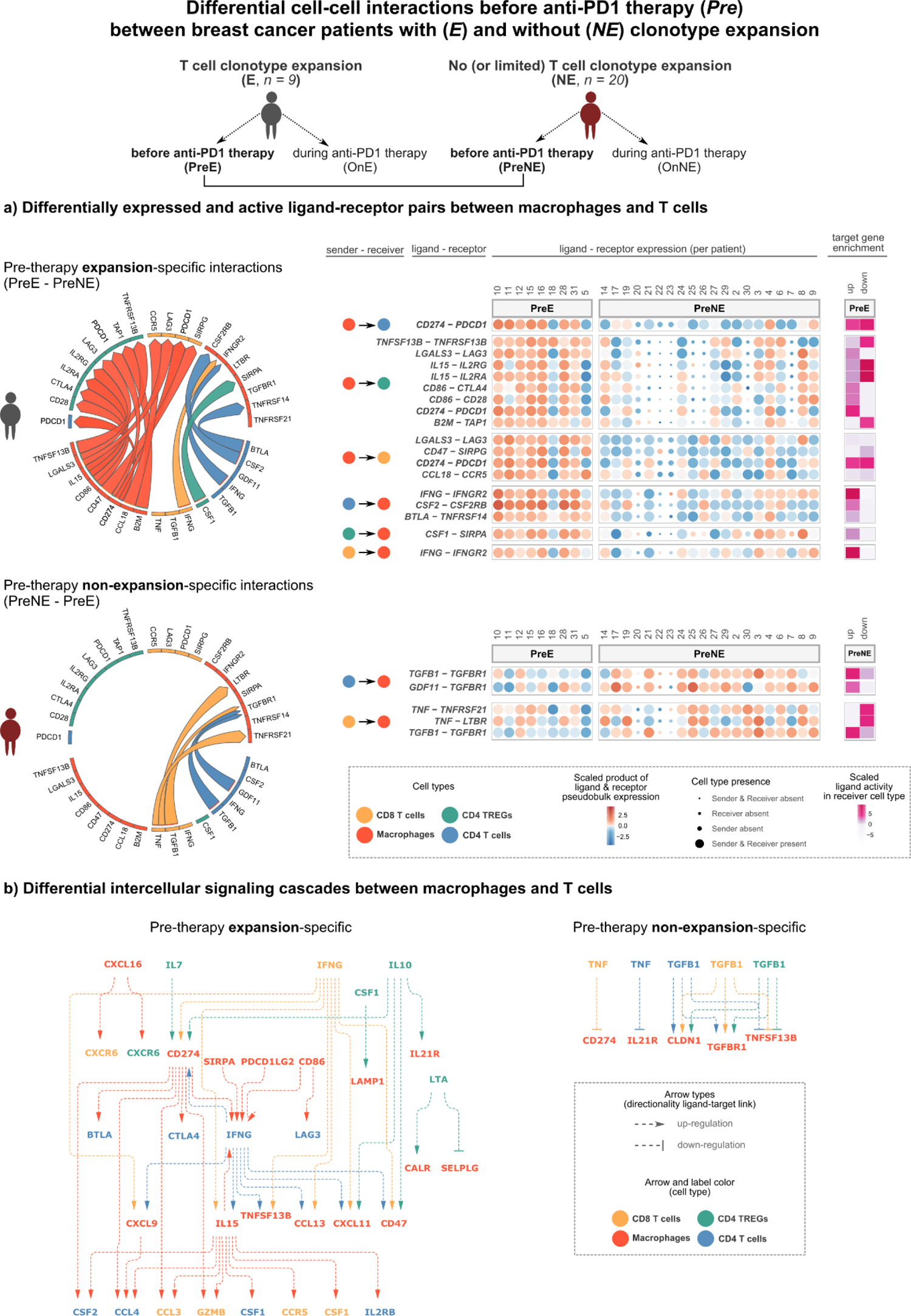
MultiNicheNet prioritizes differential pre-therapy communication patterns between breast cancer patients with and without T cell clonotype expansion in response to anti-PD1 immunotherapy. a) MultiNicheNet was applied to scRNA-seq from Bassez et al. to compare pre-therapy cell-cell communication between expander (preE) and non-expander patients (preNE). The most differential interactions between macrophages and T cells are shown and divided per patient group. Left: chord diagram visualization of the prioritized interactions. The arrowhead indicates the direction from sender to receiver cell type, and the color of the arrow indicates the sender cell type that expresses the ligand. Right: For each interaction, ligand-receptor pseudobulk expression and scaled ligand activity values are visualized. Scaled ligand activity values are calculated as z-score normalized NicheNet ligand activity values (Methods), calculated per receiver cell type. The higher these values, the more enriched target genes of a specific ligand are among the set of up- or downregulated genes in the preE or preNE group. The size of the dots indicates whether a sample had enough cells (>= 10) for a specific cell type to be considered for DE analysis. b) MultiNicheNet predicts an intercellular regulatory network showing potential intercellular signaling cascades specific for each patient group. This network consists of predicted ligand-target links for which the target gene encodes for a prioritized differential ligand or receptor. A predicted “up-regulatory” link indicates a positive across-patients expression correlation between the ligand-receptor and the prior-knowledge-supported target gene in the receiver cell type. A predicted “down-regulatory” link indicates an anti-correlation. To generate this plot, we considered the ligand-receptor pairs in the top 50 most differential interactions from macrophages to T cells and vice versa. Target genes should be among the 250 genes with the highest regulatory potential to be regulated by the specific ligand and should show expression correlation with the specific upstream ligand-receptor pair (Pearson or Spearman correlation > 0.33).

Checkpoint interactions from macrophages towards CD8 T cells in this top 50 are NECTIN2-TIGIT and LGALS3-LAG3. We also observe many chemokine interactions in the top 50 E-specific interactions (**Supplementary Figure 2)**: CXCL9-CXCR3 (macrophages - CD4 T cells); CCL7, CCL8, CCL13, CCL18, and CXCL11 towards CCR5 (macrophages - CD8 T cells); CXCL9, CXCL10, CXCL1, and CCL13 toward CXCR3 and CXCL16-CXCR6 (macrophages - CD4 regulatory T cells). Another strongly differential ligand in the E-specific tumor microenvironment seems to be IL15 (**Supplementary Figure 2)**. Interestingly, combining anti-PD1 therapy with IL15 signaling activation might potentially be a novel promising form of immunotherapy^57^.

If we now look at the interactions from T cells toward macrophages, we see that IFNG, CSF1, CSF2, and BTLA signaling are the most specific interactions in patients with clonotype expansion (**Figure 2a**). Like PD1 signaling, IFNG gamma signaling is one of the known predominant factors determining anti-PD1 therapy response^56^. Other top predicted E-specific signals include IL10, IL21, and IL7 (**Supplementary Figure 3**), which are all involved in modulating macrophage activity^58–60^. Additionally, interactions between CCR1 on macrophages and chemokines such as CCL3 and CCL5 (from CD8 T cells) and CCL4 (from CD4 T cells) seem to be specific for the tumor microenvironment of E patients (**Supplementary Figure 3**). Noteworthy, MultiNicheNet also predicts some potentially exciting interactions (e.g., IGF2L-IGFLR1 from T cells to macrophages) that have not been extensively studied before in the context of tumor microenvironment interactions and therapy response (**Supplementary Figure 3**).

Whereas we only discussed expansion-specific interactions in the previous paragraphs, MultiNicheNet also predicted some interactions specific for the patients without or with limited clonotype expansion. TGFB1, GDF11, and TNF are the most specific communication pathways between T cells and macrophages in NE patients (**Figure 2a**). Whereas we expected to retrieve anti-inflammatory signals like TGFB1 and GDF11 in non-expander patients^61^, retrieving TNF as NE-specific interaction was more surprising, even though it is known to have a complex dual role in the tumor microenvironment^62^. The bubble heatmap plot shows us that the predicted receptors of TNF are here LTBR and TNFRSF21 - two non-canonical receptors - and that these ligand-receptor interactions are not very clearly differentially expressed (**Figure 2a**). Their prioritization seems to be due to their strong predicted downregulatory activity in NE. However, the algorithm cannot discriminate between downregulatory activity in NE or upregulatory activity in E: it just indicates that TNF target genes are more strongly expressed in E patients. This might be due to the downregulation of these genes in NE patients. Still, it might also be possible that TNF is actually more active in E and that regulation of its activity occurs at the post-transcriptional level. Based on inspecting this plot, we might thus become less confident in this prediction compared to, for example, the prediction of TGFB1-TGFBR1: this is an interaction between a ligand and its canonical receptor, and both the interaction and its target genes are more strongly expressed in NE patients.

This last point illustrates the benefit of checking visualizations that show the data behind the predictions. The bubble heatmaps (as shown in **Figure 2a** and **Supplementary Figures 2 & 3**) show both the expression and activity of the ligand-receptor pairs. Moreover, they give the user insight into potential patient-to-patient heterogeneity. They also indicate if there were sufficient cells of a particular cell type in a sample to be included in the DE analysis. Furthermore, MultiNicheNet provides another visualization for users to explore which specific target genes are enriched in a receiver cell type (**Supplementary Figure 4a**). These ligand-target links are inferred based on prior knowledge in the same way as described in the original NicheNet paper. However, the user can also filter interactions based on expression correlation between the ligand-receptor pair and the target gene (**Supplementary Figure 4a-b**). We expect that target genes downstream of a ligand-receptor interaction will follow the same expression trend as the ligand-receptor pair. Consequently, outlier patients based on ligand-receptor expression will also be outliers in target gene expression (for example, patient 18; **Supplementary Figure 4a**). As the original NicheNet publication describes, we recommend that users verify the prior knowledge used to predict certain ligand-target links. Performing this verification for a subset of target genes downstream of PDL1-PD1 demonstrates that many edges in the intracellular signaling network are additionally supported by data sources newly added to NicheNet-v2 (**Supplementary Figure 5** and **Supplementary Table 3c**). Finally, MultiNicheNet provides the possibility to predict intercellular regulatory networks and compare them between the conditions of interest. These networks link prioritized ligands from one cell type to their ligand-encoding or receptor-encoding target genes in other cell types. The putative intercellular regulatory network between macrophages and T cells points to a potential interplay between IL15, IFNG, and PDL1 (*CD274*) in patients with clonotype expansion and connects many of the above-described signals (**Figure 2b**). Literature supports several links in this network (e.g., *CCR5* regulation by IL15^63^ and *CD274* regulation by IFNG^64^).

Although the previous paragraphs described the main interactions inferred among macrophages and T cells, we also investigated interactions among the other cell types in the tumor microenvironment (**Supplementary Note 3**). MultiNicheNet uncovered several interesting interactions involving dendritic cells (DCs), natural killer (NK) cells, and malignant cells. For example, MultiNicheNet retrieved a loss of antigen presentation in cancer cells in non-expander patients. Together with decreased PD-L1 and IFNG signaling, this is a known essential determinant of non-responsiveness to immune checkpoint blockade therapy^56^. Altogether, the results of this case study showcase that MultiNicheNet can prioritize both well-known and novel cell-cell signaling patterns linked to anti-PD1 therapy response in breast cancer patients.

### MultiNicheNet elucidates differences in therapy-induced communication changes between and within response groups

We here describe how MultiNicheNet can exploit the flexibility of generalized linear models in the pseudobulk-edgeR framework to handle complex multifactor experimental designs and address non-trivial questions. We applied MultiNicheNet to the same breast cancer data from Bassez et al. to compare *cell-cell interaction changes during anti-PD1 therapy (“on” versus “pre”)* between the E patients and the NE patients (**Figure 3a**)^23^. This analysis exemplifies how to study differential dynamics of cell-cell communication between conditions or patient groups. MultiNicheNet predicts a stronger increase in several immunomodulatory interactions (including chemotactic interactions) during anti-PD1 therapy in patients with clonotype expansion (**Figure 3a** and **Supplementary Table 3d**). Examples of increasing chemotactic interactions are CCL19-CCR7 between fibroblasts and B cells, CXCL12-CXCR4 between endothelial cells and B cells, and CXCL9-CXCR3 between macrophages/fibroblasts/DCs and (regulatory) CD4 T cells (**Figure 3a** and **Supplementary Table 3d**). These predictions suggest that T and B cells may be more strongly recruited into the tumor during anti-PD1 therapy, specifically in expander patients. Notably, previous research also demonstrated that the CXCR3 chemokine system is a key determinant of anti-PD1 therapy efficacy^65^.

**Figure 3.**
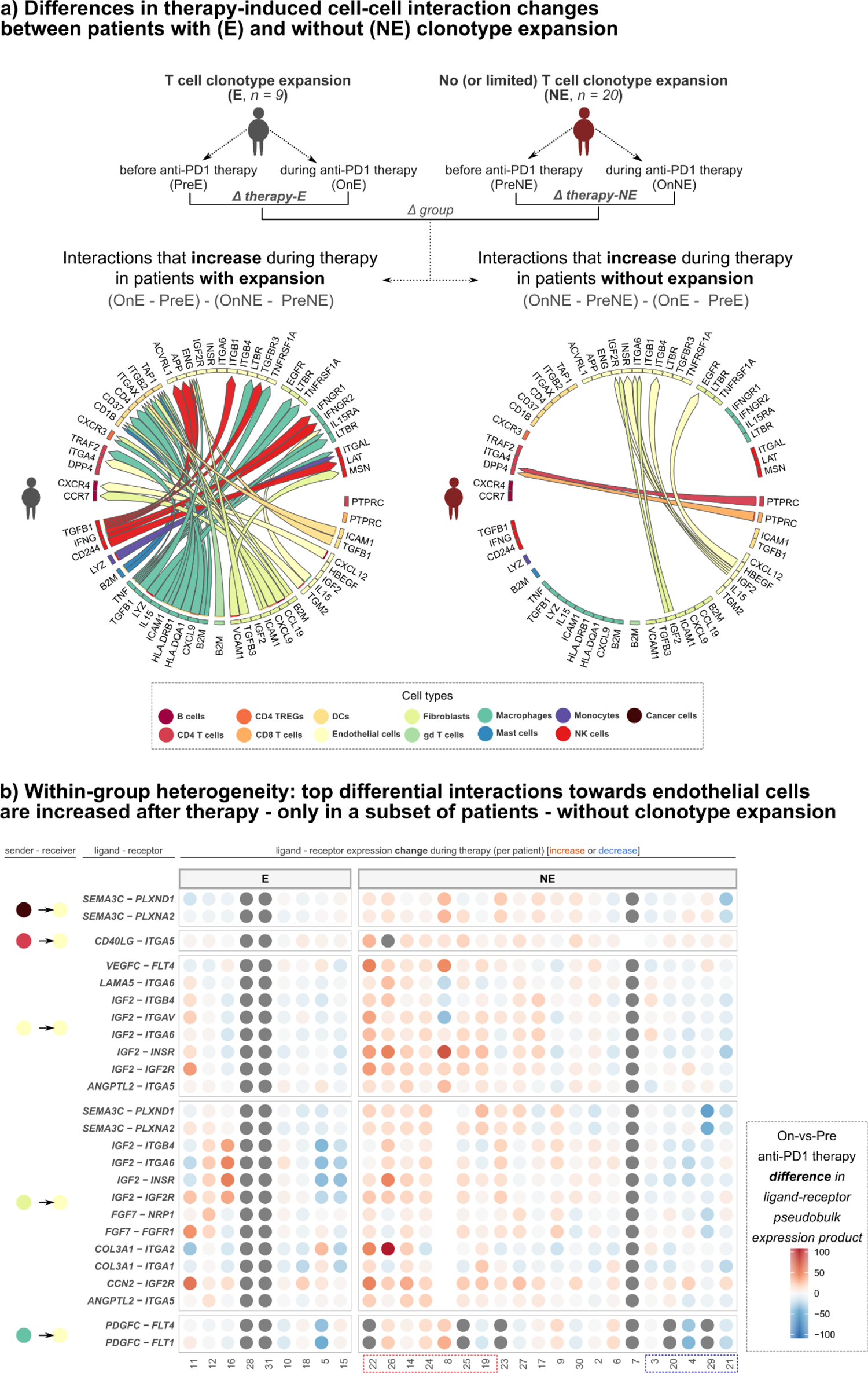
MultiNicheNet prioritizes differences in therapy-induced intercellular communication changes between and within response groups. a) MultiNicheNet was applied to scRNA-seq from Bassez et al. to compare on-therapy versus pre-therapy differences in cell-cell communication between breast cancer patients with T cell clonotype expansion in response to anti-PD1 therapy (expander - E) and patients without expansion (non-expander patients - NE). The top 50 differential ligand-receptor pairs are depicted in chord diagrams, divided into the expander-specific pairs (left) and non-expander-specific pairs (right). The arrowhead indicates the direction from sender to receiver cell type, and the color of the arrow indicates the sender cell type that expresses the ligand. b) Bubble plot showing the on-vs-pre therapy difference in ligand-receptor pseudobulk expression value (product of normalized expression) for each ligand-receptor pair and each NE patient. Dots colored in grey indicate the absence of the sender and/or receiver cell type in a certain sample. The red box at the bottom indicates the NE patients in which angiogenesis-related signals increased during anti-PD1 therapy, and the blue box indicates NE patients in which these signals were decreased.

In contrast to our expectations, several intercellular signaling patterns also increased during therapy in the *non-responder* patient group (**Figure 3a** and **Supplementary Table 3e**). Endothelial cells were one of the predominant receivers, together with fibroblasts and malignant cells (**Supplementary Table 3e**). The majority of molecules participating in interactions toward endothelial cells were previously shown to be involved in the regulation of angiogenesis (semaphorin-plexin, IGF2, FGF7, and PDGFC^66–69)^ or lymphangiogenesis (VEGFC, FLT4, ITGA1, ITGA2, and ITGA5^70, 71^)(**Figure 3b**). These processes play a role in determining the response to cancer immunotherapy, and a combination of anti-angiogenesis and immunotherapy has been proposed to work synergistically^72^.

Importantly, we can clearly see inter-patient heterogeneity in these (lymph)angiogenesis-related interactions in the NE group (**Figure 3b**). Even though these interactions seem to increase in expression and activity at the group level, this group-level increase appears to be driven by a subset of the NE patients (**Figure 3b** and **Supplementary Figure 6**). Moreover, we can even observe a decrease in these interactions in a small subset of NE patients. This demonstrates the benefit of MultiNicheNet’s visualizations in uncovering inter-patient heterogeneity.

Next, we further explored the potential biological and clinical relevance of these subgroups of non-responding patients. However, we could not link these patient groups to a relevant clinical subdivision such as breast cancer type or tumor architecture (hot-vs-cold). Nor could we connect this subdivision to the degree of clonotype expansion or relative cell type abundances^23^. Therefore, we assessed *pre-treatment* differences in intercellular signaling between the angiogenesis-increasing NE-subgroup and the angiogenesis-decreasing NE subgroup (patients indicated by the red box versus the blue box in **Figure 3b**). MultiNicheNet elucidates various interesting differential communication patterns. The group of patients with an increase in angiogenesis interactions show higher TNF signaling toward malignant cells, higher expression levels and activity of Annexin and multiple chemokines toward macrophages, and higher IL1 and ACKR1 signaling toward endothelial cells (**Supplementary Figure 7** and **Supplementary Table 3f**). On the contrary, the main characteristic of patients with a decrease in angiogenesis interactions is the presence of a diverse set of interactions toward endothelial cells. These interactions include TGF-beta, JAG-NOTCH, semaphorin-plexin, VEGFC, and IGF2 (**Supplementary Figure 8** and **Supplementary Table 3g**). Remarkably, these factors are also involved in (lymph)angiogenesis^73, 74^ and partially overlap with the interactions that increase during therapy in the angiogenesis-increasing group of NE patients. Whereas this analysis revealed various differential signaling pathways within NE patients, further research would be necessary to explain these differences and their importance.

In conclusion, the MultiNicheNet analysis of anti-PD1 therapy response revealed a therapy-induced increase in chemotactic interactions in expander patients and (lymph)angiogenesis-related within-group differences in the non-expander patients. Given the crucial role of angiogenesis and lymphangiogenesis, these results warrant a further examination of the biological and clinical relevance of this intra-group heterogeneity.

### MultiNicheNet infers tumor-specific cell-cell interactions that are supported by spatial co-localization analysis

For the following application, we looked for studies that generated multi-sample scRNA-seq and spatial transcriptomics data of the same samples. Both data modalities would enable us to verify whether highly prioritized interactions show spatial co-localization. The study conducted by Ji et al. fulfills this criterion^24^. Ji et al. performed scRNA-seq of tumor and healthy skin tissue of patients with cutaneous squamous cell carcinoma (cSCC) and spatial transcriptomics of tumor tissues from a subset of these patients^24^. We performed MultiNicheNet to unravel differences in cell-cell signaling between tumor and healthy skin tissue (while considering the paired nature of the data) (**Figure 4a, Supplementary Table 4a-b**, and **Supplementary Figure 9**).

**Figure 4.**
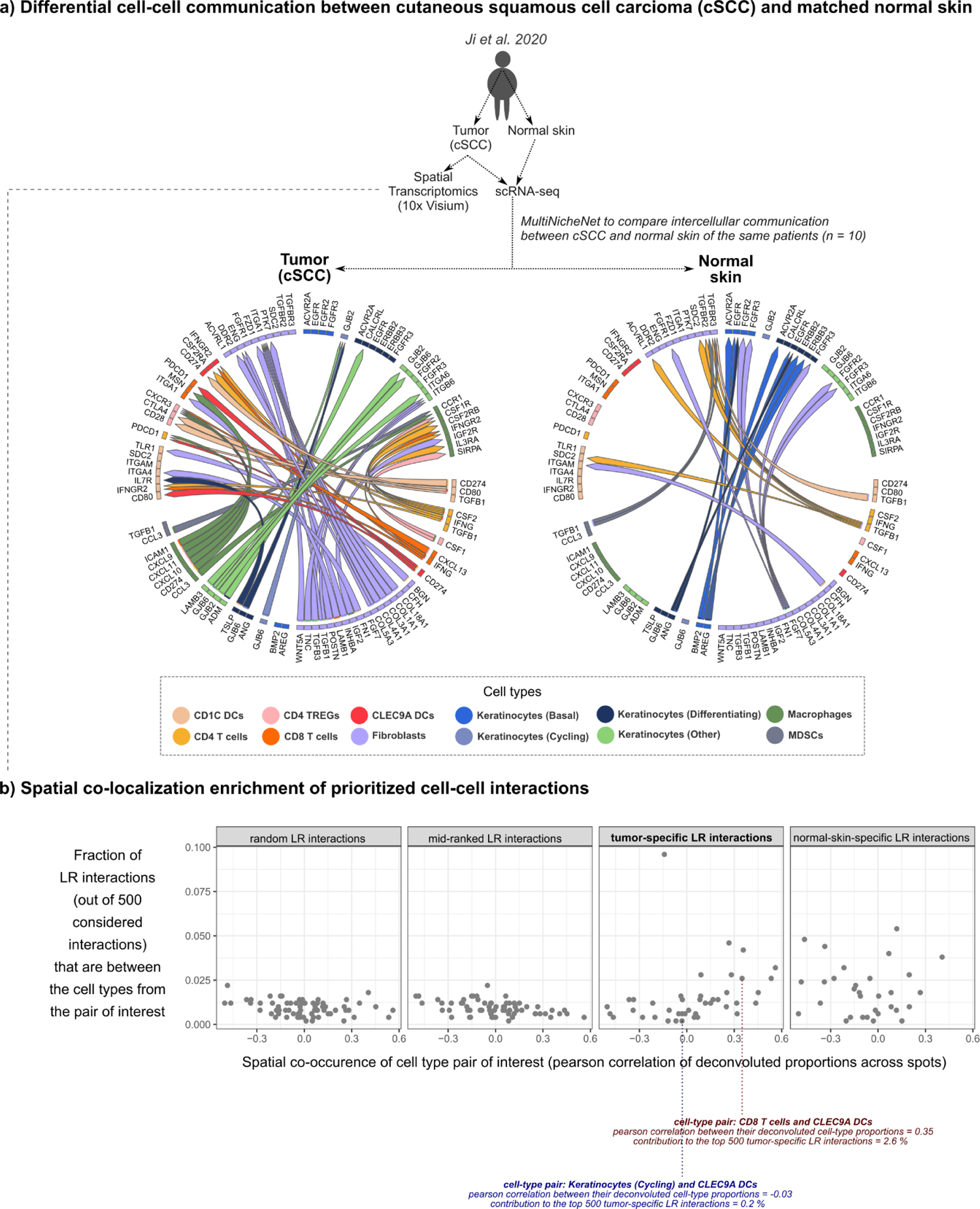
MultiNicheNet prioritizes differential cell-cell communication patterns between cutaneous squamous cell carcinoma (cSCC) and matched healthy skin from the same patient. a) MultiNicheNet’s top 75 differential ligand-receptor pairs between tumor and healthy skin are depicted in chord diagrams, divided per tissue type. The arrowhead indicates the direction from sender to receiver cell type, and the color of the arrow indicates the sender cell type that expresses the ligand. b) Spatial co-localization analysis of MultiNicheNet-prioritized cell-cell communication patterns. For the top 500 tumor-specific ligand-receptor (“LR”) interactions, we retrieved the cell types involving these interactions. Next, we determined the fraction of all interactions occurring between each combination of sender-receiver cell type pairs. This was also done for normal skin-specific interactions, a set of random interactions, and a set of interactions in the middle of the ranking. Spatial localization of cell types was determined by applying RCTD for deconvolution of the 10x Visium spatial transcriptomics data of tumor tissue (from one patient). Each dot represents one cell-type pair.

The tumor-specific interactions can mainly be divided into two parts: interactions among the different immune cell types and interactions among the fibroblasts and tumor cells (keratinocytes)(**Figure 4a, Supplementary Table 4a**, and **Supplementary Figure 9**). As expected, we retrieve the IFNG - PDL1 axis between T cells and myeloid cells (DCs and macrophages)^75^. Noteworthy, MultiNicheNet predicts several interactions between fibroblasts themselves, such as WNT, TGF-beta, inhibin, and collagen signaling. Fibroblasts also seem to interact with the tumor cells by producing TNC, TGFB3, and TGFB1 that can bind putative receptors on tumor-specific keratinocytes. These interactions were also described by Ji et al. in the original publication^24^. Some of the rare interactions between fibroblasts and immune cells are FN1-ITGA4 with DCs and IGF2-IGF2R with macrophages. Literature suggests that FN1 can improve antigen presentation in DCs and that IGF2 can modulate macrophage function^76^.

The intercellular regulatory network also shows this split between the interactions among immune cells versus interactions among fibroblasts and keratinocytes (**Supplementary Figure 10**). Nevertheless, some interactions seem to connect both systems, such as the IGF2-IGF2R interaction between fibroblasts and macrophages and the TSLP-IL7R interaction between keratinocytes and myeloid cells (DCs and macrophages). IGF2 is a known modulator of macrophage function, and TSLP is an epithelial cell-derived cytokine that is known to activate DCs^76, 77^. The observation of a myriad of TSLP-specific target genes in DCs provides further support for the potentially pivotal role of TSLP in modulating DCs in the cSCC tumor microenvironment (**Supplementary Figure 11** and **Supplementary Table 4c**). Moreover, previous research suggests that TSLP-activated DCs are linked to T cell hyporesponsiveness due to PD-L1 upregulation^78^. This confirms what we observe in the intercellular regulatory network, where TSLP appears at the top of the hierarchy of the predicted signaling cascades (**Supplementary Figure 10**).

Whereas the authors of the original study used spatial transcriptomics data to refine their cell-cell communication analysis^24^, we decided to use this data to verify spatial co-localization of MultiNicheNet-prioritized interactions. We would expect that the majority of cell-type pairs with the most differential interactions would spatially co-localize. This hypothesis was also the basis of one of the benchmark procedures described in a recent comparison study of cell-cell communication inference methods^10^. As expected, tumor-specific ligand-receptor pairs were mainly inferred between cell types that show co-localization in 10x Visium spatial transcriptomics data of tumor tissue (**Figure 4b**, **Supplementary Figure 12**, and **Supplementary Table 4d**). On the contrary, a random set of ligand-receptor pairs, pairs in the middle of the prioritization ranking, and healthy skin-specific pairs did not show co-localization enrichment in tumor spatial transcriptomics data (**Figure 4b**).

To summarize, the results of this case study indicate that a spatially-agnostic MultiNicheNet analysis can prioritize biologically relevant interactions between cell-type pairs that are spatially co-localized.

### MultiNicheNet compares cell-cell communication between PBMCs of adult COVID-19, MIS-C, and healthy siblings and prioritizes differential signals supported by serum protein analyses

The next case study will demonstrate that MultiNicheNet can compare communication patterns between three conditions. We applied MultiNicheNet to scRNA-seq data from Hoste et al. who performed single-cell sequencing to shed light on the disease pathophysiology of “multisystem inflammatory syndrome in children” (MIS-C)^18^. MIS-C is a novel immune dysregulation syndrome that can arise in rare instances a few weeks after pediatric SARS-CoV-2 infection. Specifically, Hoste et al. performed single-cell sequencing on peripheral blood mononuclear cells (PBMCs) from patients with MIS-C, healthy siblings, and adults with severe Coronavirus disease 2019 (COVID-19)^18^.

Differential cell-cell communication analysis with MultiNicheNet reveals several MIS-C-specific communication signals, such as IFNG (from T cells to monocytes), the CCR5-binding chemokines CCL3 and CCL4 (from NK and T cells to proliferating T cells) and RETN and the alarmins S100A8 and S100A9 produced by CD14+ monocytes (**Supplementary Figures 13-14** and **Supplementary Table 5**). Five independent studies describing different patient cohorts indicate that these signals are more present at the protein level in serum of MIS-C patients compared to healthy controls and/or COVID-19 patients^18, 28, 79–81^. According to both Sacco et al. and Hoste et al., the serum of MIS-C patients contains higher levels of the protein IFNG and its induced chemokines CXCL9 and CXCL10 compared to healthy controls and COVID-19 patients (respectively pediatric and adult)^18, 80^. Higher serum protein levels of IFNG were also reported by Carter et al.^79^, higher levels of CXCL10 by Ramaswamy et al^28^, and higher levels of both CXCL9 and CXCL10 by Diorio et al^81^. Moreover, both Sacco et al. and Diorio et al. detected higher protein levels of the chemokines CCL3 and CCL4 in the serum of MIS-C patients compared to healthy controls^80, 81^. For CCL3, protein serum levels were also higher in MIS-C compared to COVID-19 patients according to Sacco et al^80^. In addition, Diorio et al. reported higher serum protein levels of RETN in MIS-C patients versus healthy controls^81^. Furthermore, Ramaswamy et al. also confirmed the stronger expression of the proteins S100A8 and S100A9 by classical CD14+ monocytes in MIS-C through flow cytometry, just like other signs of myeloid dysfunction such as lower expression levels of CD86 and HLA-DR molecules^28^. Consistent with this finding, MultiNicheNet retrieves CD86 and HLA-DR molecules as members of ligand-receptor pairs specific for healthy siblings (**Supplementary Figure 13** and **Supplementary Table 5b**).

We want to emphasize that higher protein levels of the members of the inferred ligand-receptor interactions do not necessarily imply that the inferred interactions are occurring. This analysis can thus not serve as a direct validation of MultiNicheNet’s predictions. However, interactions between members that are expressed at the protein level are more likely to take place than interactions consisting of members that are not expressed at the protein level. As described above, MultiNicheNet includes predicted ligand activities in the prioritization because ligand activity might provide additional support that a ligand is also expressed at the protein level. The results of this MIS-C case study illustrate this idea: top-predicted MIS-C specific ligands with differential expression at the RNA level and high ligand activities are also differentially expressed at the protein level (**Supplementary Figure 14**). Future use-cases that also include ligands with differential RNA expression but with low ligand activities could further emphasize the relative importance of ligand activities in predicting protein expression of the ligand.

An additional benefit of the ligand activity analysis is that it allows exploring ligand-induced expression signatures for ligands that do not show expression in the data. These latter ligands might only be expressed shortly before the specific data was captured, or they might be produced by non-profiled cell types. We want to emphasize that ligand activities are always calculated for all ligands in the database during a MultiNicheNet run, and that users can thus easily access these results afterward. Ligand activity analysis for all the cell types in the scRNA-seq data of Hoste et al. points to the presence of a clearer type I interferon signature in adult COVID-19 patients compared to MIS-C patients (**Supplementary Figure 15**). Supporting this result, Sacco et al. and Hoste et al. detected higher serum levels of IFNA2 (one of the main type I interferons) in COVID-19 patients compared to MIS-C patients^18, 80^. Type I interferon signaling was not inferred as a COVID-19-specific interaction by the regular MultiNicheNet analysis because no type I interferon ligand-receptor pairs were expressed. This is probably because the primary producers of type I interferons, plasmacytoid DCs^82^, were not included in the analysis.

Furthermore, the ligand expression-agnostic activity analysis points to a possible role of a few other noteworthy ligands. These ligands were not retrieved by the regular MultiNicheNet analysis, but they exhibit a high predicted upregulatory activity in MIS-C patients and are MIS-C-specific proteins according to the serum proteomics data of Diorio et al^81^. One of these ligands is IL10, the ligand with the highest activity score for CD14+ monocytes and one of the only differentially expressed proteins between MIS-C patients and patients with severe COVID-19^81^. Another ligand is PLA2G2A, the ligand with the highest activity score for TRDV2+ gd activated T cells and, strikingly, the most strongly differentially expressed protein between MIS-C patients and healthy controls^81^.

In brief, MultiNicheNet uncovered several cellular communication signals that might contribute to MIS-C pathophysiology. Several independent studies reported higher protein levels of these signals in the serum of MIS-C patients.

### MultiNicheNet corrects for batch effects in integrated lung atlas data to reveal dysregulated intercellular communication in idiopathic pulmonary fibrosis

In the previous applications, we showcased MultiNicheNet’s potential on multi-sample data generated by one lab. In the last case study, we will apply MultiNicheNet on integrated data from four studies comparing healthy lungs to lungs from patients with idiopathic pulmonary fibrosis (IPF). This analysis exemplifies that MultiNicheNet can be used to compare cell-cell communication between health and disease from integrated atlas data.

Starting from the Azimuth-integrated atlas of ten scRNA-seq datasets of healthy and diseased human lungs, we searched for datasets that had profiled lungs from both healthy patients and patients with IPF^42^. Four datasets fulfilled this criterion^46–49^. Together with the provided Azimuth-harmonized cell type annotations, we used the raw counts of these datasets and ran MultiNicheNet while correcting for the source dataset (**Methods**). MultiNicheNet retrieved several IPF-specific communication patterns representing biological processes known to play a role in IPF (**Figure 5a**, **Supplementary Figure 16,** and **Supplementary Table 6**). These include TGF-β signaling, aberrant extracellular matrix (ECM) deposition and remodeling, and recruitment of fibrogenic myeloid cells^83–85^. Interactions involving TGFB1 and TGFB2 represent the first process (**Figure 5a** and **Supplementary Table 6a**)^83^. The predicted interactions between collagens, integrins, and the ECM remodeling mediators metalloproteinases (MMPs) and their inhibitors (TIMPs) illustrate the second process (**Figure 5a** and **Supplementary Table 6a**)^84^. The chemoattractive interaction CXCL12-CXCR4 between fibroblasts and AREG-and-TGFB1-producing monocytes is indicative of the third process (**Figure 5a** and **Supplementary Table 6a**)^85^. In line with this, is the finding of the IPF-specific CSF1-CSF1R interactions towards macrophages: monocyte-derived recruited alveolar macrophages are known to depend on CSF1 rather than CSF2 (which is the crucial growth factor for alveolar macrophages in homeostasis) (**Supplementary Table 6a**)^85, 86^.

**Figure 5.**
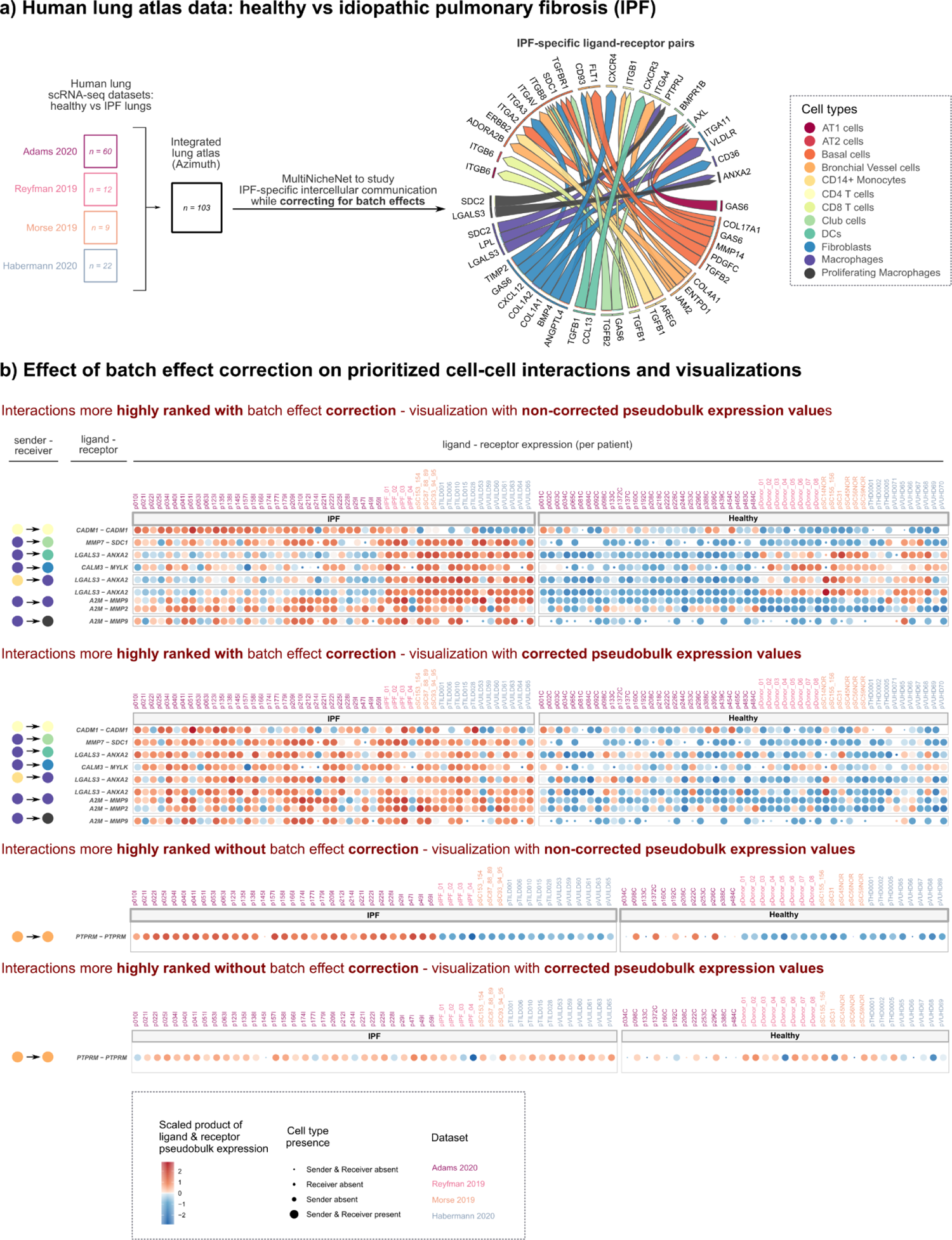
MultiNicheNet corrects for batch effects in integrated scRNA-seq atlas data to compare cell-cell communication patterns between lungs from patients with idiopathic pulmonary fibrosis (IPF) and healthy patients. **a)** MultiNicheNet was applied to an integrated scRNA-seq lung atlas built from four source datasets that profiled both healthy and IPF lungs, while considering each source dataset as a separate batch. IPF’s most specific ligand-receptor pairs are depicted in chord diagrams. The arrowhead indicates the direction from sender to receiver cell type, and the color of the arrow indicates the sender cell type that expresses the ligand. **b)** For a subset of interactions for which the prioritization score strongly depended on batch correction, we compared corrected and non-corrected ligand-receptor pseudobulk expression values. The size of the dots indicates whether a sample had enough cells (>= 5) for a specific cell type to be considered for DE analysis.

Moreover, MultiNicheNet prioritized some interactions of which the role in IPF has not been well described. One such interaction is between CXCR3 from CD8 T cells and the chemokine CCL13 produced by DCs, macrophages, and fibroblasts (**Figure 5a** and **Supplementary Table 6a**). Another interaction concerns the receptor AXL on DCs and the ligand GAS6 produced by several cell types such as fibroblasts, basal cells, club cells, and type 1 alveolar epithelial cells (AT1) (**Figure 5a** and **Supplementary Table 6a**). GAS6-AXL is involved in the phagocytosis of apoptotic cells and debris^87^, a crucial process in lung fibrosis, and is also linked to the survival and migration of DCs^88^. Interestingly, both interactions are only prioritized when performing batch correction because they are not part of the top 1000 interactions when executing the analysis without correction (**Supplementary Table 6a,c**). In the following paragraphs, we will elaborate on the influence of the batch effect correction on MultiNicheNet’s prioritizations.

If strong batch effects are present, we expect substantial differences in the MultiNicheNet output since correction will affect most prioritization criteria in MultiNicheNet: differential expression of the ligand and receptor; ligand activity which is calculated based on the set of DE genes in the receiver cell type; and the cell-type and condition-specific expression of the ligand and receptor, which is calculated from the normalized pseudobulk values (**Methods**). Improper handling of batches will thus likely lead to improper downstream prioritization. To exemplify the effect that the correction can have on the DE analysis output, we show the genes that are only DE in macrophages after correction in **Supplementary Figure 17**. This geneset contains several genes linked to essential macrophage biological processes such as proliferation (*CCND1*, *WEE1*, *POLD2*, *RGCC*, *EYA2*), motility (*MYL9*, *ASAP1*, *PLXNA1*, *PLXNC1*, *PLXNB1*, *TPM1*, *TPM4*, *KIF13B*, *EZR*), metabolism (*FABP5*, *TREM2*, *LEPR*, *LPL*, *FAM3C*, *BCAT1*, *SCARB1*), ECM and ECM remodeling (PAPLN, *VIM*, *TIMP3*, *DPP4*), costimulation (*CD40*), and phagocytosis (*MERTK*, *TAGLN2*). Consequently, predicting ligand activities without including these genes could lead to missing crucial biological signals.

Whereas the previous analysis focused on DE genes in one cell type, we also looked at which ligand-receptor pairs would be more or less clearly DE with or without correcting batch effects (**Figure 5b**). One of these ligand-receptor pairs is LGALS3-ANXA2, which was found to be upregulated in IPF between myeloid cells (macrophages, monocytes, and DCs) only after batch correction (**Figure 5b**). LGALS3 has been shown to be involved in the pathogenesis of IPF by activating macrophages and fibroblasts^89^, and to improve the ability of macrophages to phagocytose apoptotic cells in chronic obstructive pulmonary disease (COPD)^90^. The specific interaction LGALS3-ANXA2 has not been linked to IPF before but was found to be anti-apoptotic in breast cancer^91^. A second interaction specifically DE after batch correction is the interaction between the matrix metalloproteinase MMP7 from macrophages and the syndecan SDC1 from club cells (**Figure 5b**). This interaction suggests possible shedding of syndecans by MMPs in IPF. Syndecan shedding is a process in which proteases like MMPs cleave the ectodomains of the syndecans from the cell membrane. These cleaved syndecan ectodomains can then bind and modulate the activity of chemokines, growth factors, and cell surface receptors, consequently influencing processes like fibrosis^92^. Moreover, expression levels of MMP7 itself are associated with IPF disease progression^93^. The last correction-specific IPF ligand-receptor pairs we will discuss are A2M-MMP2 and A2M-MMP9 between macrophages (**Figure 5b**). A2M is a general protease inhibitor and might thus be involved in inhibiting the ECM-remodeling activity of MMPs^94^.

One of the main interactions more strongly DE without batch correction is the predicted PTPRM-PTPRM interaction between bronchial vessel cells (**Figure 5b**). This interaction illustrates one of the scenarios for which batch correction is necessary. The *PTPRM* gene is more strongly expressed in one of the datasets compared to the others, and this dataset has more samples with a sufficient number of bronchial vessel cells in the IPF group (**Figure 5b**). As a consequence, the *PTPRM* gene is considered to be IPF-specific.

To conclude, MultiNicheNet’s ability for batch correction enables performing differential cell-cell communication analysis from integrated atlas data. Although most of the prioritized interactions were also retrieved without batch correction (**Supplementary Table 6**), some correction-specific DE genes and interactions are associated with critical processes in IPF pathogenesis. This illustrates that improperly handling batches might conceal relevant biological signal in the data.

## Discussion

In this paper, we presented two main contributions to the field of cell-cell communication modeling, namely NicheNet-v2, an updated NicheNet prior knowledge model of ligand-receptor and ligand-target links, and MultiNicheNet, a novel framework for differential cell-cell communication analysis.

The improvements upon NicheNet-v1’s prior knowledge include the use of the up-to-date and comprehensive Omnipath-derived ligand-receptor network and the addition of several new signaling and gene regulation data sources. As a consequence, ligand-target predictions improved substantially according to our benchmark. The primary improvement of the ligand-target model is the inclusion of experimentally-determined ligand-target links inferred from ligand treatment datasets. For the 119 ligands for which these datasets were available, top-predicted target genes are now supported by both network-based signal propagation prediction and *in vitro* experiments. Enrichment of several of these target genes in a receiver cell may thus strongly point to the upstream activity of the particular ligand. To conclude, we expect that using NicheNet-v2 will improve the ligand-receptor prioritization and ligand-target predictions for (Multi)NicheNet analyses. Moreover, other cell-cell communication tools that use the NicheNet ligand-target model as part of their methodology (e.g., Scriabin^19^, LRLoop^95^, CINS^96^) or as part of their downstream interpretation (scITD^44^) will benefit as well. Nevertheless, we still see many opportunities for future work regarding the NicheNet prior model. One future improvement could be considering the multi-subunit architecture of receptors (similarly to, for example, CellphoneDB^4^ and CellChat^7^). Another potential improvement could be including non-protein ligands in the model, as done in, e.g., MEBOCOST^97^ and CellphoneDB-v4^98^.

As the second main contribution of this paper, we presented MultiNicheNet, a novel algorithm for differential cell-cell communication inference from multi-sample multi-condition scRNA-seq data. In four case studies, we demonstrated MultiNicheNet’s power in addressing different cell-cell communication questions in several biological systems. We showcased that MultiNicheNet can retrieve expected and well-studied biological patterns but that it also can elucidate sensible yet undiscovered patterns. From an algorithmic standpoint, MultiNicheNet is a flexible multi-criteria prioritization framework based on a pseudobulk aggregation DE approach to analyze multi-sample datasets in a statistically solid way^22^. MultiNicheNet has several benefits compared to the most widely used cell-cell communication tools when applied for differential cell-cell communication inference from multi-sample data. First, MultiNicheNet considers differential and cell-type-specific expression together with downstream target gene enrichment to prioritize ligand-receptor pairs. Second, it does so while taking into account inter-sample heterogeneity. Moreover, MultiNicheNet can address complex experimental designs, which enables tackling exciting non-trivial questions like investigating differences in intercellular dynamics between health and disease states. Furthermore, MultiNicheNet can correct for covariates and batch effects, such as data source effects in integrated atlas data. For this reason, we anticipate that MultiNicheNet will be a valuable tool for the Human Cell Atlas community to exploit the wealth of the collected data to elucidate the role of cell-cell communication in disease pathophysiology.

Additionally, MultiNicheNet offers a variety of easily interpretable visualizations to provide users with further insights into the data and predictions. We think it is of paramount importance for a hypothesis-generating tool that users can explore why certain predictions are made before deciding on which communication patterns to validate experimentally. As illustrated by the analysis of group differences in anti-PD1 therapy response, some visualizations allow for exploring inter-sample heterogeneity, potentially revealing hidden patterns of within-group differential cell-cell communication. Another novel type of downstream analysis and visualization is the inference of intercellular regulatory networks. These networks, which link ligands to ligand/receptor-encoding target genes, can only be constructed because MultiNicheNet links differential ligand-receptor pairs to their predicted target genes. By revealing how cell types might influence each other and how this differs between conditions, these networks can help investigate diseases from a tissue-centric perspective instead of a classic cell-type perspective. Although the case studies indicated that these networks could report expected links, the outcome from the proposed inference approach should be interpreted cautiously. Ligand-target links in this network are predicted based on correlation in expression across patients and prior knowledge support of ligand-to-target signaling. Yet, correlation does not necessarily imply causation, even though prior knowledge may suggest a potential causal link. Including context-specific perturbation data and spatiotemporal information would be necessary to refine the links in this network. Nevertheless, we still include this analysis type in the software framework to point users to potential intercellular crosstalk and help them further prioritize signals. We think this concept of intercellular regulatory networks is an interesting idea that fits within the notion of multicellular programs as proposed by others^43, 44^. Extending these approaches is an exciting avenue for further research to advance our ability to unravel tissue circuitry in health and disease.

In this paragraph, we will discuss the limitations of MultiNicheNet. Because MultiNicheNet is based on DE, cell types must be present in all compared conditions. This limits the analysis of condition-specific cell types. In some cases, it is possible to circumvent this issue by using a less fine-grained level of cell type annotation. Other limitations of MultiNicheNet are similar to the limitations of the multi-sample scRNA-seq DE methods that are at the basis of it. One disadvantage of the currently implemented method is that the pseudobulk aggregation step reduces the original single-cell resolved data to one point estimate per sample-cell-type combination. The next disadvantage is that the DE method can be underpowered when applied to datasets with a few samples. This is less of a problem for MultiNicheNet’s prioritization of DE ligand-receptor pairs because we use the DE p-values and log fold change (logFC) values to rank the pairs and not for hard thresholding. But, this issue might be more problematic for defining the entire set of DE genes in the receiver cell type, a necessary step to calculate ligand activities. Using the multiple-testing corrected p-value for filtering can result in only a few DE genes when there is a low number of samples. In those cases, we suggest that users consider the biological effect size and filter on the logFC value. Another disadvantage is that the pseudobulk approach requires a sufficient number of cells for accurate DE analysis. This approach is thus less suited to handle rare cell types because samples are omitted from the DE analysis for a certain cell type if they have fewer cells than a pre-defined cutoff (by default 10 in MultiNicheNet, but user-adaptable). When less than two samples are then left in one condition, DE analysis cannot be performed, and this cell type is excluded from the MultiNicheNet analysis. This issue also occurs in the case of batch effect correction, where at least one sample per condition-batch combination should be present. If one combination does not comply with this rule, the cell type is excluded as well. This was the case for myofibroblasts in the IPF case study, which is suboptimal, given their important role in IPF pathophysiology^99^. Therefore, using a coarser level in the cell type annotation hierarchy is often necessary. For example, in the IPF case study, we used the provided cell type annotation that considered non-proliferating lung macrophages as one cell type. However, this is a heterogeneous population consisting of alveolar macrophages, interstitial macrophages, and recruited monocyte-derived macrophages^85^. Potential subpopulation-specific DE patterns can then be concealed. Although these issues are significant, the currently implemented DE approach is still considered state-of-the-art in the multi-sample DE analysis field^21, 22, 50^. Nevertheless, we anticipate future developments in the DE analysis field, and we will include potentially more appropriate DE methods when available.

Whereas we focused on the limitations of the MultiNicheNet algorithm in the previous paragraph, we will now discuss the limitations of this study in general. The major limitation is that we did not perform a comprehensive quantitative assessment of MultiNicheNet’s performance. However, this is a common limitation for the entire computational cell-cell communication research field because of the lack of an extensive ground truth of essential communication patterns in various *in vivo* biological systems. That performing a conclusive benchmark of cell-cell communication tools is very challenging has been noted before by other researchers^2,3,10^. To partially circumvent this limitation, we employed a strategy consisting of three parts:

1. Using state-of-the-art quantitatively benchmarked approaches for submodules of MultiNicheNet, or performing such a benchmark ourselves;
2. Assessing whether alternative data modalities are in line with MultiNicheNet’s predictions in the case studies;
3. Performing a *qualitative* comparison of MultiNicheNet against other tools.

For the first part, we benchmarked the prior knowledge model of NicheNet-v2 to assess target gene prediction and ligand activity prediction performance *in vitro*. Both prediction tasks are crucial elements of the MultiNicheNet framework. In addition, we chose the pseudobulk-aggregation-edgeR approach for DE analysis based on recently published benchmarks^21, 22, 50^. Noteworthy, because MultiNicheNet is a flexible and modular framework, it will be easy to plug in potential future improvements of both submodules. For the second part, we searched for case study datasets that would enable us to check the accordance of MultiNicheNet’s predictions with other data modalities. In the cSCC case study, we used spatial transcriptomics data for verifying cell-cell interactions in a similar way as Dimitrov et al. described in their comparative study^10^. According to this spatial co-localization analysis, MultiNicheNet predicted tumor-specific cell-cell interactions that were indeed spatially co-localized. In the MIS-C case study, we found differential protein expression in the serum of patients for almost all top-ranked interactions. These additional data modalities may thus suggest that MultiNicheNet captured biological signal in the specific case studies. But, we want to emphasize that both data modalities do not necessarily provide strong evidence that the top-predicted interactions are occurring, nor that they are pivotal for the system under study. Because of these limitations and the limited number of datasets, we do not think that these data modalities can currently be used to quantitatively compare the performance of different tools in a conclusive way. Therefore, we compared several tools in a qualitative way (**Supplementary Note 4**), highlighting their primary use cases, underlying assumptions, potential pitfalls, and unique benefits. We think that several of these tools are complementary for the study of differential cell-cell communication patterns across patient groups, and that using them in conjunction might be a powerful analysis strategy. For example, multi-sample-specialized tools (DIALOGUE^43^, scITD^44^, MultiNicheNet, and Tensor-cell2cell^45^) shed light on inter-sample heterogeneity at the cost of potentially missing relevant patterns for rare cell types. In contrast, the opposite is true for tools that perform cell-cell communication analysis at single-cell resolution (Scriabin^19^ and NICHES^8^) (**Supplementary Note 4**). Compared to the other methods, MultiNicheNet stands out in its ability to combine ligand-receptor and ligand-target prioritization with the possibility of elegantly addressing batch effects and complex multifactorial experimental designs.

To conclude, we are convinced that MultiNicheNet is a necessary addition to the ecosystem of cell-cell communication tools. Given the anticipated increase in multi-sample datasets and atlas-generation efforts, we envision that MultiNicheNet will be a valuable tool to elucidate key cell-cell communication processes in health and disease states.

## Methods

### Construction of the NicheNet-v2 model

NicheNet uses a model of potential ligand-target links that are predicted based on data integration of ligand-receptor, signaling, and gene regulatory networks. To build an updated version of the NicheNet ligand-target prior model (NicheNet-v2), we used the same procedure as described in the original NicheNet publication^11^. However, we substantially updated the underlying networks.

One of the most important updates concerns the ligand-receptor network in NicheNet. For NicheNet-v2, we replaced the original NicheNet-v1 ligand-receptor network with a novel ligand-receptor network that comprises mainly ligand-receptor interactions documented in the comprehensive Omnipath intercellular communication database^52^. The Omnipath ligand-receptor network was processed to ensure high-quality ligand-receptor interactions with the correct directionality of the ligand-receptor links. As a processing strategy, we first used the annotations in the intercellular communication database to get confident annotations of ligands (“transmitters”) and receptors (“receivers”). In the next step, we queried the Omnipath interaction network for interactions between these confident transmitters and receivers. In addition to Omnipath, we also incorporated extra ligand-receptor interactions from Verschueren et al^53^. Finally, we also added interactions not documented in Omnipath but part of the curated databases at the basis of NicheNet’s original ligand-receptor network.

Regarding the signaling network, interactions from Omnipath^100^ and Pathwaycommons^101^ (version 12) were updated, and all other data sources were kept as they were. We also included additional interactions from two novel databases: HuRi^102^ and HumanNet^103^.

Finally, the gene regulatory network was updated and expanded with novel data sources. PathwayCommons^101^ and ReMap^104^ were updated (to respectively version 12 and 2022), and MOTIFMAP^105^ was removed. Novel regulator-target interactions were added from KnockTF^54^ (a database of gene expression signatures after regulator perturbation) and Dorothea^106^ (a comprehensive database of regulator-target interactions). Moreover, we also extended the gene regulatory network by adding direct ligand-target links. We used co-expression-based links between ligands and targets based on the HumanNet^103^ co-expression resource as the first source of direct ligand-target links. As the second and most relevant source, we added *in vitro* experimentally-determined links between ligands and their target genes. These links were gathered from “ligand treatment” datasets, which are datasets from experiments where the transcriptome of cells is analyzed before and after stimulation with a specific ligand. Differentially expressed genes after ligand treatment can then be considered target genes of the particular ligand. More specifically, these links were added from the original NicheNet^11^ publication and CytoSig^55^, a database documenting thousands of transcriptome profiles for human ligand responses. Whereas this type of link was only used for model validation and optimization in the original NicheNet study^11^, we now also included these highly-relevant links to the prior knowledge model of NicheNet-v2. Consequently, we had to adapt the evaluation and optimization procedure to ensure an appropriate validation (see section “Evaluation of NicheNet-v2 model”).

The specific processing steps of each new data source are described in more detail in **Supplementary Note 1**. Information regarding each data source and results of network analyses are presented in **Supplementary Note 1** and **Supplementary Table 1**.

### Evaluation of the NicheNet-v2 model

#### Evaluation of the performance of NicheNet-v2 in target gene and ligand activity prediction

Target gene and ligand activity prediction performance was evaluated on ligand treatment datasets as described in the original NicheNet publication. A difference is that we now have two gold standard (GS) datasets: 1) the same 111 ligand treatment datasets from the original NicheNet publication^11^ and 2) 52 datasets from CytoSig. To define the CytoSig GS, we started from the provided “Cytokine signatures” file, which provides median logFC values for each gene after ligand treatment, across a subset of datasets in which the *in vitro* transcriptional response was considered to be relevant for human physiological responses^55^. To handle the different ranges of logFC values across the different ligand signatures, we z-score normalized the logFC values per signature and considered genes a target of a ligand when |z-score normalized logFC| >= 2.5. These genes were thus considered to be a “true positive” target of a ligand, and other genes were not considered as targets (“true negative”). **Supplementary Table 2a** provides an overview of all the ligand treatment datasets.

To ensure a correct evaluation procedure, we removed all NicheNet-ligand treatment-derived links from the model before using the NicheNet-ligand treatment-GS for evaluation. The same was done for the CytoSig-GS links. Because some ligand treatment datasets from the NicheNet publication are also part of CytoSig (this overlap was determined based on GEO accession numbers), we removed CytoSig’s ligand-target links from overlapping datasets before evaluating on NicheNet’s ligand treatment GS datasets.

As the original NicheNet publication described, ligand activities are predicted based on the most predictive “target-gene-prediction performance metric”^11^. In contrast to the original study, this metric was now the area under the precision-recall curve (AUPRC) instead of the Pearson correlation (for the NicheNet GS: average area under the receiver-operating-characteristic curve (AUROC) for ligand prediction was 0.884 when using the AUPRC-target-prediction versus 0.863 when using the pearson-target-prediction, the average AUPRC for ligand prediction was 0.458 versus 0.430; for the CytoSig GS: the average AUROC for ligand prediction was 0.860 versus 0.827, the average AUPRC was 0.318 vs 0.302).

To compare the unoptimized and optimized NicheNet-v2 models (**Supplementary Note 2**), we used the following parameters to create the unoptimized model: the ligand-signaling network hub correction factor and gene regulatory network hub correction factor 0; the cutoff on the Personalized PageRank vector 0.9; and damping factor 0.5. We defined the optimized model’s parameters via parameter optimization as described in the next section.

### Parameter optimization via NSGA-II

Because new data sources were added to the NicheNet-v2 model, the model hyperparameters and data source weights needed to be updated through parameter optimization. As described previously^11^, we performed multi-objective optimization to optimize both target gene and ligand activity prediction performance on all ligand treatment datasets (NicheNet and CytoSig). We opted for NSGA-II^107^ (*nsga2r* package), a genetic algorithm that is a state-of-the-art method that allows for multi-objective optimization of expensive black-box functions. We chose this over model-based optimization (the technique used in the original NicheNet publication) because of its shorter running time. However, as demonstrated in the NicheNet paper, both options rendered very similar results. Optimization was performed during 15 iterations with a population size of 360 and a tournament size of 2. Nsga2r’s default values were chosen for the other parameters of the optimization algorithm. As done previously, the optimization criteria were: AUROC for target gene prediction, AUPRC for target gene prediction, AUROC for ligand activity prediction, and AUPRC for ligand activity prediction. In contrast to the NicheNet-v1 optimization process, proposed parameter values for the data source weights were set to zero or one when they were respectively < 0.025 or > 0.975. The rationale behind this was to facilitate the removal of potential low-quality data sources instead of keeping them with very low weights. We did not perform this approach for ligand-receptor data sources to avoid some ligands being absent in case of data source weights of 0.

We conducted the following cross-validation-like strategy to avoid overfitting and over-optimistic performance estimation. NicheNet’s and CytoSig’s ligand treatment datasets were each randomly divided into five groups, and a fivefold cross-validation was performed (**Supplementary Table 2b**). Per fold, 4/5 of the NicheNet and 4/5 of the CytoSig ligand treatment datasets were used for calculating the optimization objectives during optimization (“training set”). The left-out 1/5 of NicheNet and CytoSig datasets (“test set”) were used for the final evaluation of the model (including the comparison to NicheNet-v1). During the training of one fold, we defined all the ligands with a corresponding GS training dataset from NicheNet in that fold. For these ligands, we then removed all NicheNet-ligand-treatment-derived ligand-target links from the gene regulatory network before model building. This strategy is necessary to avoid leakage from the model building to the model evaluation and optimization (because both the links in the network and GS come from the same datasets). Noteworthy, the gene regulatory network still contains the NicheNet-ligand-treatment-links of the ligands from the 1/5 of left-out datasets. Therefore, the training process can learn how these NicheNet-ligand-treatment-links help predict the independent GS links from CytoSig. The same was done for ligands and their links from the CytoSig ligand treatment datasets. At the end of the optimization process of each fold, we built a model with the median parameter values of the 25 parameter settings with the highest geometric average over the four objectives described above. We then applied this model to the corresponding left-out test GS datasets to calculate the target gene and ligand activity prediction performances used to evaluate the model (as described in “*Evaluation of the performance of NicheNet-v2 in target gene and ligand activity prediction”*). This procedure was repeated for every fold, and these performance results were then aggregated over all folds to obtain the results shown in **Supplementary Figure 1** and **Supplementary Table 2**.

Finally, we defined the hyperparameter values and data source weights for the NicheNet-v2’s final ligand-target matrix that is used for application purposes. These parameter values were calculated as the median of the final parameter settings of all folds (**Supplementary Table 1e-f**). Noteworthy, for the final application model, all CytoSig and NicheNet ligand treatment target links are kept in the gene regulatory network before model construction.

As a result of this approach, the parameter optimization procedure optimizes for both the ligands with ligand-treatment-derived links in the gene regulatory network and the ligands without (these are the majority of ligands). However, model performance is still assessed unbiasedly without data leakage.

### MultiNicheNet algorithm description

MultiNicheNet aims to study differences in intercellular communication between conditions of interest from multi-sample multi-condition single-cell transcriptomics data. MultiNicheNet prioritizes ligand-receptor interactions based on the following criteria: differential expression of the ligand and its receptor(s); differential ligand activity in the receiver cell type; cell-type and condition-specific expression of the ligand and its receptor(s); the fraction of samples in the condition of interest with sufficient expression of the ligand-receptor interaction. The steps to calculate each of these prioritization factors are described in the next paragraphs.

#### Required user input for a MultiNicheNet analysis

The following elements are required as input for MultiNicheNet:

1. a scRNAseq raw count matrix (dimensions *G* x *C*, with *G* the number of genes and *C* the number of cells)
2. a metadata matrix providing the sample, group, and cell-type label per cell (and batch and covariate information if applicable)
3. a list of contrasts of interest to indicate the cell-cell communication comparisons that should be assessed. Contrast can range from simple (e.g. *condition A – condition B*) to more complex (e.g. *condition A – [condition B + condition C]; [condition A – condition B] – [condition C – condition D]*).
4. a list of the sender and receiver cell types of interest to indicate between which cell-type pairs cell-cell communication should be analyzed. This can be all cell-type pairs in the data but also specific sender-receiver cell-type combinations, e.g., to restrict the analysis to cell types located in the same tissue microenvironment.
5. a ligand-receptor network and corresponding ligand-target matrix (in this study, the NicheNet-v2 version was used for both).

### Step 1: Performing DE analysis after pseudobulk aggregation

In the first step, MultiNicheNet starts from the provided matrix of raw counts to generate a pseudobulk count matrix for each cell type. Pseudobulk aggregation is performed by summing all the counts of that cell type per sample. This results in *k* matrices with dimensions *G* x *S* with *S* the number of samples and *k* the number of cell types.

Next, these pseudobulk matrices are used for DE analyses via edgeR^51^. We chose the options to sum the raw counts to generate pseudobulk matrices and EdgeR for DE analysis based on the benchmark of “differential state analysis approaches” performed by Crowell et al^22^. Differential testing is performed for each provided contrast. Batch and covariates variables can be added to the model matrix to perform batch/covariate-corrected DE analysis. DE analysis is only performed for cell types with at least a certain minimum number of cells (*min_cells*) in at least two samples per group/condition. The default value of *min_cells* is ten^22^. If a batch or categorical covariate variable is added to the model matrix, cell types should have more than *min_cells* in at least one sample per group-batch/covariate combination. In addition to filtering cell types, gene filtering is also performed. Similarly to the implementation of *muscat*, this is done by the function *filterByExpr* of the *edgeR* package^22, 51^. However, we changed the default filtering parameters because we found those to be rather stringent for pseudobulk expression data for most scRNA-seq multi-sample human cohort datasets: *min.count = 7* (instead of 10) and *large.n = 4* (instead of 10). The first adaptation reduces the minimum number of counts required for a gene to be considered as expressed in a sample. The second adaptation reduces the number of samples in which a gene should be expressed to be retained.

By default, edgeR first models the expression of each gene by fitting a quasi-likelihood negative binomial generalized linear model. Next, it will compute a test statistic for each gene for the model parameter of interest. The null distribution of these test statistics is theoretically expected to be an F-distribution. However, in large-scale inference settings, deviations from the theoretical null distribution are often observed. Efron^108^ has proposed four reasons for why the theoretical null distribution may fail: failed mathematical assumptions, correlation across features (here: genes), correlation across samples/subjects, and unobserved confounders. To avoid these issues, Efron^108^ proposed a strategy to empirically estimate the null distribution of the test statistics and use this to estimate empirical p-values. Therefore, we additionally implemented the strategy of estimating empirical p-values in case the theoretical null distribution would be invalid^109^. These empirical p-values can then be corrected for multiple testing with the method of Benjamini and Hochberg to control the False Discovery Rate (FDR)^110^. This strategy was implemented in the MultiNicheNet (“*multinichenetr”*) package in the same way as described in the satuRn paper and package^109^. As described in the *multinichenetr* package vignettes, we recommend that users check the p-value distributions for each cell type after DE analysis and assess whether these p-value distributions indicate a potential violation of the theoretical null. In case of violation (when the p-value distribution would not be uniform or uniform with a peak at p < 0.05), we recommend users to adapt the analysis settings (for example, add necessary covariates to the model) or continue the pipeline with the empirical p-values.

Finally, the DE logFC and p-value are stored for each gene for each cell-type-contrast combination. This enables defining all significantly DE genes in each receiver cell type and keeping track of the strength of differential expression of each ligand and receptor in all sender and receiver cell types. All this information will be used for prioritization as described in the corresponding paragraph.

### Step 2: Calculating normalized pseudobulk expression values

In the second step of the pipeline, we will calculate normalized pseudobulk expression values. These values are used later on for downstream visualizations and to define the cell-type- and condition-specific expression of each ligand and receptor (which is one criterion in the final prioritization). This step, and all the next steps, are only performed for the cell types that were retained after the cell type filtering step before the DE analysis.

MultiNicheNet performs pseudobulk count normalization for each cell type separately. First, library size normalization is performed by dividing the raw pseudobulk counts by the effective library size, in the same way as described in the edgeR tutorials^51^. Second, these normalized pseudobulk counts are multiplied by one million, followed by adding a pseudocount of one and a log2 transformation. Hereby, pseudocount addition will allow that a zero-count value as input gets transformed into a log-normalized value of 0, without substantially changing the log transformation of non-zero values.

When a user indicates that batch correction should be performed, the pseudobulk values will be normalized in the following way. First, the raw count matrix is batch-corrected by applying Combat-seq^111^. Combat-seq uses a negative binomial regression model that considers the input experimental design and returns a corrected count matrix in integers. We opted for Combat-seq correction of pseudobulk counts because batch correction is performed as in the edgeR DE analysis described in the previous step. The Combat-seq-corrected pseudobulk count matrix is then normalized and log2-transformed in the same way as described above. We want to emphasize that we only recommend performing this Combat-seq correction of pseudobulk counts only for batch effects and not for covariates like sex and age.

Finally, the log-normalized pseudobulk gene expression values are stored for each cell-type-sample combination. This information will be used for prioritization and visualization as described in the corresponding paragraphs.

### Step 3: NicheNet ligand activity analysis and ligand-target inference

In this step, the output of the DE analysis is used to calculate NicheNet ligand activities. For each receiver cell type and contrast of interest, a set of upregulated and downregulated genes is determined. Based on these genesets, “upregulatory” and “downregulatory” NicheNet ligand activities are calculated for all ligands in the database. The AUPRC in target gene prediction is used as the ligand activity metric, with all genes in the NicheNet ligand-target matrix as the background^11^. To determine the exact set of upregulated and downregulated genes per contrast, the default logFC thresholds are respectively 0.50 and −0.50 and the default p-value threshold is 0.05. Using the adjusted p-values is recommended in general, except when only a very few genes (< 10) are significantly upregulated or downregulated. This can occur when the data has a limited number of samples because pseudobulk DE methods can be too conservative^20^. It is recommended to check the number of genes in the final up -and downregulated genesets because NicheNet ligand activity prediction might be less accurate when the number of genes in the geneset is very small (< 20) or very high (> 2000).

Both upregulatory and downregulatory ligand activities are stored for each ligand-receiver-contrast combination. These activity values are then z-score normalized per contrast-receiver combination. We performed this normalization because our benchmark of ligand activity prediction indicates that the relative ranking of ligands per analysis is more informative than the absolute ligand activity value^11^.

Finally, target gene inference is performed per ligand-receiver-contrast combination in the same way as described in the original NicheNet publication^11^, by using the *get_weighted_ligand_target_links* function from the *nichenetr* package. This will only consider target gene predictions when a target gene belongs to the *top n* of targets of that ligand according to the ligand-target model of regulatory potential (default *top n = 250)*.

### Step 4: Prioritization of differential ligand-receptor pairs

In this step, prioritization of differential ligand-receptor pairs will be performed by considering the information calculated in the previous steps. In general, the scores from each criterion will be scaled between 0 and 1, followed by a weighted aggregation.

The first criterion is the level of differential expression of the ligand in the sender cell type. The logFC of each ligand in each sender is transformed into a value between 0 and 1 by calculating the rank over all ligand-sender-contrast combinations, and dividing this by the maximally possible rank. For example, the ligand with the highest logFC value out of 1000 ligand-sender-contrast combinations has a rank of 1000, which after division by 1000 gives a score of 1. The second most strongly upregulated ligand gets then a score of 0.999, etcetera. We preferred a rank-based scaling over a z-score or min-max scaling such that the final score reflects whether a ligand is DE or not, rather than the absolute strength of differential expression. Using a z-score or min-max scaling could lead to overemphasizing very strongly DE ligands. We wanted to avoid this because we consider the biological relevance of very strongly DE versus moderately DE (e.g., logFC of 4 versus 1.5) less relevant for differential ligand prioritization than the difference between moderately DE and not DE (e.g., logFC of 1.5 versus 0.1). In addition to the logFC-transformed score for ligand differential expression, MultiNicheNet also calculates a p-value-transformed score. To calculate this score, the negative log10 is calculated, followed by multiplication with the sign of the logFC value such that strongly upregulated ligands get the highest scores, and strongly downregulated ligands the lowest. Subsequently, rank-based scaling over all ligand-sender-contrast combinations is performed.

The second criterion is the level of differential expression of the receptor in the receiver cell type. Both a logFC- and p-value-based score are calculated, in the same way as described in the previous paragraph for ligands. The only difference is that rank-based scaling is now performed across all receptor-receiver-contrast combinations instead of ligand-sender-contrast.

The third criterion is the ligand activity of the ligand in a receiver cell type for a specific contrast. Min-max scaling of the z-score-normalized ligand activity values (with cutoffs on the bottom and upper 1% quantile to reduce outliers’ impact) is performed over all ligand-receiver-contrast combinations. In contrast to the DE-based scores, we used min-max quantile scaling to assign more weight to ligands with substantially higher ligand activity values. This scaling is performed for both the up-and downregulatory activities per contrast. As the final ligand activity criterion score, the maximum of the scaled up- or downregulatory activity is selected. In this way, we do not prioritize ligands that have both high up-and downregulatory activity over ligands with only one of the two.

We did this because ligands that only induce upregulation or downregulation of target genes are not necessarily less relevant than ligands that do both.

Whereas these three criteria are calculated per contrast, the following criteria will be calculated per condition. Therefore, MultiNicheNet users must connect each contrast to a specific group if they want to calculate the next criteria. For most questions, this is very straightforward. For example, for the contrast *“condition A – [condition B + condition C]”*, the condition of interest is “condition A”; for the contrast *“condition B – [condition A + condition C]”*, the condition of interest is “condition B”.

For the next criterion, the log-transformed normalized pseudobulk expression values of each ligand are first averaged per condition. Then, min-max scaling is performed for each ligand separately across all condition-sender combinations. As a result, this score reflects whether a ligand is expressed in both a condition-specific and sender-specific way. For receptors, a similar procedure is applied across receiver instead of sender cell types.

The last criterion represents the fraction of samples (per condition) with sufficient expression of both ligand and receptor. For each sample and ligand-receptor pair, it is assessed whether the ligand-receptor pair is sufficiently expressed by the corresponding sender-receiver cell types in that sample. By default, MultiNicheNet considers a ligand/receptor expressed when the encoding gene has a non-zero value in >= 5% of cells of the sender/receiver. Next, the fraction of samples with sufficient expression is determined for each condition.

After the calculation of all the aforementioned criteria, a weighted aggregation is performed to get a final prioritization score for each sender-receiver-ligand-receptor-contrast combination. The score after weighted aggregation is divided by the sum of the used weights to scale the final prioritization score in the range between 0 and 1. This score then reflects the most relevant differential ligand-receptor pairs, and a score of 1 would mean that a particular sender-receiver-ligand-receptor-contrast combination obtains the maximum score for each prioritization criterion. Noteworthy, all contrasts are considered together so that it is possible to assess which contrast has the most strongly differential interactions. The used weights can be adapted by the user according to insight into the specific dataset and personal preference to emphasize certain criteria more strongly. However, we recommend following parameters to balance the different criteria evenly.

- The weight used to multiply with the ligand-logFC score and the ligand-p-value score = 1
- The weight used to multiply with the receptor-logFC score and the receptor-p-value score = 1
- The weight used to multiply with the ligand activity score = 2
- The weight used to multiply with the condition-sender-specificity score of the ligand = 2
- The weight used to multiply with the condition-receiver-specificity score of the receptor= 2
- The weight used to multiply with the score reflecting the fraction of samples with sufficient expression per condition = 1

The output of the weighted aggregation step is stored in one final prioritization table which is given as part of the analysis output to the MultiNicheNet user. Other parts of the analysis output include data tables like the normalized pseudobulk expression table, the ligand activity table, and the ligand-target table. Based on these data tables, downstream visualizations can be generated as described in the corresponding section.

### MultiNicheNet applications

#### MultiNicheNet analysis on data from Bassez et al. (breast cancer case study)

The scRNA-seq read counts and metadata from the study from Bassez et al.^23^ were downloaded from https://lambrechtslab.sites.vib.be/en/single-cell, and used to create a SingleCellExperiment object^112^. We only used the data from cohort A because more detailed cell type annotations were provided for this cohort. From the lymphoid and myeloid detailed annotations, we generated more coarse-grained annotations to have sufficient cells per sample-cell-type combination. As “CD8 T cells”, we considered all cells labeled as: “CD8_EMRA”, “CD8_N”, “CD8_EX_Proliferating”, “CD8_EX”, “CD8_RM”, and “CD8_EM”. As “CD4 T cells”, we considered all cells labeled as: “CD4_EM”, “CD4_N”, “CD4_EX”, and “CD4_EX_Proliferating”. As “CD4 TREGs cells”, we considered all cells labeled as: “CD4_REG” and “CD4_REG_Proliferating”. As “NK cells”, we considered all “NK_CYTO” and “NK_REST” cells. As “gdT cells”, we considered “Vg9Vd2_gdT” and “gdT” cells. All myeloid cells with the labels “C1_CD14”, and “C2_CD16” were considered as “monocytes”, and all other non-DC-labeled myeloid cells as “macrophages”. These labels, together with the broad labels indicating other cell types (e.g., fibroblasts and endothelial cells) were used as input for MultiNicheNet. Sample ids were extracted from the “sample_id” metadata column. All analyses were performed with all cell types in the data and with the default parameters of MultiNicheNet, except that the non-corrected p-values were used. However, the cell types “Myeloid_cell” and “T_cell” were not included in the visualizations shown in this paper. We did not consider these cell types for visualizations because they consisted of non-deeply annotated myeloid and T cells.

For the analysis described in the results section “MultiNicheNet highlights critical pre-therapy cell-cell signaling patterns linked to therapy response in breast cancer patients”, condition labels were extracted from the “expansion_timepoint” metadata column, and the following contrasts were assessed: *“PreE – PreNE” and “PreNE – PreE”*.

For the multifactorial analysis described in the results section “MultiNicheNet elucidates differences in therapy-induced communication changes between and within response groups”, condition labels were extracted from the “expansion_timepoint” metadata column, and the following contrasts were assessed: *“[OnE – PreE] – [OnNE – PreNE]” and “[OnNE – PreNE] – [OnE – PreE]”*. Instead of the default p-values, p-values estimated from the empirical null distribution were used because the DE p-value distributions indicated that some model assumptions were violated (**Supplementary Figure 18**).

For the MultiNicheNet analysis to assess differential cell-cell communication within the group of non-expander patients, we divided the patients in three groups: “Angio” group consisting of patients with patient_ids “BIOKEY_26”, “BIOKEY_22”, “BIOKEY_14”, “BIOKEY_25, “BIOKEY_24”, “BIOKEY_8”, and “BIOKEY_19”, the “NO_angio” group consisting of patients with patient_ids “BIOKEY_21”, “BIOKEY_20”, “BIOKEY_4”, “BIOKEY_3”, and “BIOKEY_29”, and the rest group consisting of the other patients. The assessed contrasts were *“Angio – NO_angio” and “NO_angio – Angio”* and only cells from the pre-therapy samples were considered.

#### MultiNicheNet analysis on data from Li et al. (cSCC case study)

The Visium spatial transcriptomics and scRNA-seq read counts and metadata from the study from Li et al.^42^ were downloaded from https://www.ncbi.nlm.nih.gov/geo/query/acc.cgi?acc=GSE144240. scRNA-seq counts from all samples were merged into one matrix before making a SingleCellExperiment object^112^. From the most detailed annotations (“level3_celltype”), we generated more coarse-grained annotations to have sufficient cells per sample-cell-type combination. As “CD8 T cells”, we considered all cells labeled as: “CD8_EM”, “CD8_EMRA”, “CD8_Exh”, and “CD8_Naive”. As “CD4 T cells”, we considered all cells labeled as: “CD4_Exh”, “CD4_Naive”, “CD4_Pre_Exh”, and “CD4_RGCC”. As keratinocytes (cycling) we considered “Tumor_KC_Cyc” and “Normal_KC_Cyc”; as keratinocytes (basal) “Tumor_KC_Basal” and “Normal_KC_Basal”; as keratinocytes (differentiating) “Tumor_KC_Diff” and “Normal_KC_Diff”; as keratinocytes (other) “TSK” and “Keratinocyte”. Sample ids were extracted from the “sample_id” metadata column, and condition labels were extracted from the “tum.norm” metadata column. The following contrasts were assessed: *“Tumor – Normal” and “Normal – Tumor”*. The “patient” id was included as covariate to the model to correct for patient-specific effects. It was not included as batch effect because we did not want to correct the expression values themselves. The analysis was performed for all cell types and with the default parameters of MultiNicheNet, except for three parameters. The *min_cells* argument for the DE analysis was set to 5 instead of 10 because only three cell types would be retained in that case. The logFC_threshold was set to 0.75 to perform a slightly more stringent selection of the gene sets of interest because many genes are DE (probably because of the paired design and strong differences between normal and tumor tissue). However, non-corrected p-values were used instead of p-values corrected for multiple testing (because of the small sample size).

To assess spatial co-localization of MultiNicheNet-prioritized cell-cell interactions, we analyzed corresponding spatial transcriptomics data. In specific, we downloaded 10x Visium spatial transcriptomics data from the two replicates samples from the tumor of patient 4. Cell type proportions per spot were obtained by applying deconvolution with RCTD^113^ and cell2location^114^. The Spotless benchmark pipeline for deconvolution tools was used to run these deconvolution methods (https://github.com/saeyslab/spotless-benchmark)115. To define spatial co-localization of cell types, the Pearson correlation between cell type proportions across spots was calculated. To define a value of “interaction potential” between cell type pairs, we based ourselves on the number of prioritized differential interactions. Per condition of interest (tumor and normal), the *top n* ligand-receptor pairs were considered (n: 250, 500, 750, 1000). Additionally, *n* randomly chosen pairs and *n* pairs in the middle of the ranking were also considered as control. For each heterotypic cell-type pair (this is, pairs with different sender and receiver cell types), the number of ligand-receptor pairs in this set of *n* pairs was counted and divided by *n*. Hereby, we get which fraction of the total interactions are between a particular sender and receiver cell type. This fraction differs per cell type pair depending on the directionality of the interaction. For example, the fraction of interactions from cell type A to cell type B is different than from cell type B to cell type A. To get one fraction value per cell type pair, we considered the maximum fraction of both directions. This fraction was then plotted versus the spatial co-colocalization of the cell type pair to get an idea of potential spatial enrichment of the prioritized differential interactions. Figures in this paper were made by considering the RCTD output for spatial co-colocalization calculations. Conclusions were the same when using cell2location’s predictions (see code availability section).

#### MultiNicheNet analysis on data from Hoste et al. (MIS-C case study)

For the MIS-C case study, we used the scRNA-seq read counts and metadata from the study from Hoste et al^18^. Sample ids were extracted from the “ShortID” metadata column, cell type labels from the “Annotation_v2.0” column, and condition labels from the “MIS.C.AgeTier” column. The same cell type labels were used as indicated in that metadata column, except that we pooled “M_Monocyte_CD14_activated” and “M_Monocyte_CD14_resting” together as “CD14+ monocytes”, and “L_B_Memory” and “L_B_Memory_Unknown” as “Memory B cells”. The provided condition labels were adapted as follows for readability reasons: “Y_P” *(MIS-C: yes; pediatric)* becomes “M” *(MIS-C)*, “N_P” *(MIS-C: no; pediatric)* becomes “S” *(sibling)* and “N_A” *(MIS-C: no; adult)* becomes “A” *(adult COVID-19)*. The analysis performed with all cell types in the data and with the default parameters of MultiNicheNet, except that the non-corrected p-values were used. The following contrasts were assessed: *“M –[S+A]/2”, “S –[M+A]/2”, and “A –[S+M]/2”*.

#### MultiNicheNet analysis on integrated lung atlas data to study cell-cell communication in IPF

For the IPF case study, the dataset “Azimuth meta-analysis of 10 datasets of healthy and diseased human lungs” was downloaded from https://cellxgene.cziscience.com/collections/2f75d249-1bec-459b-bf2b-b86221097ced4142. Only datasets with profiles from healthy controls and patients with IPF were kept: Adams 2020, Reyfman 2019, Morse 2019, and Habermann 2020^42, 46–49^. Cell types with less than 5 cells per disease-dataset combination were omitted from the data. Sample ids were extracted from the “donor” metadata column, cell type labels from the “annotation.l1” column, condition labels from the “disease” column, and batch labels from the “dataset_origin” column. Batch effect correction for the dataset of origin was applied for both DE analysis and pseudobulk expression normalization as described in the section “MultiNicheNet algorithm description”. The analysis was performed with all cell types in the data and with the default parameters of MultiNicheNet, except that the *min_cells* parameter was set to 5 instead of 10. The following contrasts were assessed: *“idiopathic.pulmonary.fibrosis – normal”, “normal – idiopathic.pulmonary.fibrosis”*.

### MultiNicheNet visualizations

In this section, we will describe additional details about some of the types of plots that can be generated for the interpretation of the MultiNicheNet results.

#### Expression and ligand activity plots

In the plots showing the expression and activity of ligand-receptor pairs (e.g., Supplementary Figure 2), the log-transformed and normalized pseudobulk expression values of the ligand and receptor in a specific sample are multiplied. Subsequently, these values are scaled per ligand-receptor pair across all patients. We decided to visualize pseudobulk expression values instead of single-cell-derived expression values (as done in the *muscat* vignettes^22^) because DE analysis is performed on the pseudobulk expression values and we wanted the downstream visualizations to be as representative of the analysis as possible. Therefore, we created the option to calculate and visualize batch-corrected log-transformed and normalized pseudobulk expression values.

#### Ligand-target heatmaps filtered by expression correlation

Ligand-target links are inferred and visualized as described in the NicheNet publication^11^. To make ligand-target heatmap visualizations, we now provide the option to the user to filter ligand-target predictions based on across-sample expression correlation between the upstream ligand-receptor pair and the downstream target gene. Therefore, the Pearson and spearman correlation is calculated between 1) the product of log-transformed normalized pseudobulk expression values of ligand and receptor of each pair of interest, and 2) the target gene in the receiver cell type. We recommend that users only keep ligand-target links if they are supported by correlation for predicted upregulated target genes (e.g., Pearson or spearman correlation > 0.33) and anti-correlation for predicted downregulated genes (e.g., Pearson or spearman correlation < −0.33).

#### Intercellular regulatory networks

Intercellular regulatory networks are networks that consist of predicted ligand-target links for which the target gene encodes for a prioritized differential ligand or receptor. They are constructed based on the (correlation-filtered) ligand-target links. The networks shown in this paper were visualized using Cytoscape^116^.

#### Ligand-target signaling networks

Potential signaling paths between a ligand and a set of target genes can be predicted and visualized as described in the NicheNet publication^11^. For the networks In this paper, we used the function *nichenetr::get_ligand_signaling_path_with_receptor* with the parameter *top_n_regulators = 2* such that only the top two predicted transcriptional regulators per target gene are shown to keep the network small.

## Data availability

No new data were generated for this study. All data used in this study are publicly available as described in the Methods section “MultiNicheNet applications”. The NicheNet-v2 networks and ligand-target matrix are available at Zenodo (https://zenodo.org/record/7074291). Databases used to create the NicheNet-v2 model are mentioned and referred to in the Methods section “Construction of the NicheNet-v2 model” and corresponding Supplementary Note 1 and Supplementary Table 1.

## Code availability

An open-source R implementation of MultiNicheNet is available at GitHub (https://github.com/saeyslab/multinichenetr). The release includes tutorials and example vignettes for the following analyses: a classic pairwise comparison, a pairwise comparison between two paired conditions, a comparison between multiple conditions, a complex multifactorial comparison investigating differences in cell-cell communication changes, and an analysis that corrects for batch effects. Moreover, the package tutorials of the original NicheNet software (https://github.com/saeyslab/nichenetr) have been adapted to include the new NicheNet-v2 model. The model evaluation and benchmarking tutorial has also been updated to include the CytoSig dataset of gold standard ligand-target links as well. Finally, code and data to reproduce the analyses from this study are available at Zenodo (https://zenodo.org/record/8016880).

## Acknowledgments

R.B. has been supported by a Ph.D. fellowship from The Research Foundation - Flanders (grant number 1181318N) and a SBO project grant (grant number S001121N). This research also received funding from the Flemish Government under the “Onderzoeksprogramma Artificiële Intelligentie (AI) Vlaanderen” program. The authors would also like to thank S. Verbandt (KU Leuven) and D. Dimitrov (University of Heidelberg) for helpful discussions about this project.

## Author Contributions

R.B. conceived and implemented MultiNicheNet with input from R.S., J.G., and Y.S. R.B. and C.S. performed the experiments with input from R.S. and Y.S. P.D.B. contributed to the construction of NicheNet-v2. S.T., L.H., and D.L. contributed to the interpretation of the case study results. R.S. and Y.S. supervised the work. R.B. wrote the manuscript with input from the co-authors who all approved the manuscript before submission.

