## Supplementary Figures for "MultiNicheNet: a flexible framework for differential cell-cell communication analysis from multi-sample multi-condition single-cell transcriptomics data"

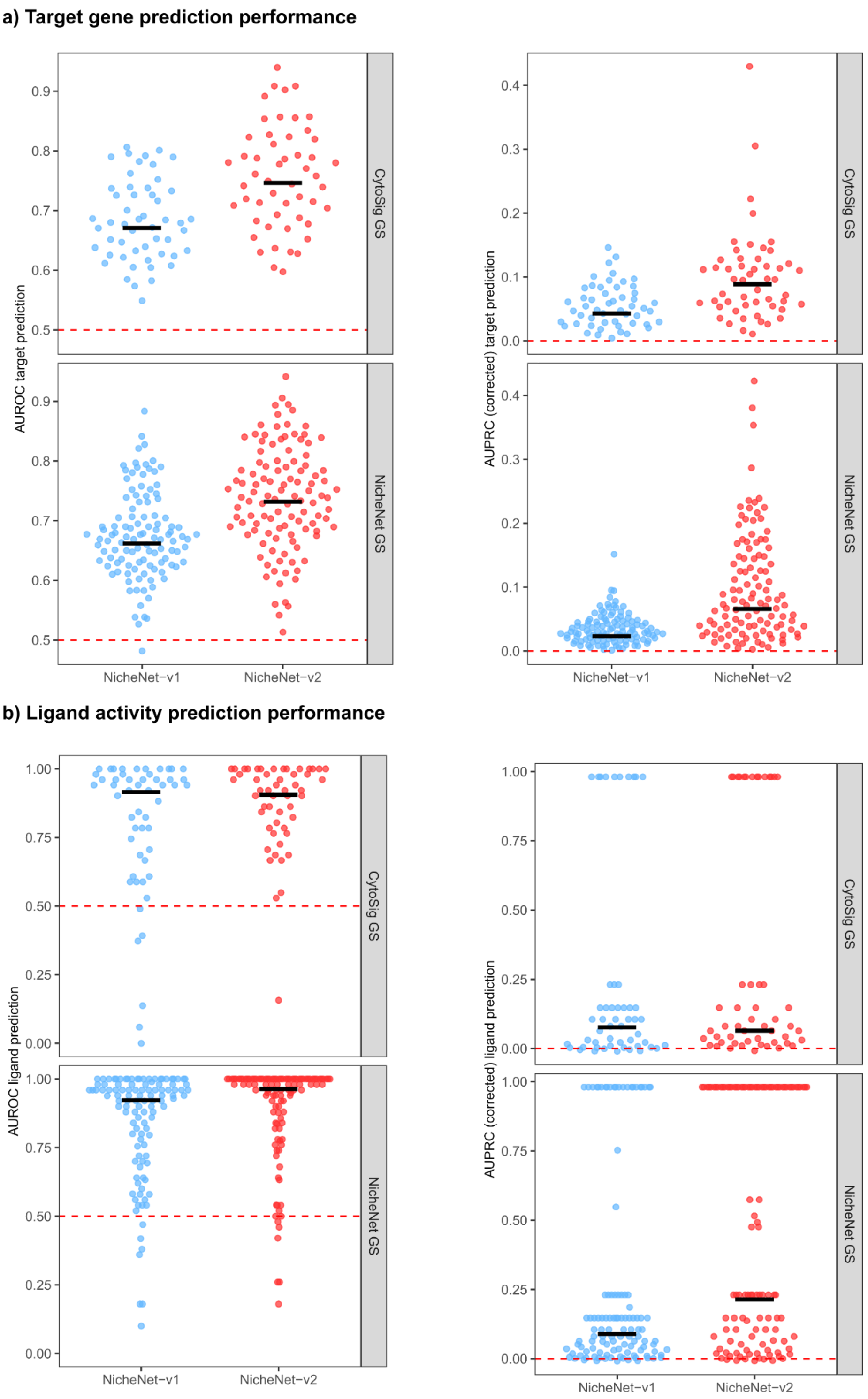


**Supplementary Figure 1 | Comparison of NicheNet-v2 and NicheNet-v1 in target gene and ligand activity prediction demonstrates that NicheNet-v2 provides a more accurate model of ligand-target regulation. a)**, Evaluation of target gene prediction performance: For 52 CytoSig and 111 NicheNet ligand treatment datasets providing transcriptome data of cells after *in vitro* ligand treatment, we assessed NicheNet-v1 and NicheNet-v2 model performance in predicting which genes are differentially expressed on treatment with a particular ligand. Each dot indicates the performance for one dataset, and the black line indicates the median performance over all datasets. **b)** Evaluation of ligand activity prediction performance: for the same datasets, we compared the performance of NicheNet-v1 and NicheNet-v2 in predicting by which ligand the cells were treated based on the differentially expressed genes after treatment. Visual representation of the performance is the same as in a). The red dashed line indicates the performance of random guessing. AUROC: area under the receiver-operating-characteristic curve; AUPRC (corrected): area under the precision-recall curve, corrected for random prediction; GS: gold standard.

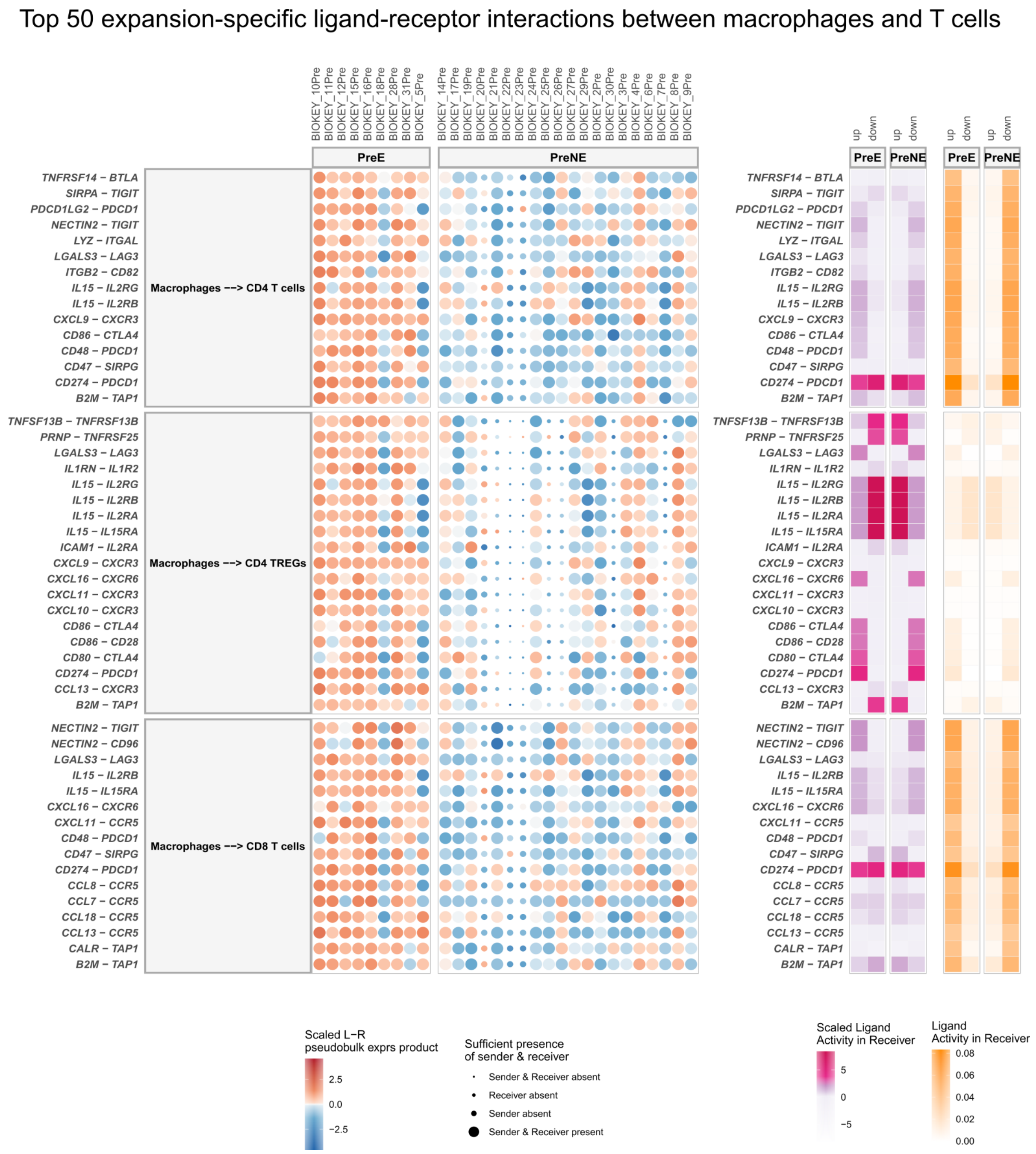


**Supplementary Figure 2 | MultiNicheNet prioritizes cell-cell communication patterns between macrophages and T cells that are specific for patients with clonotype expansion after anti-PD1 therapy.** MultiNicheNet was applied to scRNA-seq from Bassez et al. to compare pre-therapy cell-cell communication between expander (preE) and non-expander patients (preNE). The top 50 preE-specific interactions from macrophages to T cells are shown. For each interaction, ligand-receptor pseudobulk expression (product of normalized log values) and (scaled) ligand activity values are visualized. Ligand activity values are the AUPRC scores indicating the performance in predicting the up- or downregulated genes in the preE or preNE group. Scaled ligand activities are z-score normalized ligand activity values, calculated per receiver cell type. The higher these values, the more enriched target genes of a specific ligand are compared to other ligands. The size of the dots indicates whether a sample had enough cells (>= 10) for a specific cell type to be considered for DE analysis. L-R: ligand-receptor.

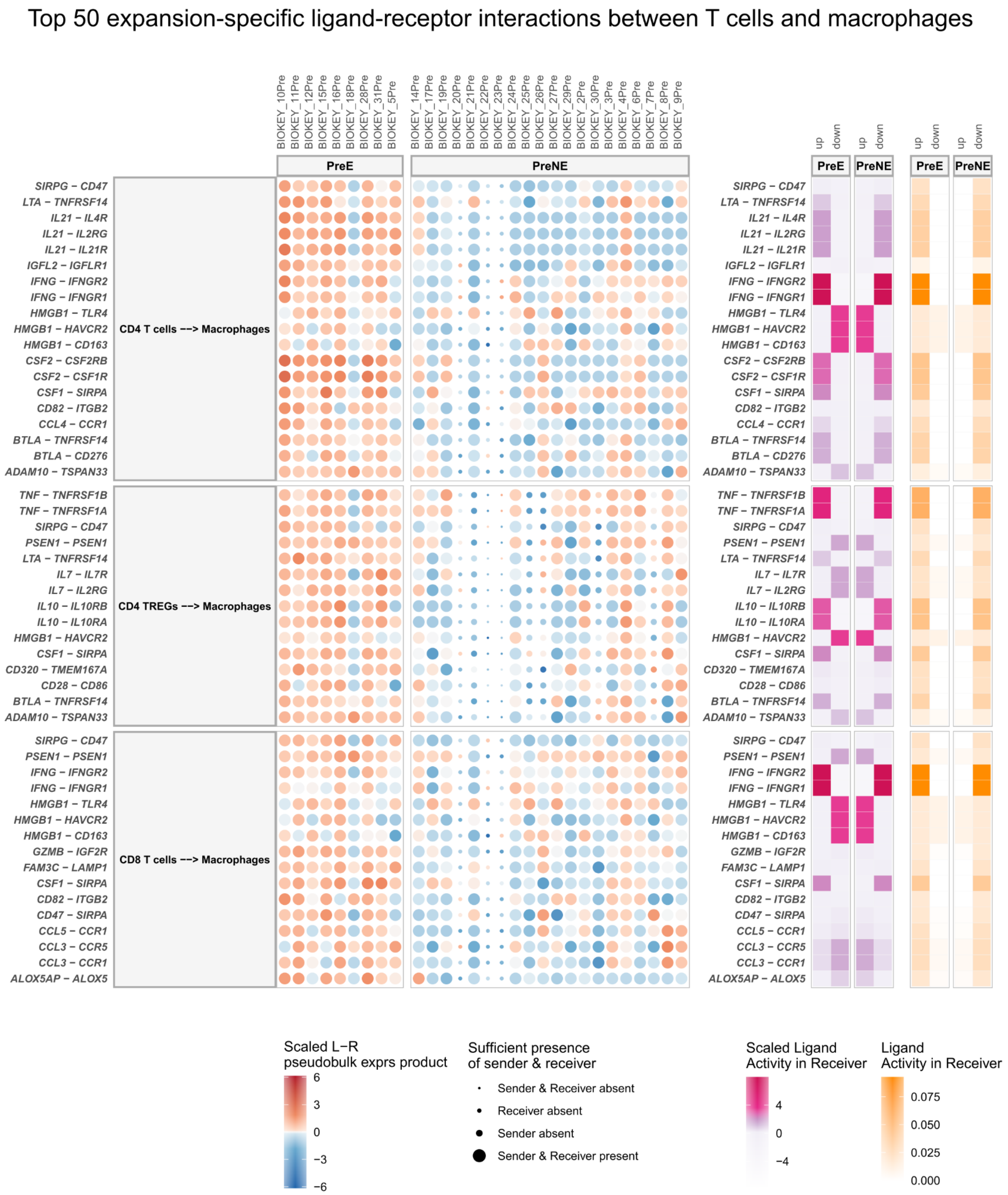


**Supplementary Figure 3 | MultiNicheNet prioritizes cell-cell communication patterns between T cells and macrophages that are specific for patients with clonotype expansion after anti-PD1 therapy.** MultiNicheNet was applied to scRNA-seq from Bassez et al. to compare pre-therapy cell-cell communication between expander (preE) and non-expander patients (preNE). The top 50 preE-specific interactions from T cells to macrophages are shown. For each interaction, ligand-receptor pseudobulk expression (product of normalized log values) and (scaled) ligand activity values are visualized. Ligand activity values are the AUPRC scores indicating the performance in predicting the up- or downregulated genes in the preE or preNE group. Scaled ligand activities are z-score normalized ligand activity values, calculated per receiver cell type. The higher these values, the more enriched target genes of a specific ligand are compared to other ligands. The size of the dots indicates whether a sample had enough cells (>= 10) for a specific cell type to be considered for DE analysis. L-R: ligand-receptor.

### Predicted target genes of expansion-specific ligand-receptor pairs [Macrophages ⟶ CD8 T cells]

a) Expression of prioritized ligand-receptor pairs and their predicted target genes.

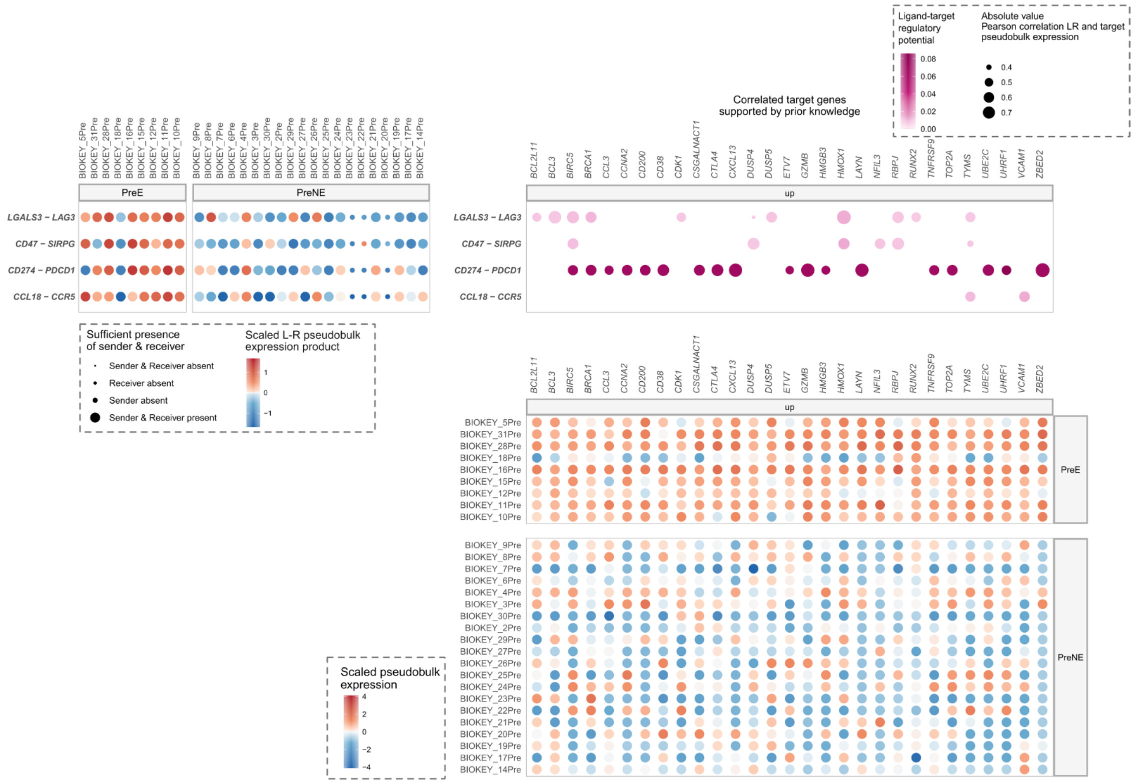


b) CD274 - PDCD1 expression versus target gene expression.

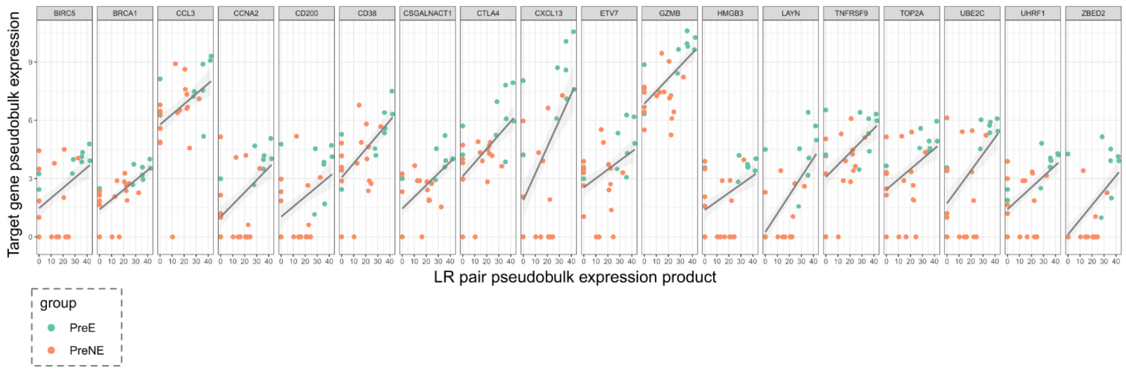


**Supplementary Figure 4 | MultiNicheNet enables exploration of target genes predicted downstream of prioritized ligand-receptor interactions. a)** Visualization of a subset of predicted target genes downstream of the MultiNicheNet-prioritized expander-specific ligand-receptor pairs (from macrophages to CD8 T cells). Shown are the same ligand-receptor pairs as depicted in Figure 2. The depicted target genes in CD8 T cells are genes that are upregulated in expander patients (preE) versus non-expander patients (preNE) (p-value <= 0.05 & log2FC > 1) and show across-patient expression correlation with the upstream predicted ligand-receptor pair (Pearson correlation > 0.50 or Spearman correlation > 0.50). The bubble plots indicate ligand-receptor pseudobulk expression (product of normalized log values) (top left), NicheNet-v2 ligand-target regulatory potential and expression correlation between ligand-receptor and target gene expression (top right), and target gene pseudobulk expression (bottom right). **b)** Visualization of ligand-receptor and target gene correlation for the ligand-receptor pair CD274-PDCD1 (PDL1-PD1). Shown are the same target genes as depicted in a). Each dot represents a patient and is colored according to the patient group. The black smoothing lines are the result of fitting a linear regression model. LR: ligand-receptor.

Ligand-to-target signaling network between CD274 and *UHRF1*, *CTLA4*, *BRCA1* and *CSGALNACT1*

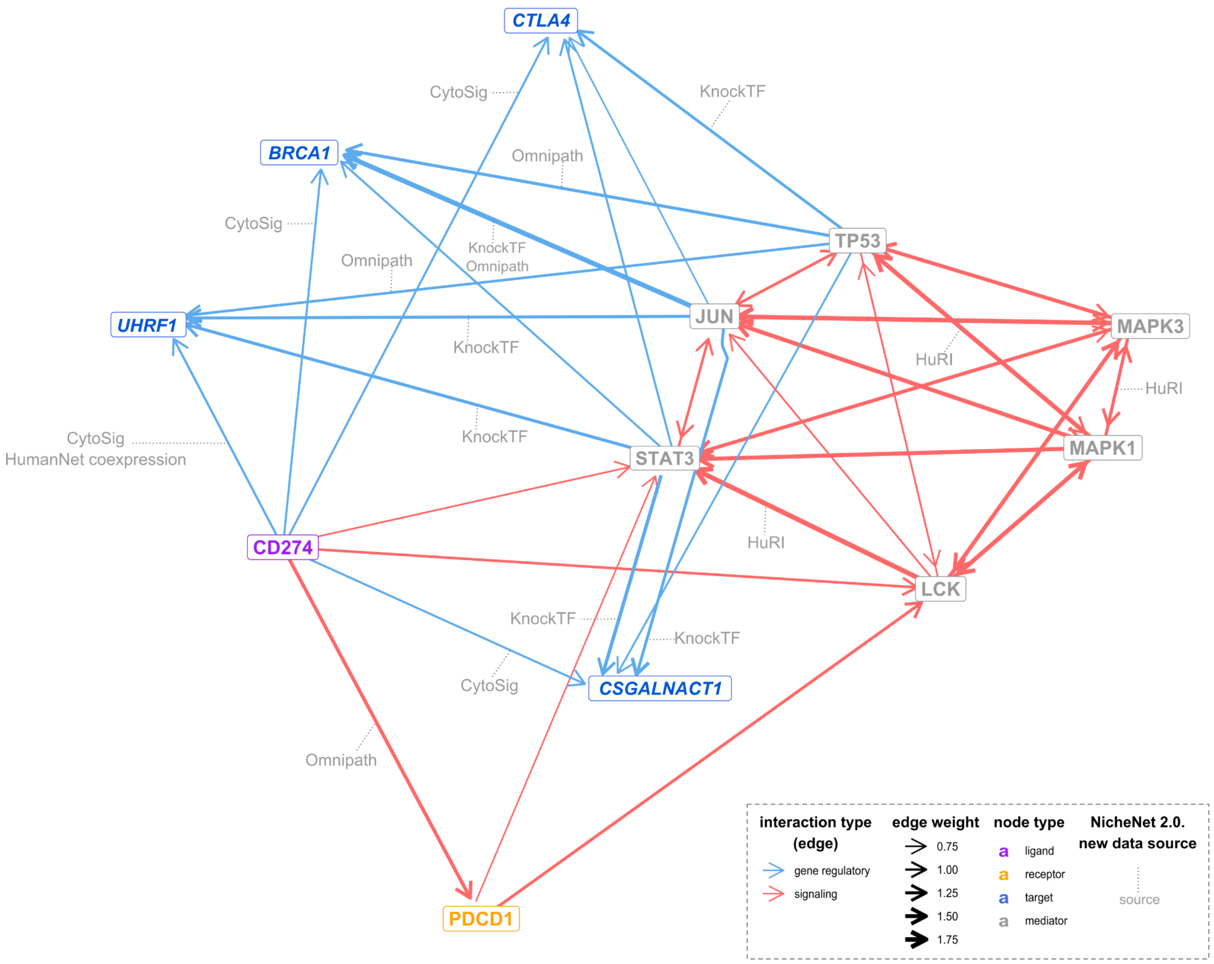


**Supplementary Figure 5 | Inference of putative signaling paths from a ligand of interest to target genes of interest.** Using NicheNet-v2's integrated networks, users can verify which putative signaling and gene regulatory interactions support the predicted ligand-target links. This example shows links from the ligand CD274 to its predicted target genes *UHRF1*, *CTLA4*, *BRCA1,* and *CSGALNACT1* (a subset of the targets that are shown in Supplementary Figure 4). The color scale is explained in the box at the bottom right. Edge line thickness is proportional to the weight of the represented interaction in the weighted integrated networks. When NicheNet-v2 data sources support a shown interaction, this is indicated in the figure.

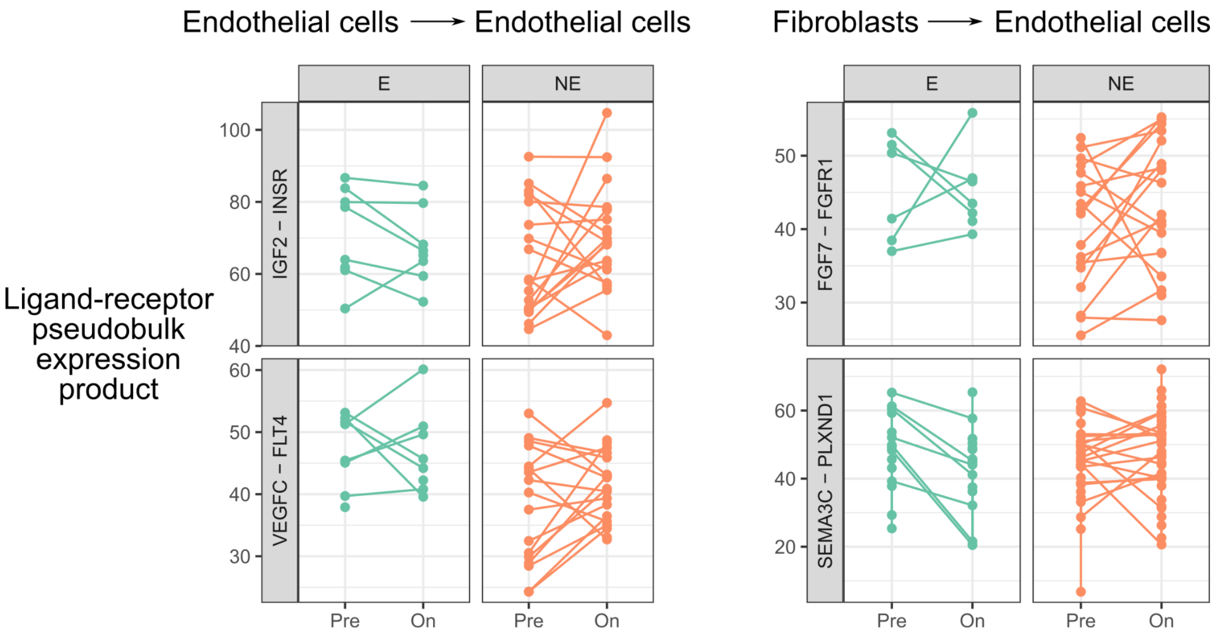


**Supplementary Figure 6 | Line plot revealing within-group differences in cell-cell communication changes in response to anti-PD1 therapy.** MultiNicheNet was applied to scRNA-seq from Bassez et al. to compare on-therapy versus pre-therapy differences in cell-cell communication between expander (E - green color) and non-expander patients (NE - orange color). Each dot represents the ligand-receptor pseudobulk expression value (product of normalized log values) for one particular ligand-receptor pair in one sample. The pre-therapy and on-therapy samples of the same patients are through a line. The cell type indications at the top depict between which sender→receiver cell types the displayed ligand-receptor pairs take place.

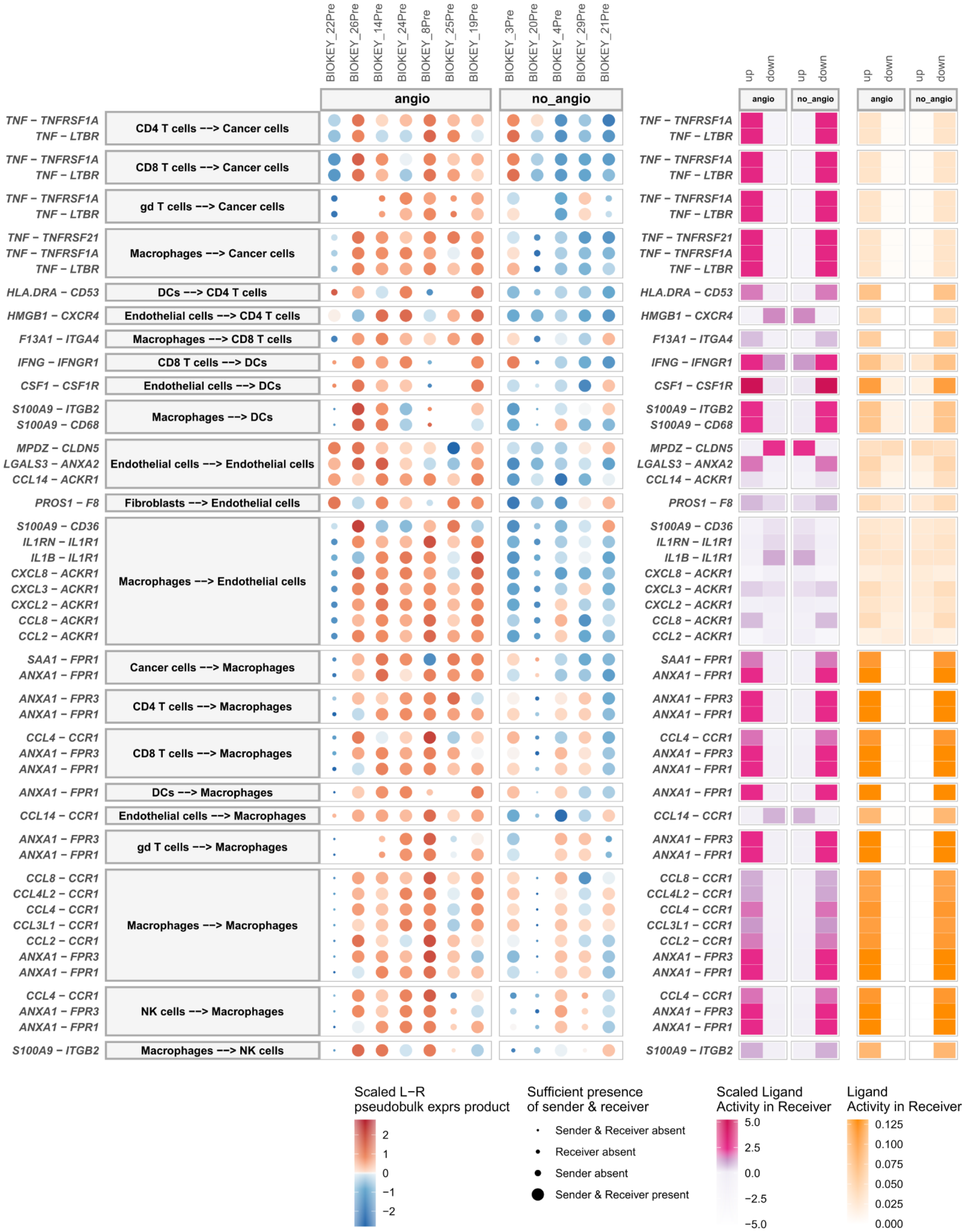


**Supplementary Figure 7 | MultiNicheNet prioritizes cell-cell communication patterns that are specific for the subset of non-expander patients in which angiogenesis-related signals increased during anti-PD1 therapy.** MultiNicheNet was applied to scRNA-seq from Bassez et al. to compare pre-therapy cell-cell communication between non-expander patients in which angiogenesis-related signals increased during anti-PD1 therapy ("angio") and non-expander patients in which these signals were decreased ("no_angio")(see Figure 3). The top 50 angio-specific interactions are shown. For each interaction, ligand-receptor pseudobulk expression (product of normalized log values) and (scaled) ligand activity values are visualized. Ligand activity values are the AUPRC scores indicating the performance in predicting the up- or downregulated genes in the angio or no_angio group. Scaled ligand activities are z-score normalized ligand activity values, calculated per receiver cell type. The higher these values, the more enriched target genes of a specific ligand are compared to other ligands. The size of the dots indicates whether a sample had enough cells (>= 10) for a specific cell type to be considered for DE analysis. Absence of a dot means there were no cells at all. L-R: ligand-receptor.

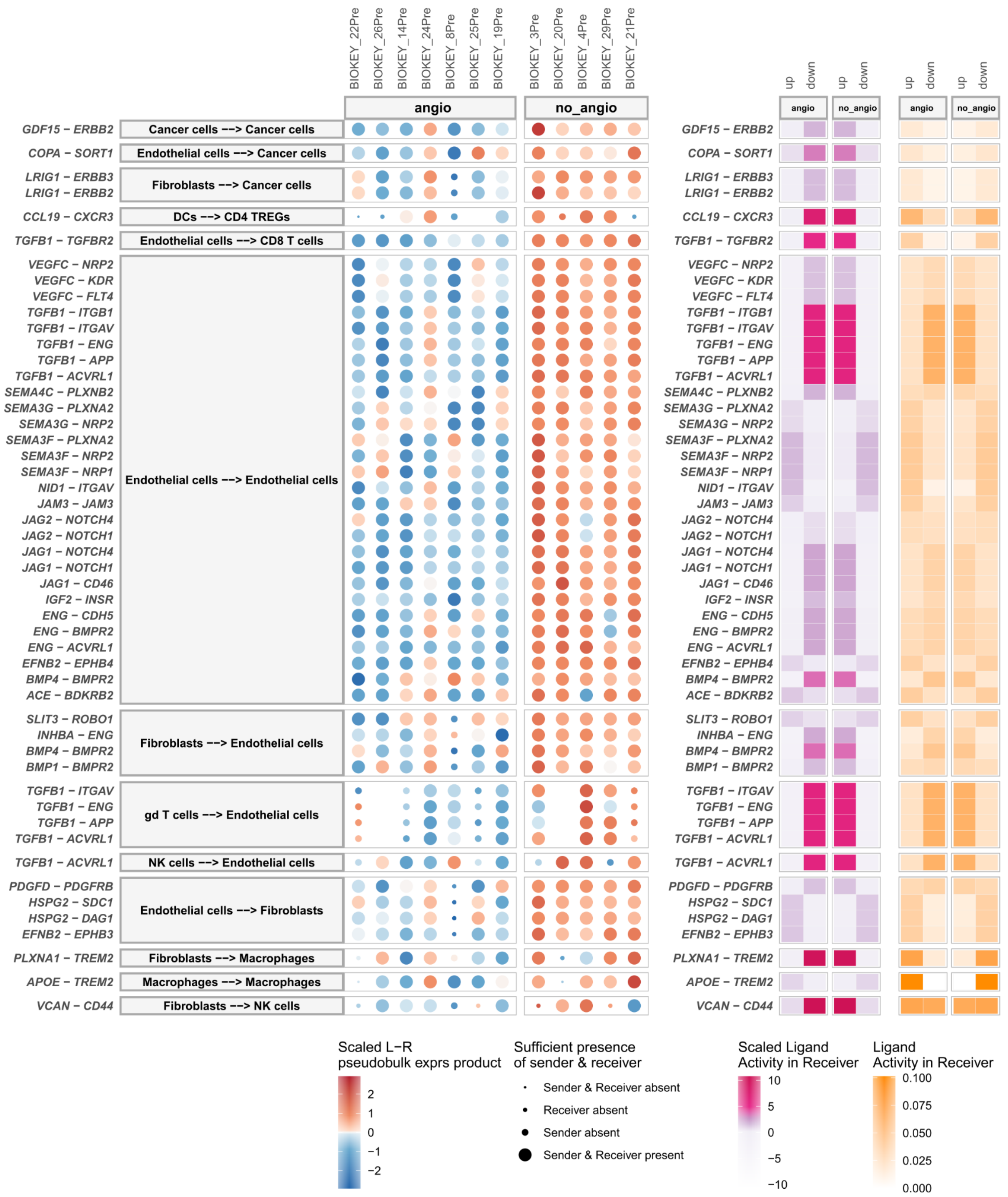


**Supplementary Figure 8 | MultiNicheNet prioritizes cell-cell communication patterns that are specific for the subset of non-expander patients in which angiogenesis-related signals decreased during anti-PD1 therapy.** MultiNicheNet was applied to scRNA-seq from Bassez et al. to compare pre-therapy cell-cell communication between non-expander patients in which angiogenesis-related signals increased during anti-PD1 therapy (“angio”) and non-expander patients in which these signals were decreased (“no_angio”)(see Figure 3). The top 50 no_angio-specific interactions are shown. For each interaction, ligand-receptor pseudobulk expression (product of normalized log values) and (scaled) ligand activity values are visualized. Ligand activity values are the AUPRC scores indicating the performance in predicting the up- or downregulated genes in the angio or no_angio group. Scaled ligand activities are z-score normalized ligand activity values, calculated per receiver cell type. The higher these values, the more enriched target genes of a specific ligand are compared to other ligands. The size of the dots indicates whether a sample had enough cells (>= 10) for a specific cell type to be considered for DE analysis. Absence of a dot means there were no cells at all. L-R: ligand-receptor.

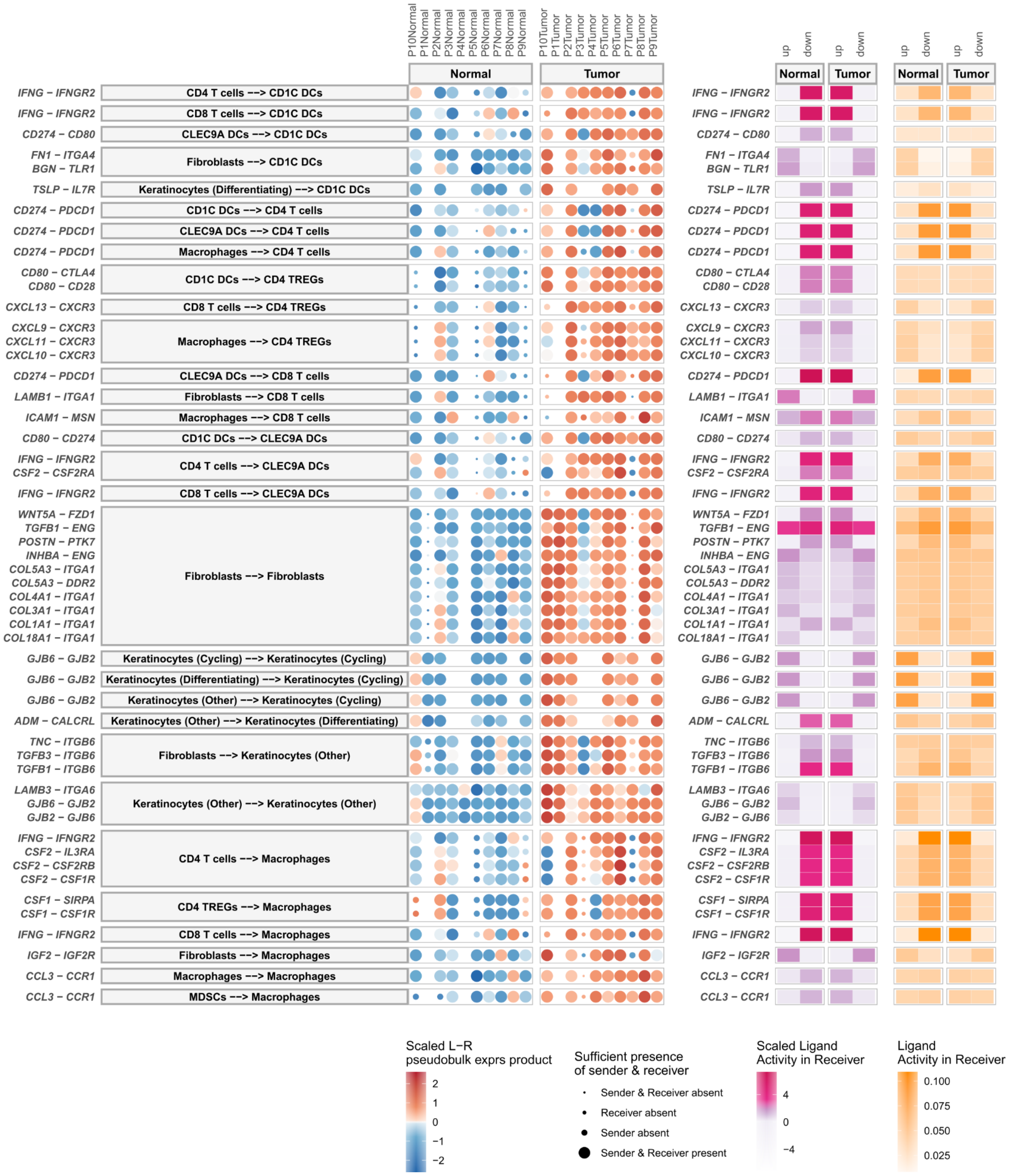


**Supplementary Figure 9 | MultiNicheNet prioritizes cell-cell communication patterns specific for cutaneous squamous cell carcinoma compared to matched healthy skin from the same patient.** MultiNicheNet was applied to scRNA-seq from Ji et al. to compare cell-cell communication between cutaneous squamous cell carcinoma tumor tissue (tumor) and healthy skin (normal). The tumor-specific interactions out of the total top 75 differential interactions are shown. For each interaction, ligand-receptor pseudobulk expression (product of normalized log values) and (scaled) ligand activity values are visualized. Ligand activity scores are the AUPRC scores in predicting the up- or downregulated genes in the tumor or normal condition. Scaled ligand activities are z-score normalized ligand activity values, calculated per receiver cell type. The higher these values, the more enriched target genes of a specific ligand are compared to other ligands. The size of the dots indicates whether a sample had enough cells (>= 5) for a specific cell type to be considered for DE analysis. Absence of a dot means there were no cells at all. L-R: ligand-receptor.

**Tumor-specifc intercellular signaling cascades in squamous cell carcinoma versus matched normal skin**

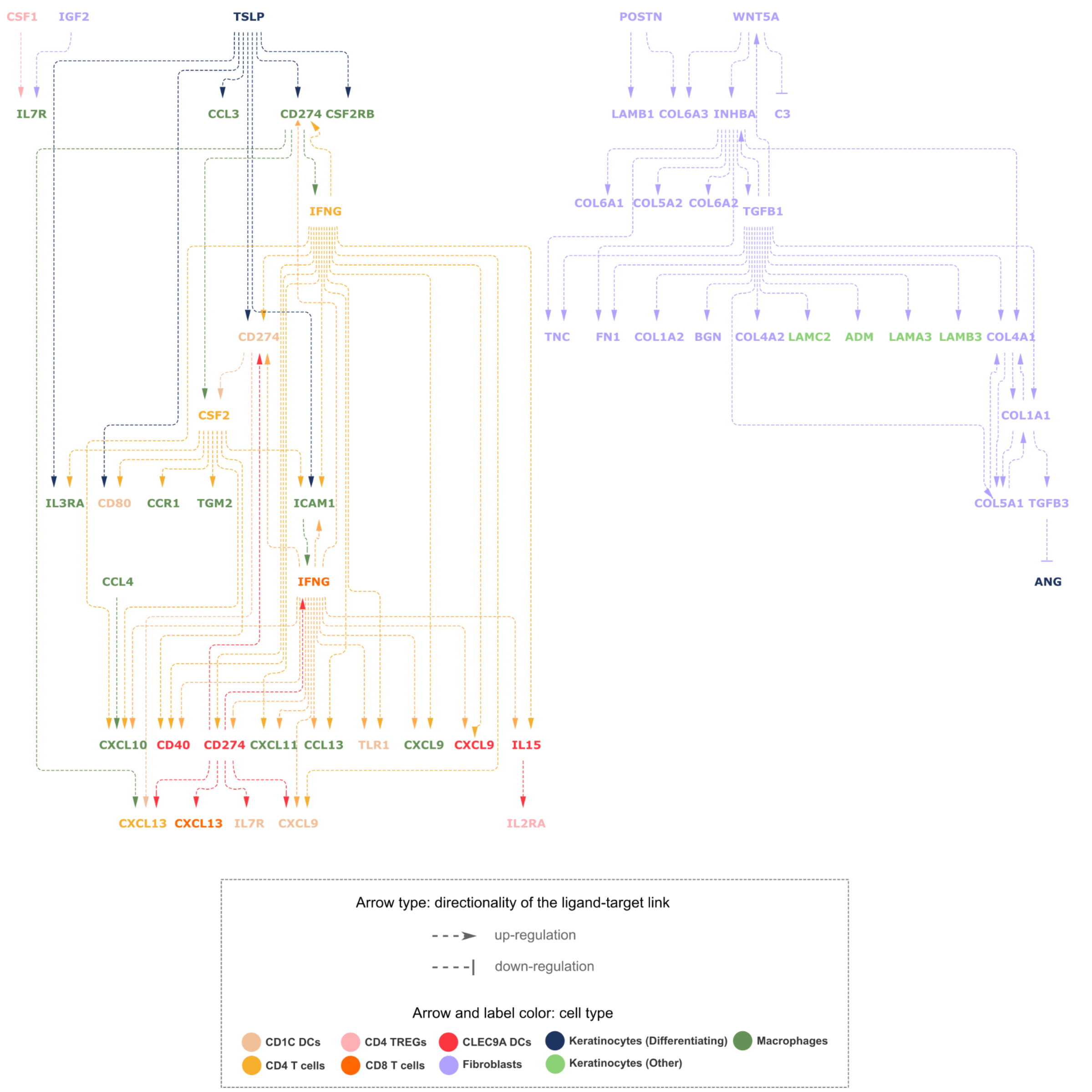


**Supplementary Figure 10 | MultiNicheNet predicts a network of intercellular regulatory interactions specific to cutaneous squamous cell carcinoma compared to healthy skin.** MultiNicheNet was applied to scRNA-seq from Ji et al. to compare cell-cell communication between cutaneous squamous cell carcinoma tumor tissue (tumor) and healthy skin (normal). We searched for ligands or receptors of which the encoding gene is a target of another ligand-receptor pair of these top tumor-specific interactions. The links in this network should thus be considered as a gene regulatory ligand-target link. Specifically, we consider the tumor-specific ligand-receptor pairs in the top 150 most differential interactions. Target genes should be among the 250 genes with the highest regulatory potential to be regulated by the specific ligand and should show expression correlation with the specific upstream ligand-receptor pair (Pearson or Spearman correlation > 0.50). A predicted "up-regulatory" link indicates a positive across-patients expression correlation between the ligand-receptor pair of the ligand and the prior-knowledge-supported target gene in the receiver cell type. A predicted "down-regulatory" link indicates an anti-correlation between the expression of the ligand-receptor pair of the ligand and the target gene in the receiver cell type.

**a) TSLP - IL7R expression vs target gene expression between "Keratinocytes (Differentiating)" and "CD1C DCs"**

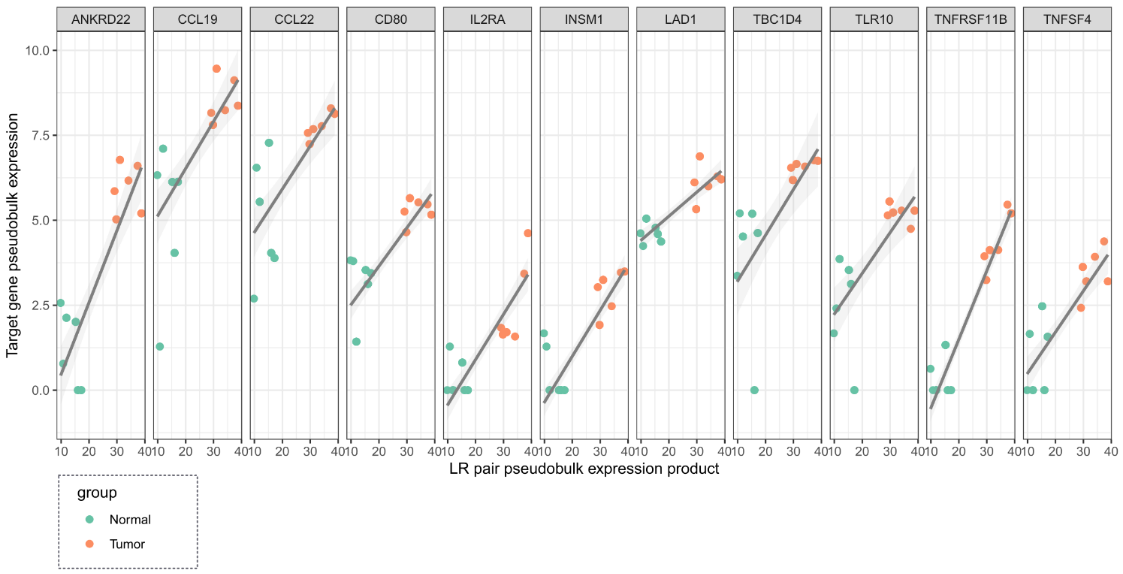


**b) Ligand-to-target signaling network between TSLP and a subset of its predicted target genes in CD1C DCs**

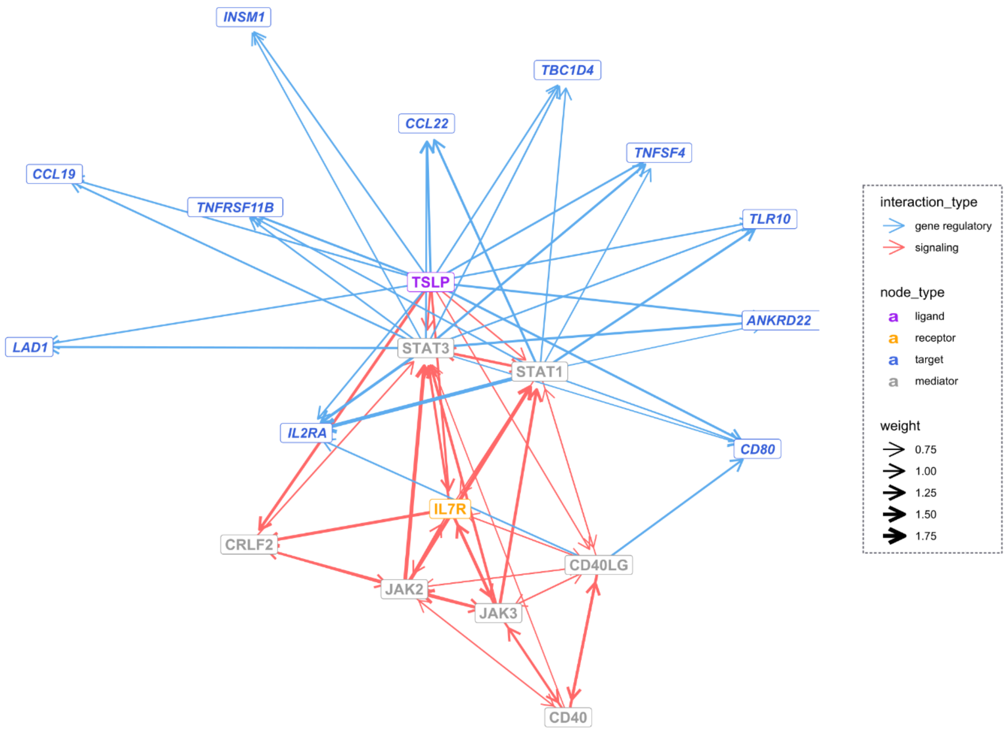


**Supplementary Figure 11 | Exploration of the observed tumor-specific TSLP signature in CD1C dendritic cells (DCs) in cutaneous squamous cell carcinoma. a)** Expression correlation between the ligand-receptor pair TSLP-IL7R and a subset of predicted target genes in CD1C DCs (Pearson or Spearman correlation > 0.75). The depicted target genes are among the 25 genes with the most prior knowledge to be regulated by TSLP, and TSLP is in the top 3 of ligands with the most prior knowledge to regulate them. Each dot represents a sample and is colored according to the tissue type. The black smoothing lines are the result of fitting a linear regression model. LR: ligand-receptor. **b)** putative signaling paths from the ligand TSLP to the target genes shown in a). Using NicheNet-v2's integrated networks, users can verify which putative signaling and gene regulatory interactions support the predicted ligand-target links. Edge line thickness is proportional to the weight of the represented interaction in the weighted integrated networks.

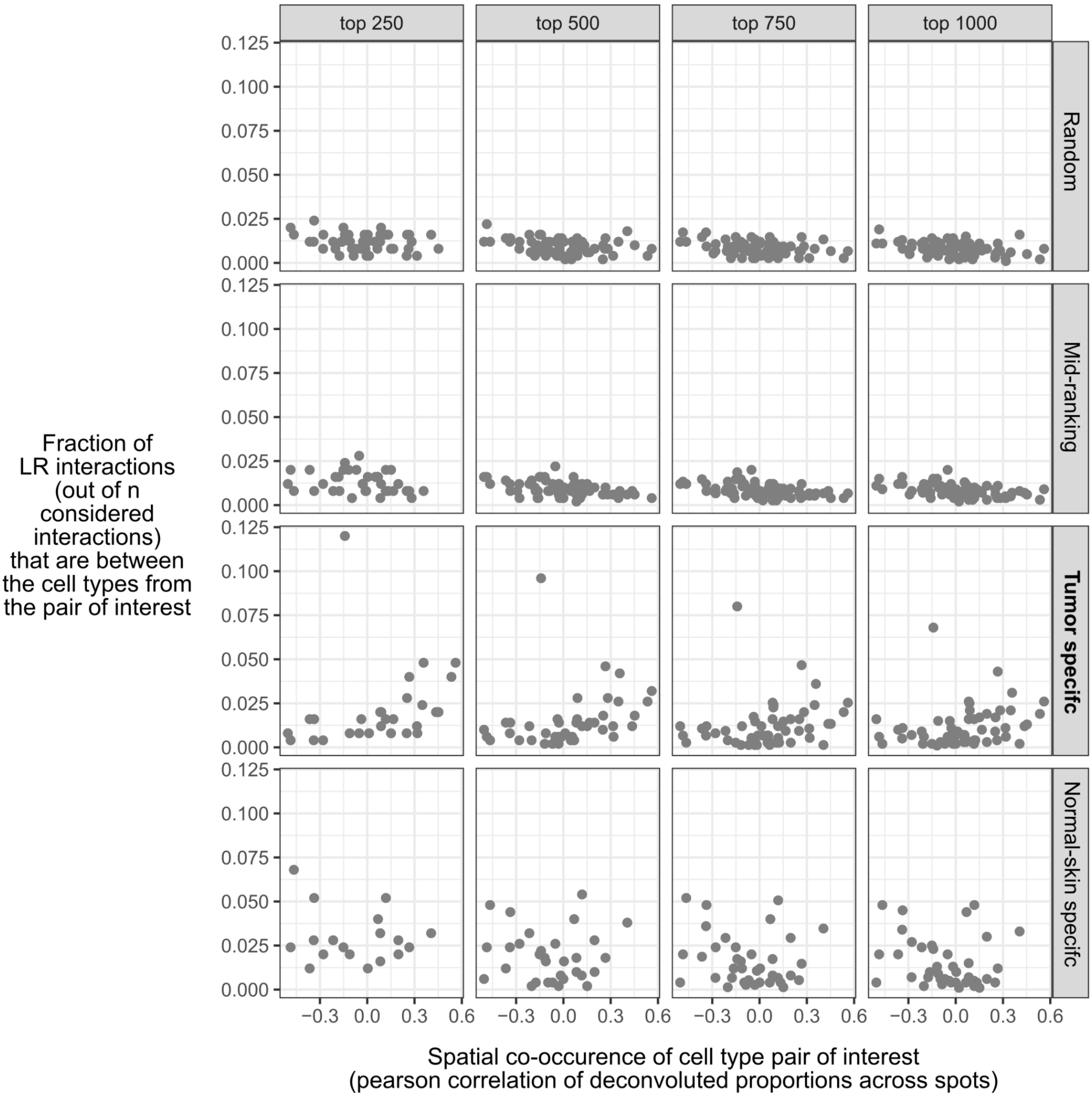


**Supplementary Figure 12 | Spatial co-localization analysis of MultiNicheNet-prioritized cell-cell communication patterns in cutaneous squamous cell carcinoma.** MultiNicheNet was applied to scRNA-seq from Ji et al. to compare cell-cell communication between cutaneous squamous cell carcinoma tumor tissue (tumor) and healthy skin (normal). For the top *n* (250, 500, …) tumor-specific ligand-receptor interactions, we retrieved the cell types involving these interactions. Next, we counted the number of interactions between each combination of sender-receiver cell type pairs, took the maximum of both directions (e.g., maximum of cell type A⟶celltype B and cell type B⟶ cell type A), and calculated the fraction of this number versus the total number of interactions (y-axis). This was also done for normal skin-specific interactions, a random set of *n* interactions, and a set of *n* interactions in the middle of the MultiNicheNet ranking. Spatial localization of cell types was determined by applying RCTD for deconvolution of the 10x Visium spatial transcriptomics data of tumor tissue (from one patient). Co-localization between cell types was quantified by the across-spot Pearson correlation of cell type proportions (x-axis). Each dot represents a cell-type pair with a certain spatial correlation value and fraction of interactions among the top *n*.

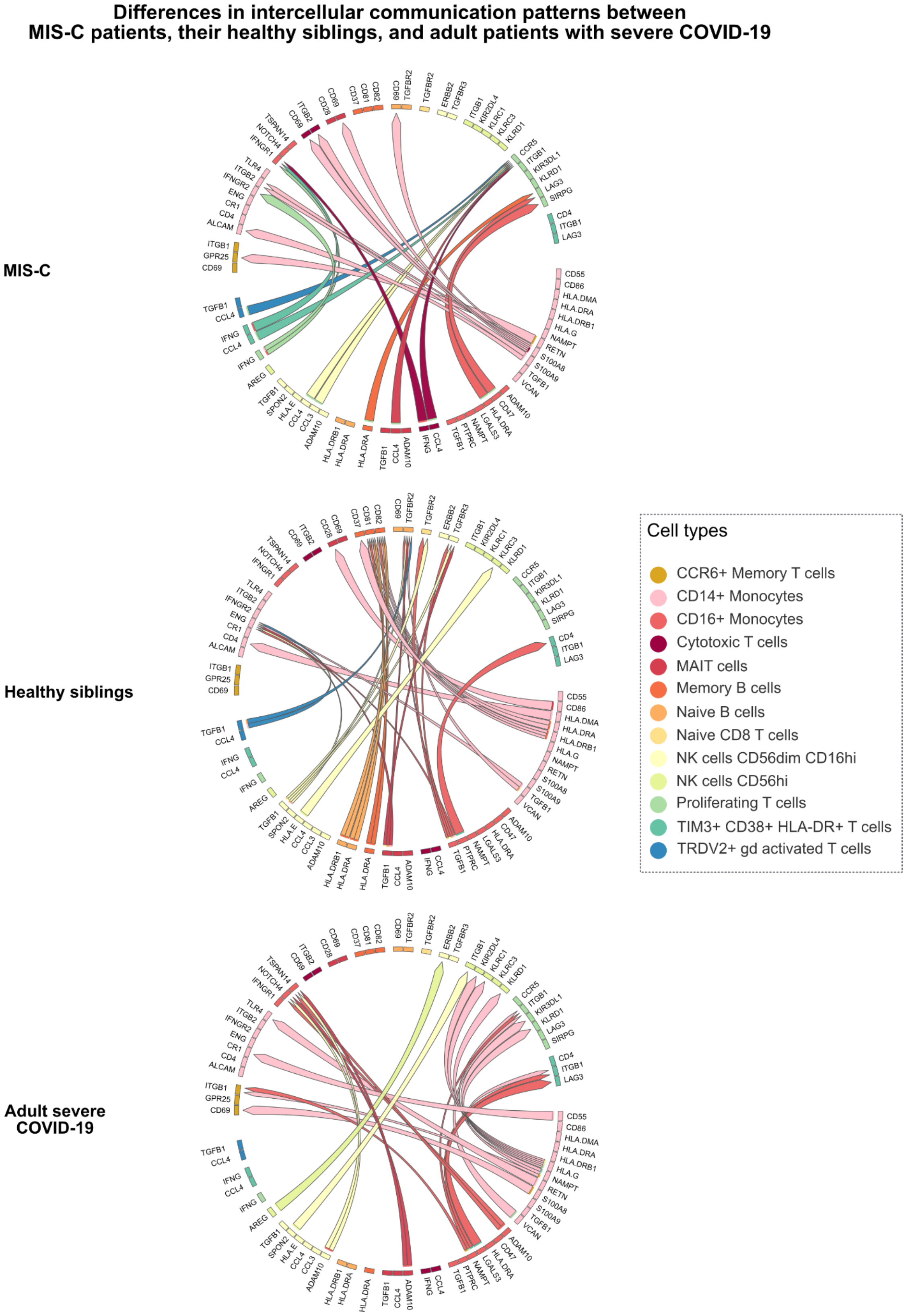


**Supplementary Figure 13 | MultiNicheNet prioritizes differential cell-cell communication patterns between MIS-C patients, their healthy siblings, and adult COVID-19 patients.** MultiNicheNet was applied to scRNA-seq from Hoste et al. to compare cell-cell communication among PBMCs between MIS-C patients, their healthy siblings, and adult COVID-19 patients. The top 75 differential ligand-receptor pairs are depicted in chord diagrams, divided per patient group. The arrowhead indicates the direction from sender to receiver cell type, and the color of the arrow indicates the sender cell type that expresses the ligand.

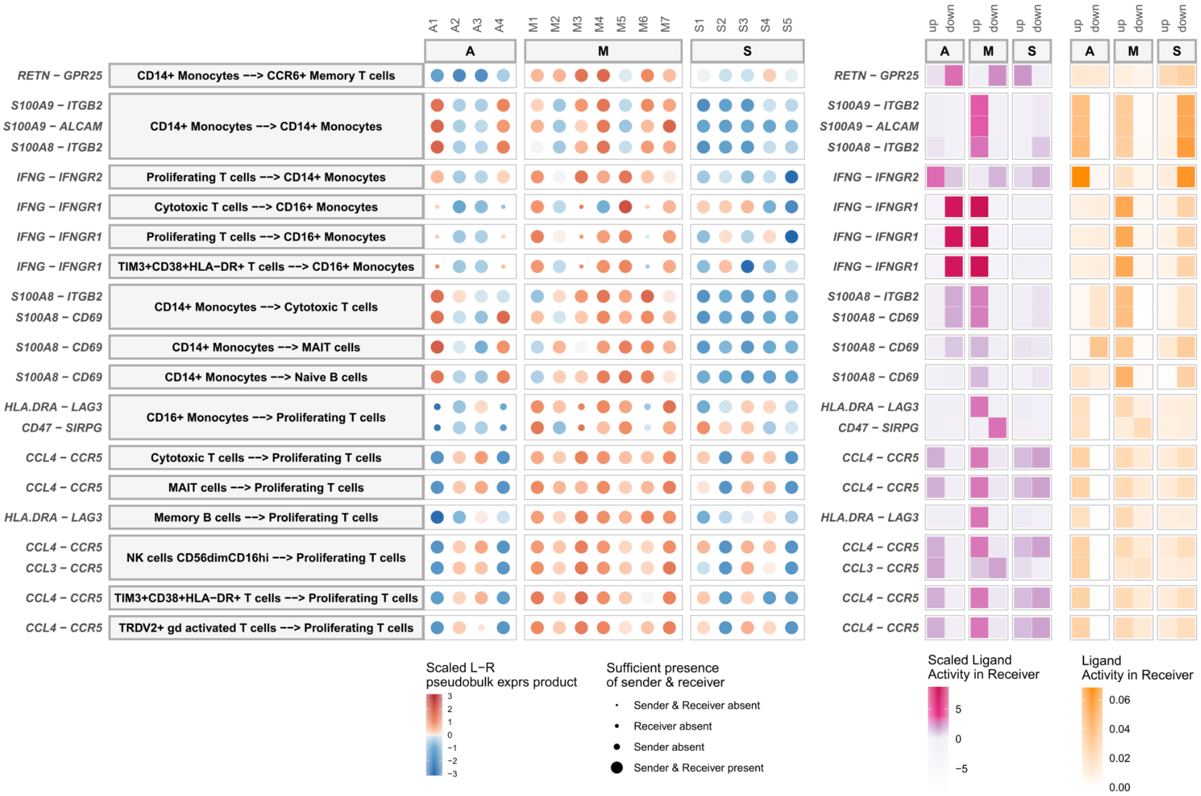


**Supplementary Figure 14 | MultiNicheNet prioritizes cell-cell communication patterns specific for MIS-C patients compared to healthy siblings and adult COVID-19 patients.** MultiNicheNet was applied to scRNA-seq from Hoste et al. to compare cell-cell communication among PBMCs between MIS-C patients (M), their healthy siblings (S), and adult COVID-19 patients (A). The MIS-C-specific interactions out of the total top 75 differential interactions are shown (i.e., the same interactions as in Supplementary Figure 13). For each interaction, ligand-receptor pseudobulk expression (product of normalized log values) and (scaled) ligand activity values are visualized. Ligand activity scores are the AUPRC scores in predicting the up- or downregulated genes in the A, M, or S patient group. Scaled ligand activities are z-score normalized ligand activity values, calculated per receiver cell type. The higher these values, the more enriched the target genes of a specific ligand are compared to other ligands. The size of the dots indicates whether a sample had enough cells (>= 10) for a specific cell type to be considered for DE analysis. L-R: ligand-receptor.

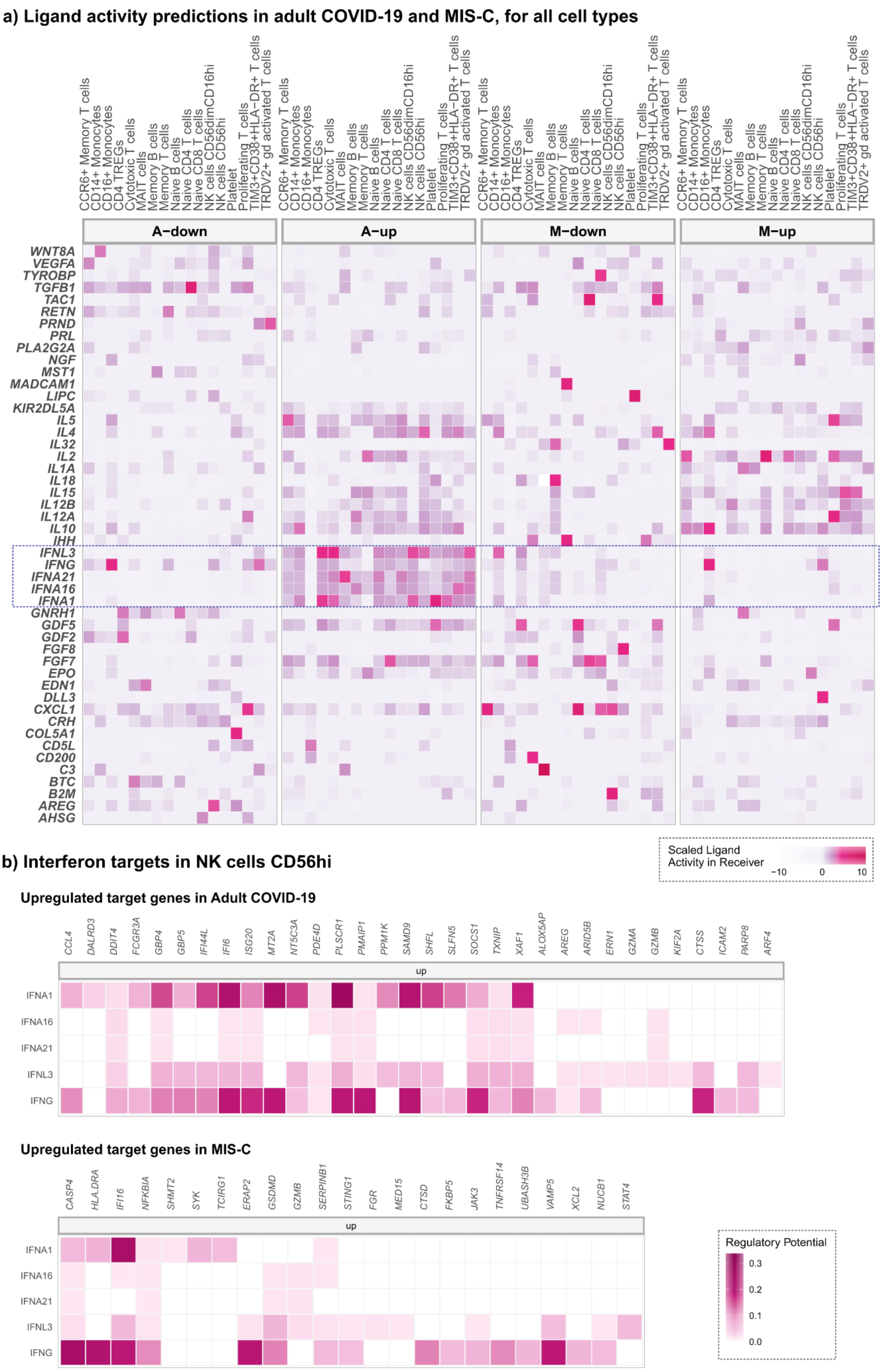


**Supplementary Figure 15 | Calculation of ligand activities in PBMCs from MIS-C patients and adult COVID-19 patients. a)** The ligand activity calculation step of MultiNicheNet was applied to scRNA-seq from Hoste et al. to calculate ligand activities in PBMCs from MIS-C patients (M) and adult COVID-19 patients (A). Shown are all ligands in the NicheNet-v2 database that are at least once predicted as the most active ligand for a particular cell-type-condition combination. Ligand activity scores are the AUPRC scores in predicting the up- or downregulated genes in the A or M patient group. Scaled ligand activities are z-score normalized ligand activity values, calculated per receiver cell type. The higher these values, the more enriched the target genes of a specific ligand are compared to other ligands. The blue box indicates the ligands from the type I and type II interferon families. **b)** NicheNet-v2-predicted ligand-target links for the indicated interferon ligands in the cell type "NK cells CD56hi".

### Differential cell-cell communication within healthy lungs versus lungs from patients with idiopathic pulmonary fibrosis (IPF)

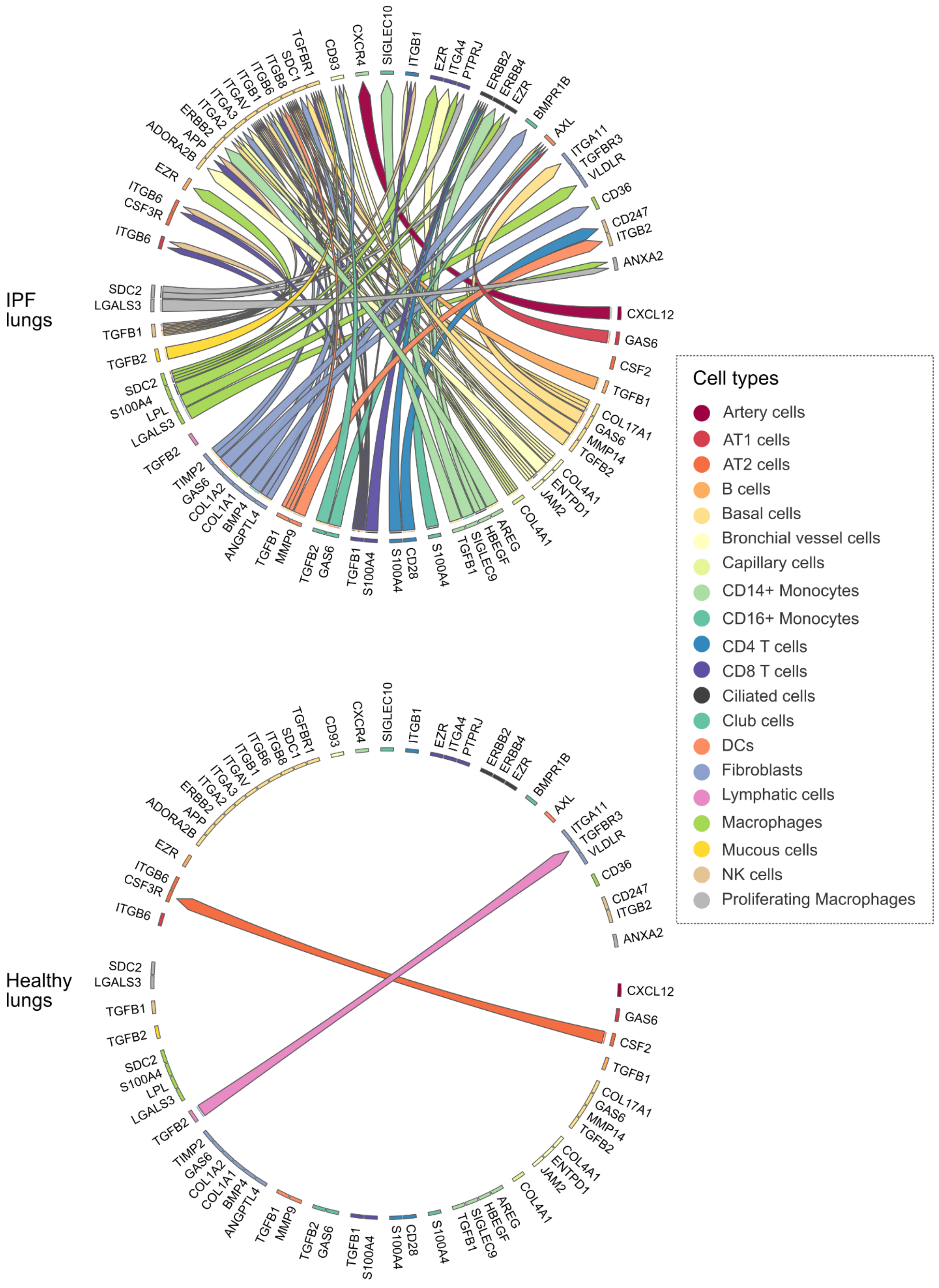


**Supplementary Figure 16 | MultiNicheNet prioritizes differential cell-cell communication patterns within lungs from patients with idiopathic pulmonary fibrosis and healthy patients.** MultiNicheNet was applied to an integrated scRNA-seq lung atlas (see Figure 5) to compare cell-cell communication between healthy lungs and lungs from patients with idiopathic pulmonary fibrosis (IPF). The top 75 differential ligand-receptor pairs are depicted in chord diagrams, divided per patient group. The arrowhead indicates the direction from sender to receiver cell type, and the color of the arrow indicates the sender cell type that expresses the ligand.

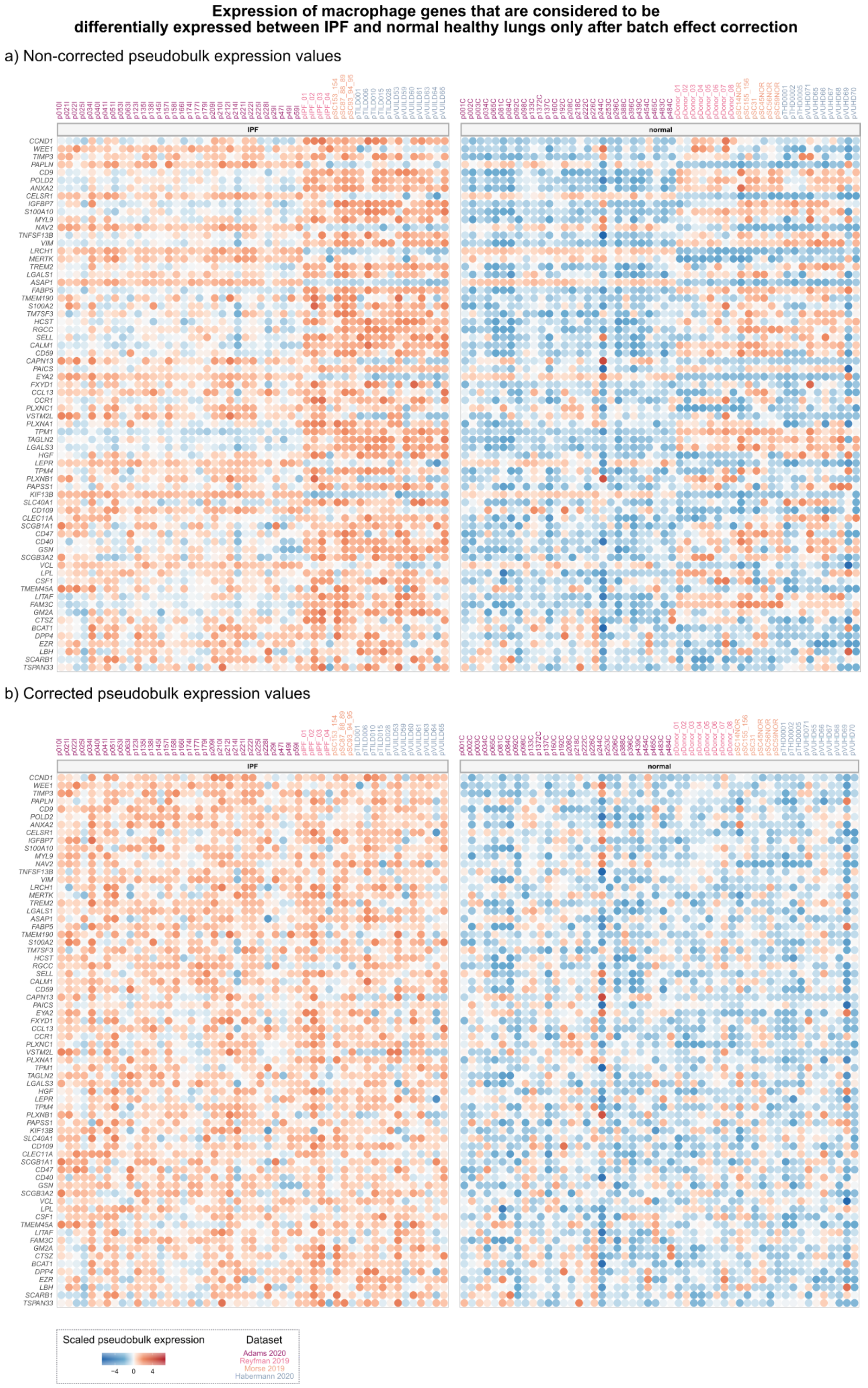


**Supplementary Figure 17 - Impact of batch effect correction on defining differentially expressed genes from integrated scRNA-seq atlas data.** The differential expression (DE) analysis step of MultiNicheNet was performed on an integrated scRNA-seq lung atlas (from four different datasets - see Figure 5) of subjects with healthy lungs and patients with idiopathic pulmonary fibrosis (IPF). DE analysis was once performed with correction for the source dataset, and once without correction. This heatmap shows genes that were significantly (adjusted p-value < 0.05) more strongly expressed in IPF macrophages only after batch correction. **a)** Visualization of non-corrected pseudobulk expression values per patient. b) Visualization of batch-corrected pseudobulk expression values per patient.

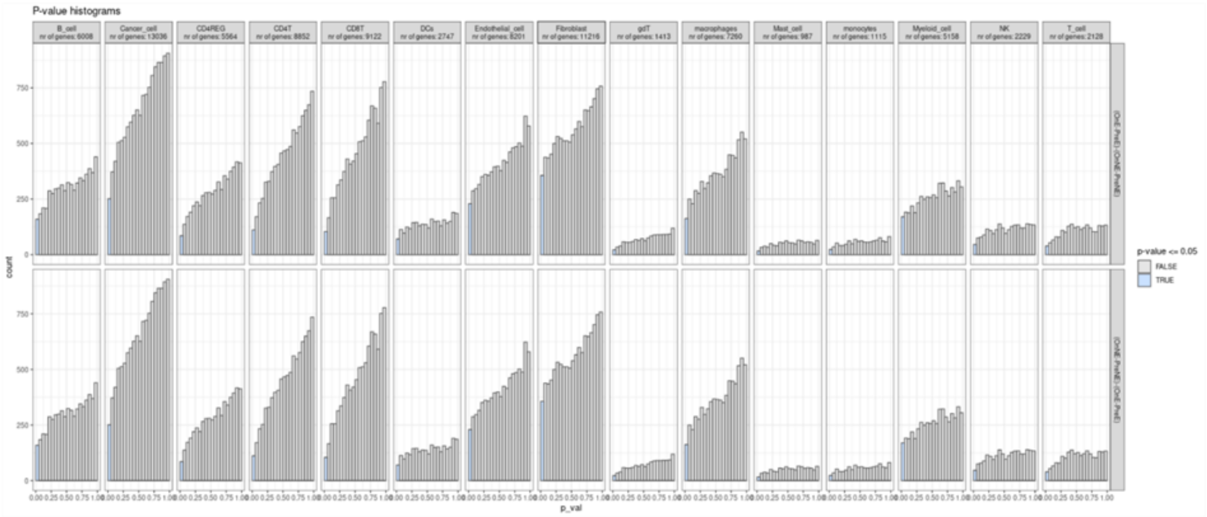


**Supplementary Figure 18 - P-value distributions indicating violation of the DE model assumptions.** MultiNicheNet was applied to scRNA-seq from Bassez et al. to compare on-therapy versus pre-therapy differences in cell-cell communication between the expander and non-expander patients. Each panel indicates the distribution of the DE analysis p-values per contrast-cell-type combination.
