## Supplementary Notes for "MultiNicheNet: a flexible framework for differential cell-cell communication analysis from multi-sample multi-condition single-cell transcriptomics data"

#### Supplementary Note 1: NicheNet-v2 data source collection and processing

##### Ligand-receptor interaction data sources

In NicheNet-v2, we replaced the ligand-receptor network with a novel ligand-receptor network consisting mainly of ligand-receptor interactions from Omnipath and Verschueren et al[1,2]. NicheNet-v1 ligand-receptor interactions[3] coming from curated data sources were also added in the end if they were not part yet of the Omnipath/Verschueren interactions. In the next sections, we will discuss how we processed these data sources to get to the final ligand-receptor network.

*Omnipath ligand-receptor interactions*

The Omnipath intercellular database is a comprehensive database that contains intercellular protein interactions from several databases (e.g., CellPhoneDb[4] and CellChat[5]). We started from this network to construct the NicheNet-v2 ligand-receptor network but we first processed the Omnipath interactions. We considered this to be necessary because we observed some lower confidence interactions and incorrect directionality between the interactions. The latter problem likely arises due to the inclusion of databases from tools for which ligand-receptor directionality does not matter (e.g., CellphoneDB[4]). However, it is crucial for NicheNet that ligand-receptor directionality is correct because we want to predict ligand-regulated target genes in the receptor-expressing receiver cell type.

We applied a two-step processing strategy. First, we used the annotations in the intercellular communication database to get confident annotations of ligands ("transmitters" in Omnipath terminology) and receptors ("receivers" in Omnipath terminology): which genes encode for “transmitter” proteins, and which for “receivers”. Secondly, we queried the Omnipath interaction network for interactions between these confident “transmitters” and “receivers”.

To get the intercellular annotations, we first used the Omnipath function `import_omnipath_intercell` while filtering out annotations only coming from the GO_Intercell or Omnipath source databases. This filter was applied to avoid having proteins like TRAF4, ACTA1, CDK4, and CCND1 being annotated as ligands, and C3 and CCND1 as receptors. We observed that only doing this step leads to missing some ligands, like NAMPT, which are secreted but not annotated as “transmitter” in Omnipath. To include this type of orphan ligands among the entire set of ligands, we used the annotation database to search for proteins that are secreted but not annotated as “transmitter”, and that have a higher Omnipath consensus score to be secreted than intracellular. We also observed that some genes are annotated as both “transmitter” and “receiver” by the same database(s), whereas some genes are annotated as “transmitter” according to some databases and as “receiver” according to other databases. Because we want to avoid that some transmitters are also described as “receiver” in the network if the receiver annotation was coming from a minority of data sources, we did some further filtering. For these latter genes, we only kept their “transmitter” annotation if the consensus score for being a transmitter was more than 1 score higher than for being a “receiver” (and vice versa).

So to conclude the previous steps, we considered as possible ligands the following proteins: 1) transmitter-only proteins, 2) proteins with more evidence of being a transmitter, 3) proteins with evidence of being both a transmitter and receiver, and 4) the orphan ligands. As possible receptors we considered: 1) receiver-only proteins, 2) proteins with more evidence of being a receiver, 3) proteins with evidence of being both a transmitter and receiver.

Manual inspection of these annotations indicated that some well-known ligands and receptors were still missing (e.g., CXCL16, DPP4, and IFNA13). Therefore we added ligands and receptors from the comprehensive Omnipath ligand-receptor network directly through the function `import_intercell_network(omnipath = TRUE, ligrecextra = TRUE)`. Because the Omnipath ligand-receptor network contains some unlikely interactions, we applied further filtering steps. First, we required that the category of the ligand should be one of: "ligand", "cell_adhesion", "cell_surface_ligand", "secreted", "interleukins_hgnc","chemokine_ligands_hgnc", "endogenous_ligands_hgnc", "surface_ligand", "secreted_enzyme"; and the category of the receptor should be one of: "receptor", "cell_adhesion", "interleukin_receptors_hgnc". Second, we removed interactions only described in the Wang data source because this data source was at the basis of many unlikely interactions we retrieved. Thirdly, we removed interactions that were only documented in the following databases or database combinations: “Omnipath", "LRdb;OmniPath", "GO_Intercell;OmniPath", "scConnect;OmniPath", "Cellinker;OmniPath", "Cellinker;scConnect;OmniPath", "Cellinker;Zhong2015;OmniPath”. Not performing these filtering steps would include non-ligands such as JAK2, NOTCH1, PIK3CA, and TP53 as ligands; and non-receptors such as JAK2, CX3CL1, and CXCL16 as receptors. Next, we used the remaining ligand-receptor network to extract the ligands, receptors, and proteins that can be both in the same way as described above.

Finally, we removed ligands if they were more likely to be intracellular than secreted or membrane-bound (according to Omnipath’s locational consensus scores). This led to the correct removal of proteins like ARF1, IRF8, MYH9, and SOCS2 as ligands. Furthermore, we added some extra ligands that were not included before but have “ligand activity” as gene ontology (GO) annotation (examples: FGF11, FGF12, and FGF14).

After having defined the list of potential ligands and receptors, we queried the Omnipath network (“Omnipath”, “ligrecextra” and “pathwayextra” datasets – with Wang resource excluded) to search for interactions between these defined ligands and receptors. Because some popularity-biased proteins have many interactions, we filtered out interactions for which the curation score was less than 2.5% of the total curation score for that ligand/receptor. For example, this filtering enables keeping TNF- TNFRSF1A/B and not TNF-NOTCH1.

*Verschueren et al. ligand-receptor interactions*

Verschueren et al. documented several novel potential cell surface interactions involving at least one member of the immunoglobulin superfamily (IgSF)[2]. We retrieved these novel interaction pairs from Supplementary Table 2. To get the correct ligand-receptor directionality, we post-processed this table by considering the ligand and receptor annotations from the processed Omnipath-derived ligand-receptor network. Next, we split up the interactions into a network containing the unique interactions, and a network containing interactions that are also documented in the processed Omnipath-derived network. Finally, we combined the unique, non-Omnipath, interactions from Verschueren and curated NicheNet-v1 databases into a separate data source.

#### Signaling interaction data sources

To construct NicheNet-v2's signaling network, we first updated the signaling and protein-protein interactions from Omnipath[6] (regular version, not intercellular) and PathwayCommons[7]. Interactions from Omnipath were now retrieved via the OmnipathR command `import_post_translational_interactions`, and split up into directional and non-directional interactions[6]. For Pathwaycommons, we downloaded network version 12[7].

Furthermore, we added two new data sources: HuRi[8] and HumanNet[9]. The HuRi database provides three types of networks: HuRI, HI-union, and Lit-BM. Each of those was downloaded (http://www.interactome-atlas.org/download) and included as a separate data source. For the HumanNet signaling protein-protein interactions we downloaded the HS-PI file (https://www.inetbio.org/humannet/download.php).

#### Gene regulatory interaction data sources

Similarly as for the signaling interactions, we updated the gene regulatory interactions from PathwayCommons to version 12[7]. We also updated the transcription factor (TF) - target links that we derived from the ReMap database of ChIP-seq profiles. First, we used the updated ReMap 2022 database[10]. To process this database into TF-target interactions, we used the same procedure as described in the original NicheNet paper[3]. However, we now used a model of regulatory potential that takes into account the regulatory range of a TF, as described by Chen et al.[11]

Furthermore, we added interactions based on TF-perturbation experiments from KnockTF v1[12]. This database provides differential expression (DE) profiles after TF perturbation. To handle the different ranges of log fold changes and p-values across the different datasets, we z-score normalized the logFC values per dataset, and considered genes a target of a TF when the absolute value of the z-score normalized value was >= 2.5.

We also added interactions from the comprehensive database Dorothea[13] (https://github.com/saezlab/dorothea/tree/master/data). We divided Dorothea's interactions into two sub-data sources: one high-confident data source with confidence levels A, B, and C; and one lower-confident with interactions with confidence level D. This was done for the mouse and human interactions that were provided.

Next, we collected data sources that provide information about potential direct ligand-target links, given their importance for NicheNet's ligand-target predictions[3]. First, we added gene regulatory interactions from the HumanNet co-expression network ("HS-CX")[9]. We only kept links with ligands as regulators (as the definition of ligands, we considered all from/source nodes of the ligand-receptor network). As a result, we added ligand-target links for which the ligand and the target are correlated in expression across tissues.

Finally, we also added direct ligand-target links from ligand treatment datasets from NicheNet[3] and CytoSig[14]. Ligand treatment datasets are datasets where cells are *in vitro* treated with a ligand and the transcriptome is measured before and after ligand stimulation. DE genes after ligand stimulation can then be considered as target genes of the ligand. By including links from this type of data, we incorporated *in vitro* experimentally defined links in the model.

For the processing of the NicheNet ligand treatment datasets, we only retained datasets where one ligand was added. Genes were considered as a target when $|logFC| > 1$ and adjusted p-value $<= 0.10$ (which is the same definition used in the original NicheNet study to define gold standard target genes[3]). Because the ligand treatment dataset collection contains multiple datasets for some ligands, we were able to divide interactions into frequent versus infrequent interactions. If a gene was a target in only one dataset, it was considered as "infrequent" target. If it was DE in more than one dataset, it was considered as "frequent" target. Infrequent and frequent ligand-target were each split into different data source: we hypothesize that both types of link might have different degrees of trustworthiness, which could be of importance during downstream data source weight optimization.

The CytoSig database provides two types of data files: the "signatures" file and the "per-dataset-file". The "signatures file" documents the median logFC of the gene after ligand treatment across all datasets. These signatures are only provided for the subset of ligands for which the *in vitro* response signature is representative of human physiological responses (determined through across-tissue correlation analysis by the authors[14]). To handle the different ranges of log fold changes and p-values across the different ligand signatures, we z-score normalized the logFC values per signature, and considered genes a target of a ligand when the absolute value of the z-score normalized value was >= 2.5. We compiled these ligand-target interactions into the so-called "Cytosig-signature" data source. The "per-dataset-file" documents the logFC of the gene after ligand treatment per dataset, for all considered ligands. To handle the different ranges of log fold changes and p-values across the different ligand signatures, we z-score normalized the logFC values per dataset, and considered genes a target of a ligand in that specific dataset when the absolute value of the z-score normalized value was >= 2.5. Then we divided target genes per ligand into "frequent" and "less frequent" targets based on whether a gene was a target in respectively more than 20% of datasets of that ligand or not. We only considered frequent targets as target genes of a ligand, and grouped these ligand-target links into the "Cytosig_all" data source.

#### Conversion of gene identifiers

Most data sources provided interactions for human and a minority for mouse. To combine these in one final model (in HGNC human gene symbols or MGI mouse gene symbols), an interaction between mouse genes was converted to an interaction between their human orthologs and vice versa (mainly from NCBI HomoloGene via annotationTools::getHOMOLOG, additional orthology relations were retrieved from ENSEMBL via biomaRt). For all data sources, interactions were only kept if both genes participating in an interaction were annotated in the database of the org.Hs.eg.db R package. This database was also used to map Entrez ID, Uniprot and ENSEMBL identifiers to gene symbols if the original network did not provide interactions in gene symbols. Conversion of gene symbol aliases to the newest version of gene symbols was performed through org.Hs.eg.db and org.Mm.eg.db (entrez gene source date: EGSOURCEDATE: 2021-Apr14; NCBI Gene Database).

#### Data source analysis

An overview of all the databases used to construct the NicheNet-v2 model, and their division into data sources, can be found in **Supplementary Table 1a-c**. We also calculated network properties such as the number of genes and the number of interactions, which we documented for the individual source and final integrated networks (**Supplementary Table 1b and 1d)**. An analysis of the overlap between databases/data sources is presented in **Supplementary Notes Figure 1.1.-1.4.** The relation between the number of interactions and the number of data sources supporting an interaction is presented in **Supplementary Notes Figure 1.5.** These analyses indicate that the collection of multiple complementary data sources resulted in substantially higher coverage, as was the objective. Since many different databases retrieve interactions from the same experiments, redundancy between some data sources could be observed as well.

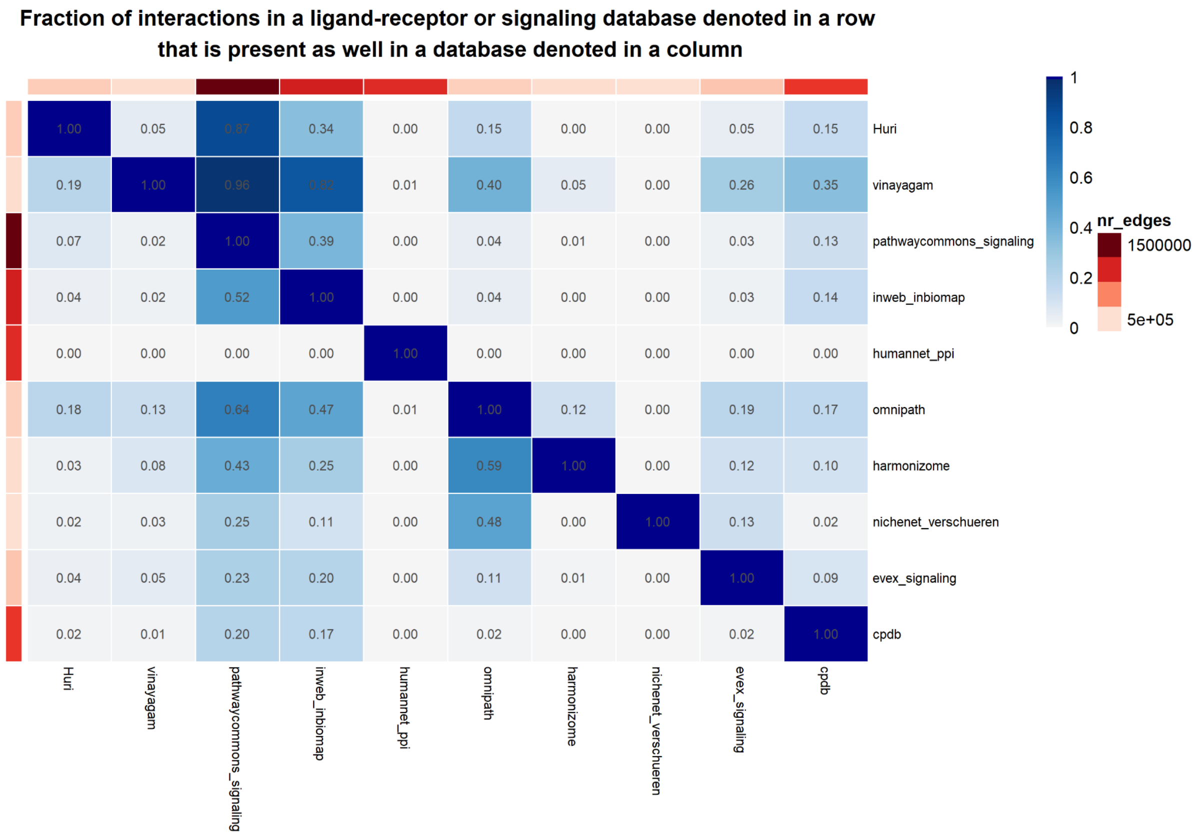


**Supplementary Notes Figure 1.1. | Overlap in interactions between the databases providing ligand-receptor and signaling interactions.** The blue color scale indicates the fraction of interactions in a row-denoted database that are also present as well in the column-denoted database.

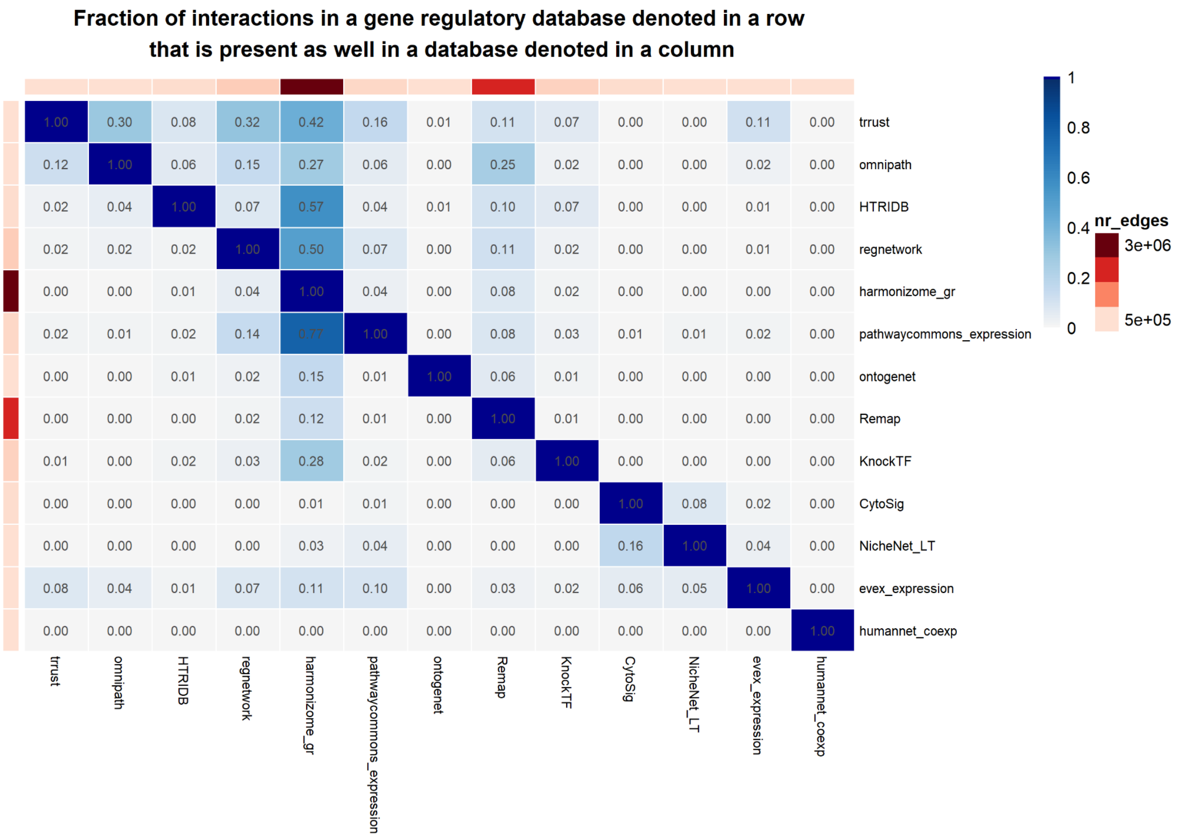


**Supplementary Notes Figure 1.2. | Overlap in interactions between the databases providing gene regulatory interactions.** The blue color scale indicates the fraction of interactions in a row-denoted database that are also present as well in the column-denoted database.

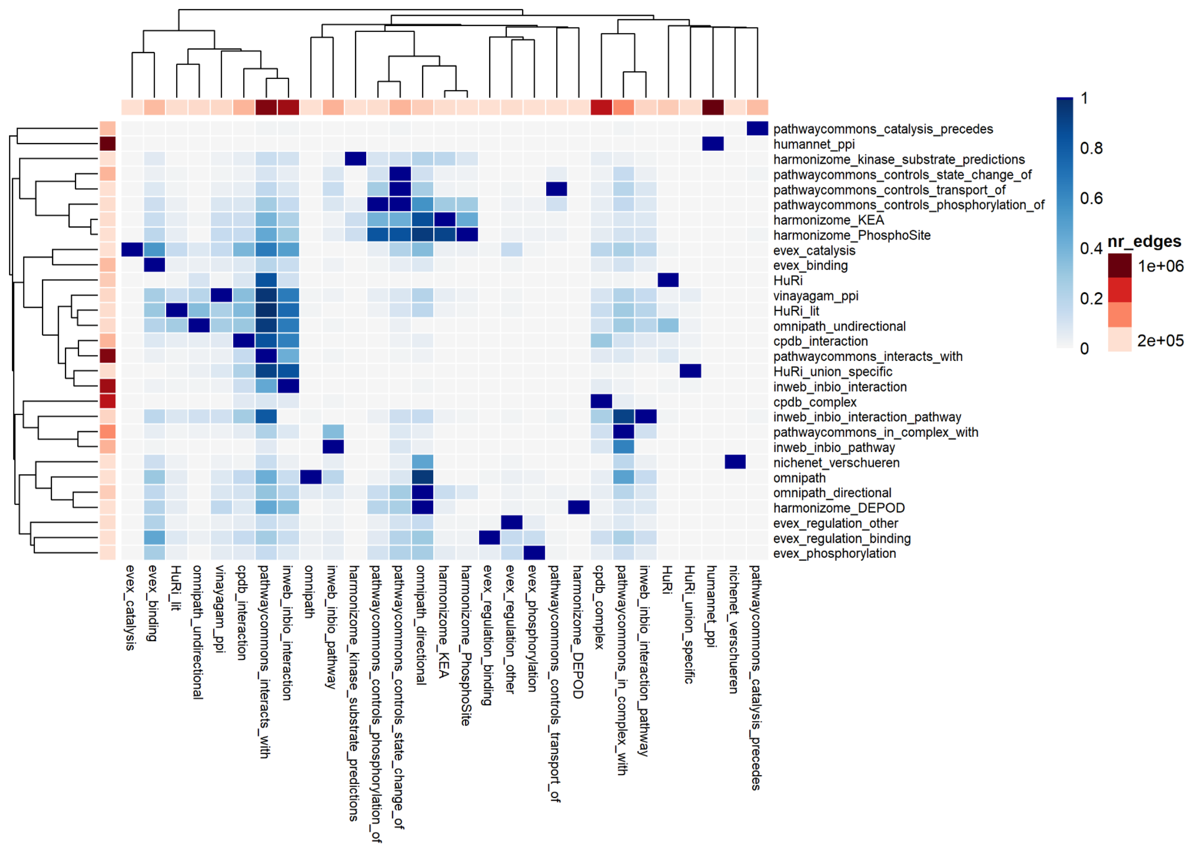


**Supplementary Notes Figure 1.3. | Overlap in interactions between the data sources providing ligand-receptor and signaling interactions.** The blue color scale indicates the fraction of interactions in a row-denoted data source that are also present as well in the column-denoted data source.

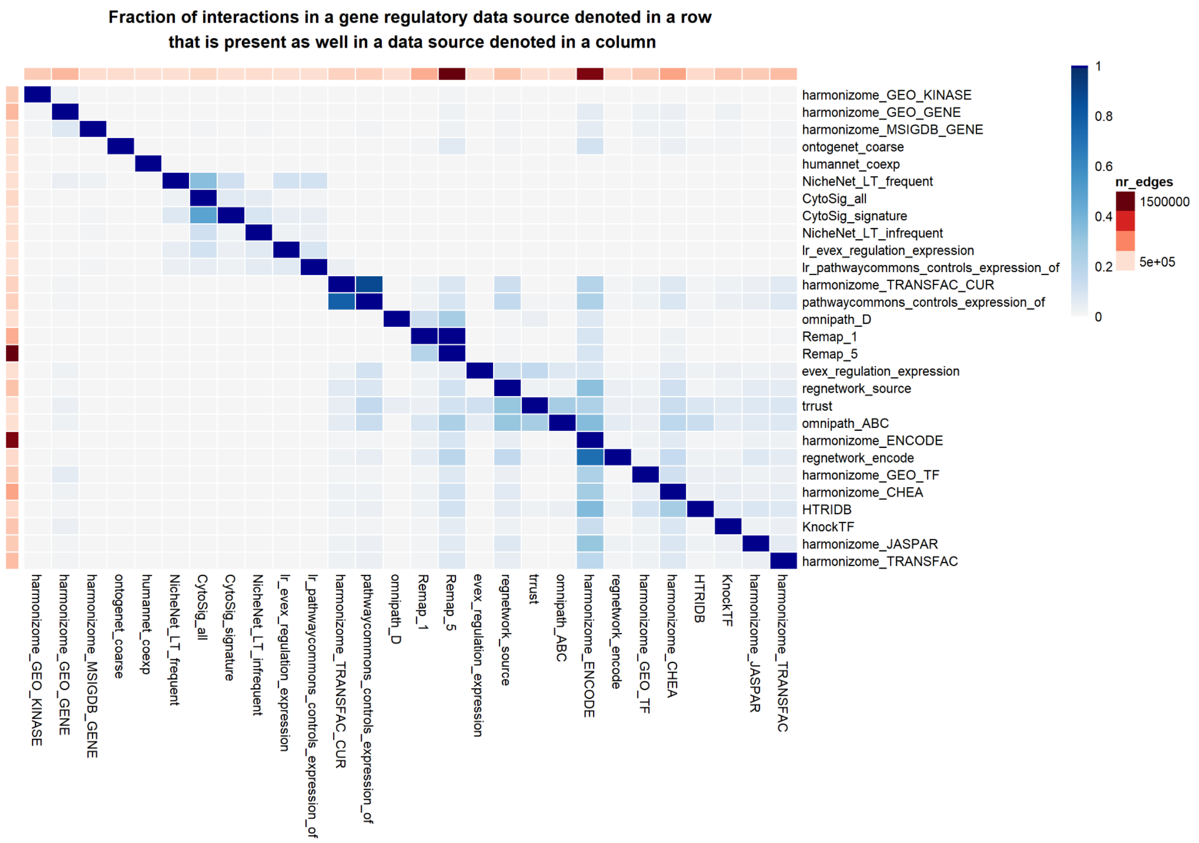


**Supplementary Notes Figure 1.4. | Overlap in interactions between the data sources providing gene regulatory interactions.** The blue color scale indicates the fraction of interactions in a row-denoted data source that are also present as well in the column-denoted data source.

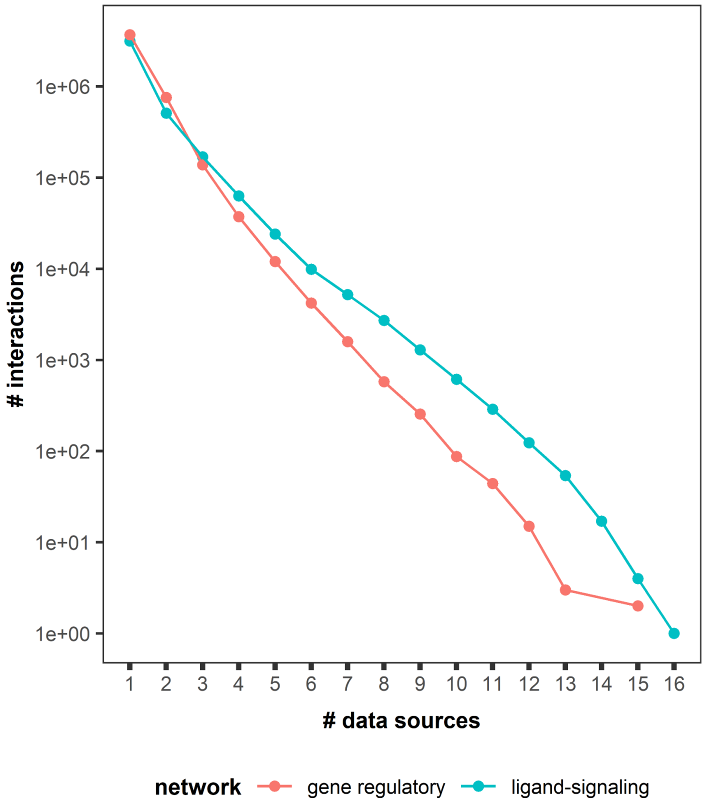


**Supplementary Notes Figure 1.5. | The majority of interactions are documented by only one or a few data sources.** The number of interactions (on a log scale) is plotted versus the number of data sources that support these interactions. In total, there are 28 gene regulatory data sources and 29 ligand signaling data sources.

### Supplementary Note 2: results of NicheNet-v2 parameter optimization

As described in the **Methods** section, multi-objective optimization with NSGA-II[15] was performed in a fivefold cross-validation-like scheme to optimize the NicheNet-v2 data source weights and hyperparameters. This supplementary note shows the results of the optimization process.

**Supplementary Notes Figure 2.1.** shows the values of the optimization criteria (target gene and ligand activity prediction performance) for all models evaluated during the optimization process. The depicted graph shows the results for one fold in the cross-validation-like procedure (namely 'fold1234', see **Supplementary Table 2b**). Results of the other folds can be observed in the Zenodo repository of all code and data required to reproduce the results of this study (https://zenodo.org/record/8016880). The main conclusions of this figure are that parameter values can have a substantial influence on performance and that optimization improves performance.

**Supplementary Notes Figure 2.2.** visualizes the evolution of the optimization criteria values over all the optimization rounds (again for one fold in the cross-validation-like procedure, namely 'fold1234'). This indicates that optimization over multiple rounds indeed resulted in higher optimization criteria values.

These results indicate that, within the training set, the optimization procedure indeed results in higher performance. However, we also wanted to assess whether optimization resulted in better model performance when performance is assessed on the independent test datasets (see **Methods**). To determine this, we constructed four different 'optimized models': 1) model using the mean parameter values of the 5 parameter settings with the highest geometric average over the four objectives per fold; 2) model using the median parameter values of the 5 parameter settings with the highest geometric average over the four objectives per fold; 3) model using the mean parameter values of the 25 parameter settings with the highest geometric average over the four objectives per fold; 4) model using the median parameter values of the 25 parameter settings with the highest geometric average over the four objectives per fold. These models were then compared to one example of an unoptimized model with neutral data source weights and default hyperparameter values (see **Methods**). Target gene prediction performance was better for the optimized models compared to the unoptimized model (**Supplementary Notes Figure 2.3.)** but ligand activity prediction performance was similar (**Supplementary Notes Figure 2.4.)**. The performance of the four different optimized models was nearly identical (**Supplementary Notes Figure 2.3.** and **2.4.)**. Note that the performance of this unoptimized model is relatively high, probably because the default hyperparameter parameter values are quite comparable to the final optimized values. This is in contrast to the weakly performing models in **Supplementary Notes Figure 2.1.** and **2.2.**

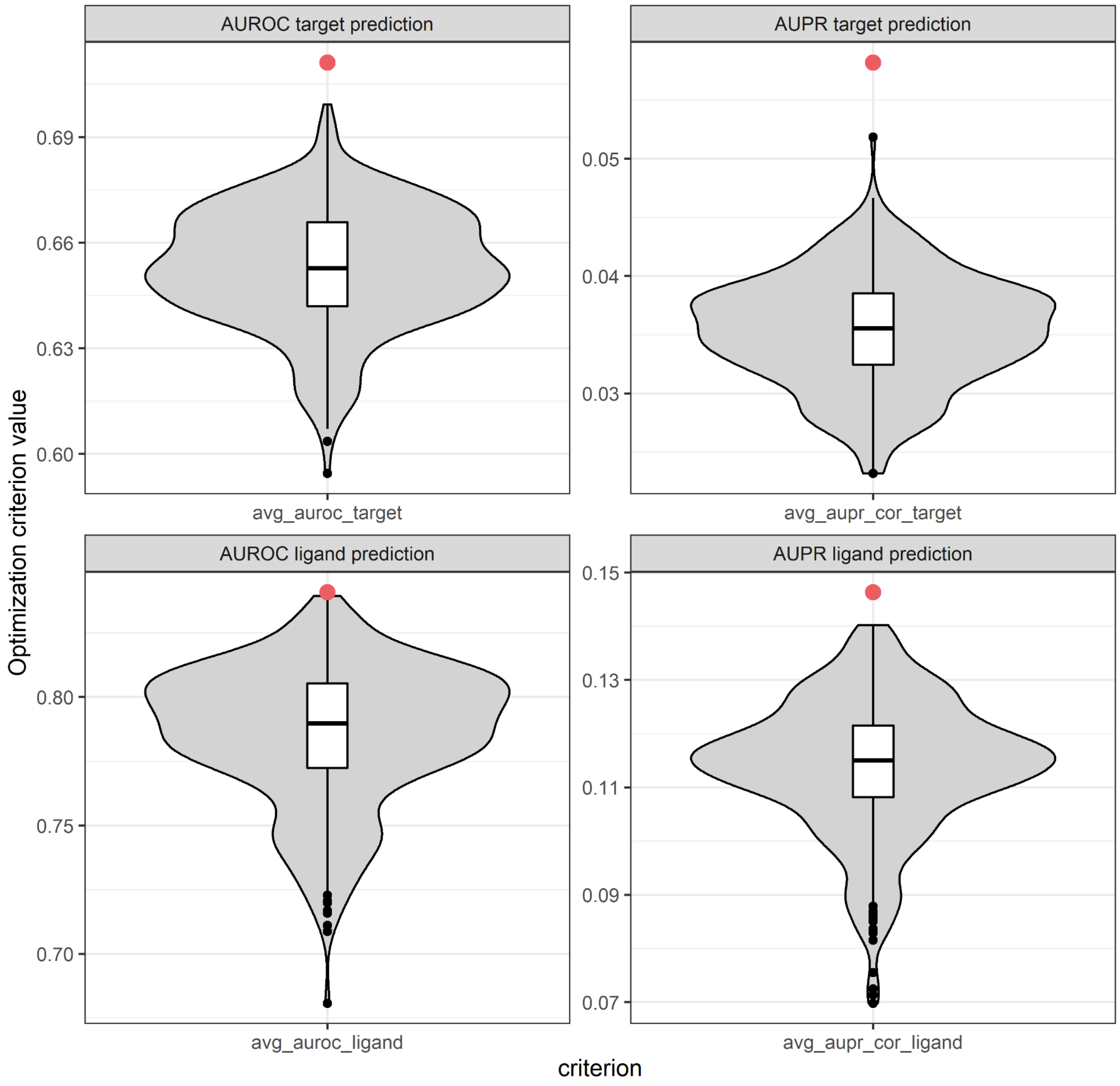


**Supplementary Notes Figure 2.1. | Target gene and ligand activity prediction performances of all models generated during the optimization process - example for fold f1234.** To assess the effect of different data source weights and hyperparameter values on the model's performance, we show median ligand prediction and target prediction performances over 131 NicheNet and CytoSig ligand treatment datasets of all 5400 models evaluated during the NSGA-II multi-objective optimization procedure. The red dot indicates the performance of the final model with the most optimal parameter settings. AUPR (corrected): area under the precision-recall curve, corrected for random prediction; AUROC: area under the receiver-operating-characteristic curve.

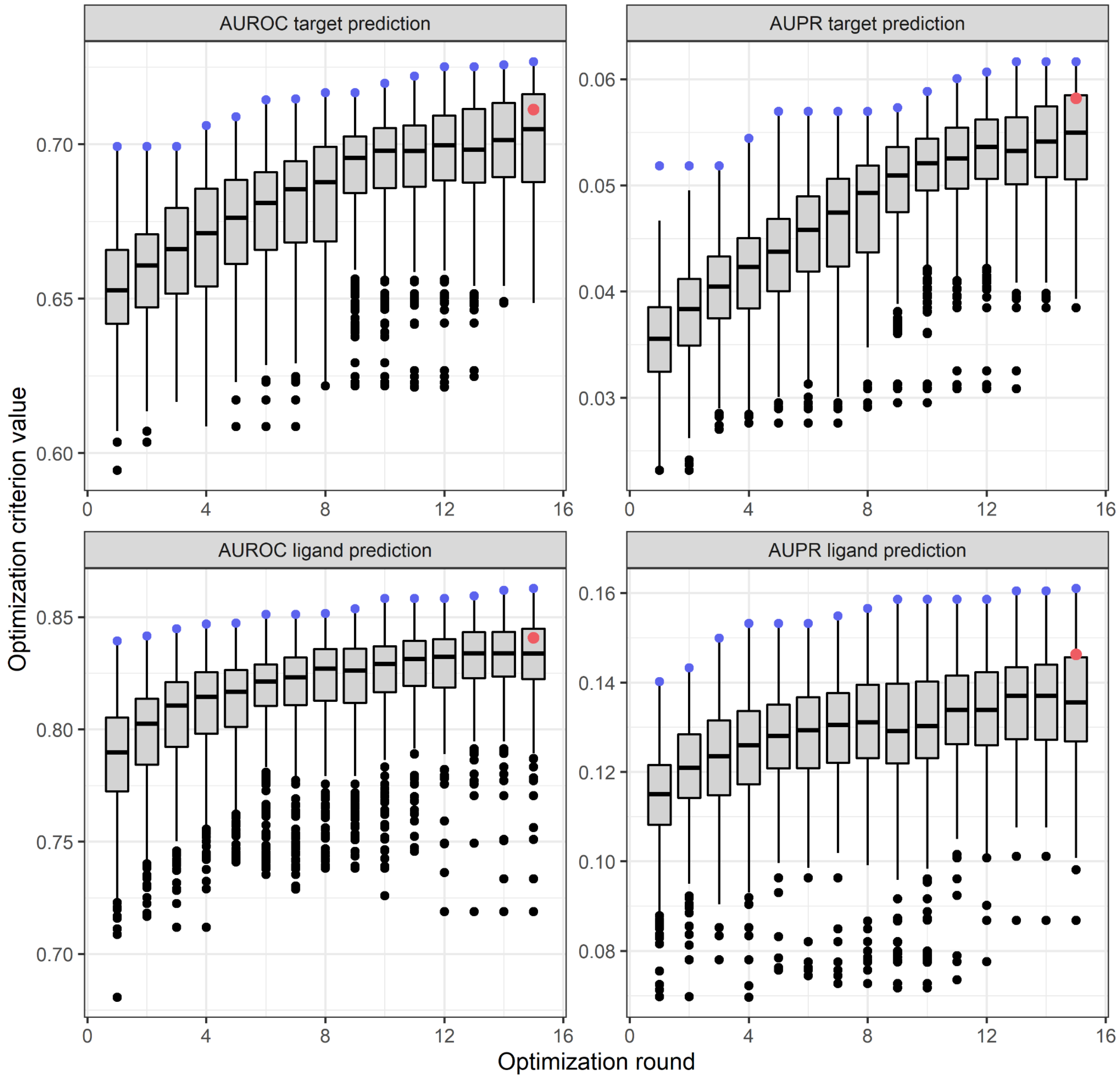


**Supplementary Notes Figure 2.2. | Evolution of the optimization criteria target gene and ligand activity prediction performances during the optimization process - example for fold f1234.** For all models evaluated in an individual optimization round during the NSGA-II multi-objective optimization procedure, boxplots are shown of the median ligand prediction and target prediction performances over 131 NicheNet and CytoSig ligand treatment datasets. Boxplots summarize results for 360 models in every optimization round. The red dot indicates the performance of the final model with the most optimal parameter setting. AUPR (corrected): area under the precision-recall curve, corrected for random prediction; AUROC: area under the receiver-operating-characteristic curve.

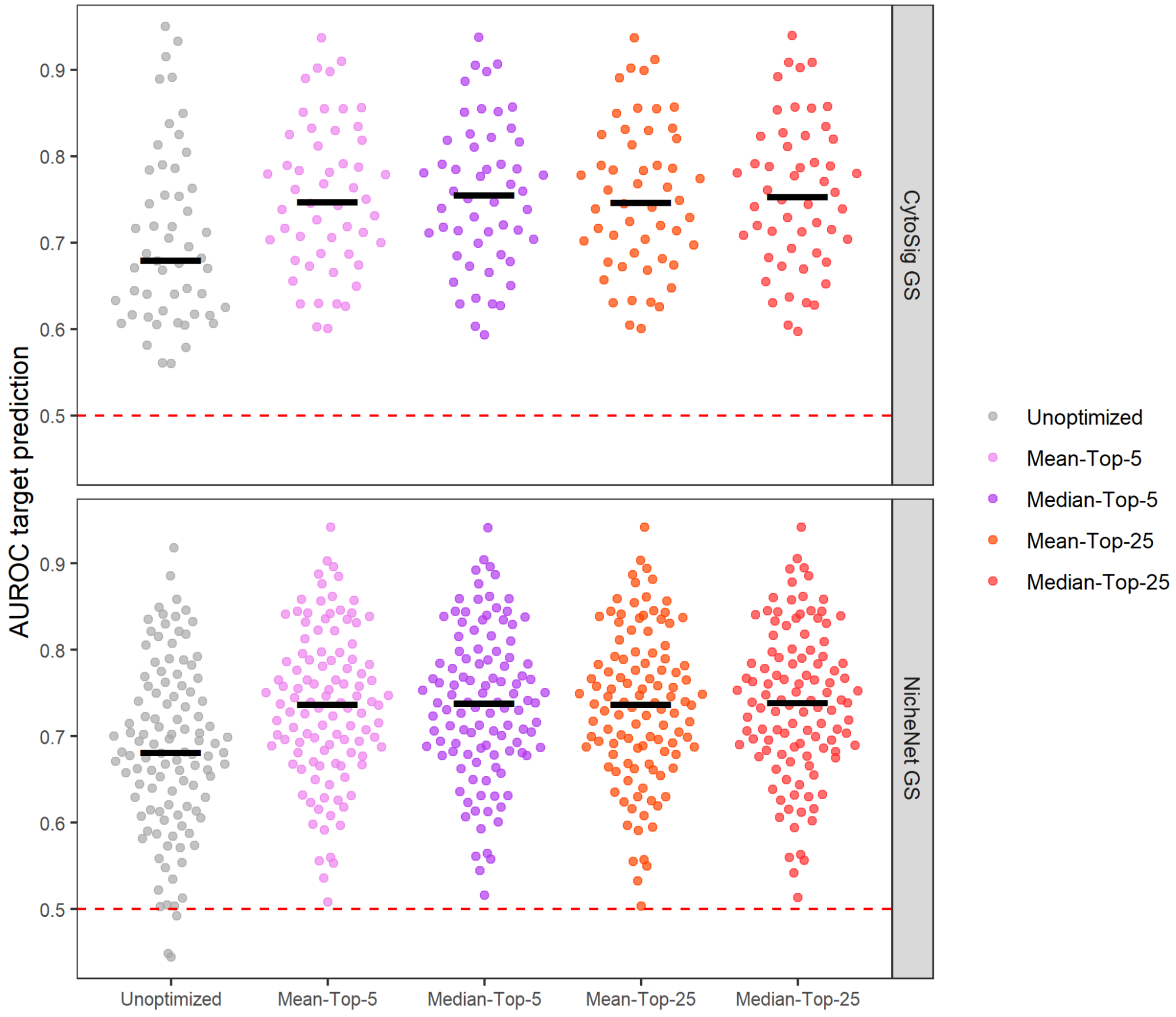


**Supplementary Notes Figure 2.3. | Target gene prediction performance: difference between an unoptimized model and optimized models.** For 52 CytoSig and 111 NicheNet ligand treatment datasets providing transcriptome data of cells after *in vitro* ligand treatment, we used a fivefold cross-validation-like strategy to assess performance in predicting which genes are differentially expressed on treatment with a particular ligand. We compared an unoptimized model to four different optimized models that were constructed by using the mean or median parameter values of the top 5 or 25 parameter settings of each of the five folds. Each dot indicates the performance for one dataset, and the black line indicates the median performance over all datasets. The red dashed line indicates the performance of random guessing. AUROC: area under the receiver-operating-characteristic curve.

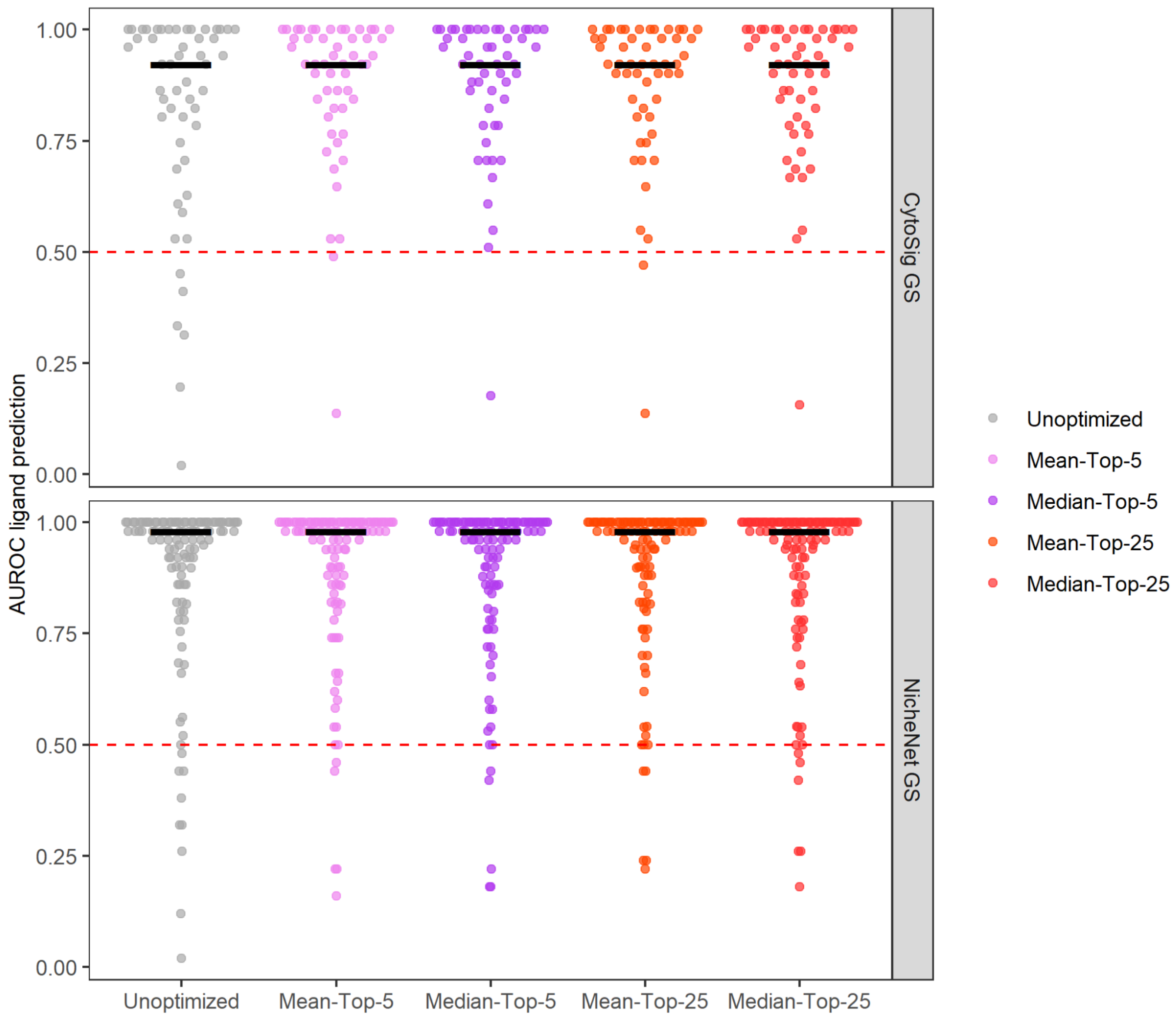


**Supplementary Notes Figure 2.4. | Ligand activity prediction performance: difference between an unoptimized model and optimized models.** For 52 CytoSig and 111 NicheNet ligand treatment datasets providing transcriptome data of cells after *in vitro* ligand treatment, we used a fivefold cross-validation-like strategy to assess performance in predicting the ligand by which the cells were treated based on the differentially expressed genes after treatment. We compared an unoptimized model to four different optimized models that were constructed by using the mean or median parameter values of the top 5 or 25 parameter settings of each of the five folds. Each dot indicates the performance for one dataset, and the black line indicates the median performance over all datasets. The red dashed line indicates the performance of random guessing. AUROC: area under the receiver-operating-characteristic curve.

### Supplementary Note 3: MultiNicheNet highlights critical pre-therapy cell-cell signaling patterns linked to therapy response in breast cancer patients

We applied MultiNicheNet to scRNA-seq data collected from breast cancer patients before and during anti-PD1 therapy[16]. We compared pre-treatment cell-cell communication between patients with T cell clonotype expansion during therapy (E, or expander patients) and patients with limited or no expansion (NE or non-expander patients). In this supplementary note, we will investigate interactions between all cell type pairs in the tumor microenvironment. The most strongly differential interactions are depicted in **Supplementary Notes Figure 3.1.** (circular visualization of top 50), **Supplementary Notes Figure 3.2.-3.3.** (bubble heatmaps of top 100), and **Supplementary Tables 3a-b** (top 1000). One of the most highly active and differential interactions in E patients involves IL-21 from CD4 T cells to CD4 regulatory T cells (**Supplementary Notes Figures 3.1.-3.2.**), which has been shown to counteract the immune-suppressive activities of regulatory T cells[17]. Other interesting E-specific interactions include increased CSF2 receptor signaling toward dendritic cells (DCs) and a SEMA4D-PLXNB2 interaction between natural killer (NK) cells and DCs (**Supplementary Notes Figure 3.2.**). Previous research described their role in modulating the activity of DCs[18,19]. Regarding NK cells in E patients, they seem to receive MIF from malignant cancer cells, which may inhibit NK cell-mediated cytotoxicity[20], and the chemokines CCL19 and CCL11 from DCs and fibroblasts (**Supplementary Notes Figure 3.2.**). Both chemokines putatively bind CXCR3, which is involved in NK cell accumulation in tumors[21]. MultiNicheNet also predicted fibroblast - NK cell interactions specific for NE patients. Their predominant interaction is THBS4-CD47 (**Supplementary Notes Figures 3.1.** and **3.3.**). Remarkably, the interaction between THBS4's family member THBS1 and CD47 inhibits NK cell proliferation[22]. Another predicted inhibitory interaction in NE patients is the immune checkpoint interaction SELL-SELPLG between CD4 regulatory T cells and CD8 T cells[23] (**Supplementary Notes Figures 3.1.** and **3.3.**).

Because of strong inter-patient heterogeneity between malignant cells, most ligand-receptor pairs involving malignant cells are less strongly DE. This makes them less likely to be part of the overall top 50 or 100 differential interactions. However, we can specifically zoom in on their interactions, which we did for interactions with T cells (**Supplementary Figures 9-11**). Interactions from T cells to malignant cells in E patients are mainly characterized by TNF, IFNG, and semaphorin-plexin interactions (**Supplementary Notes Figure 3.4.**). E-specific interactions from malignant cells to T cells primarily play a role in antigen presentation (HLA-DRB1/HLA-DRB5/HLA-DRA/HLA-DBQ1 - LAG3/CD82 and B2M-TAP1) (**Supplementary Notes Figure 3.5.**). This implies that malignant cells from NE patients show lower levels of HLA molecules involved in antigen presentation. Just like lower PD-L1 and IFNG signaling, loss of antigen presentation in cancer cells is known to be one of the major determinants of non-responsiveness to immune checkpoint blockade therapy[24]. In addition, MultiNicheNet also reveals some immune checkpoint interactions like PVR-TIGIT (between malignant and CD4 and CD8 T cells) and NECTIN2-TIGIT (between malignant and CD8 T cells) in E patients (**Supplementary Notes Figure 3.5.**). On the contrary, several NE-specific interactions between malignant and T cells are interactions between ephrin ligands and Eph receptors (**Supplementary**

**Notes Figure 3.6.**). Concordant with this finding is the recently described role of Ephs in suppressing immune responses in the tumor microenvironment[25].

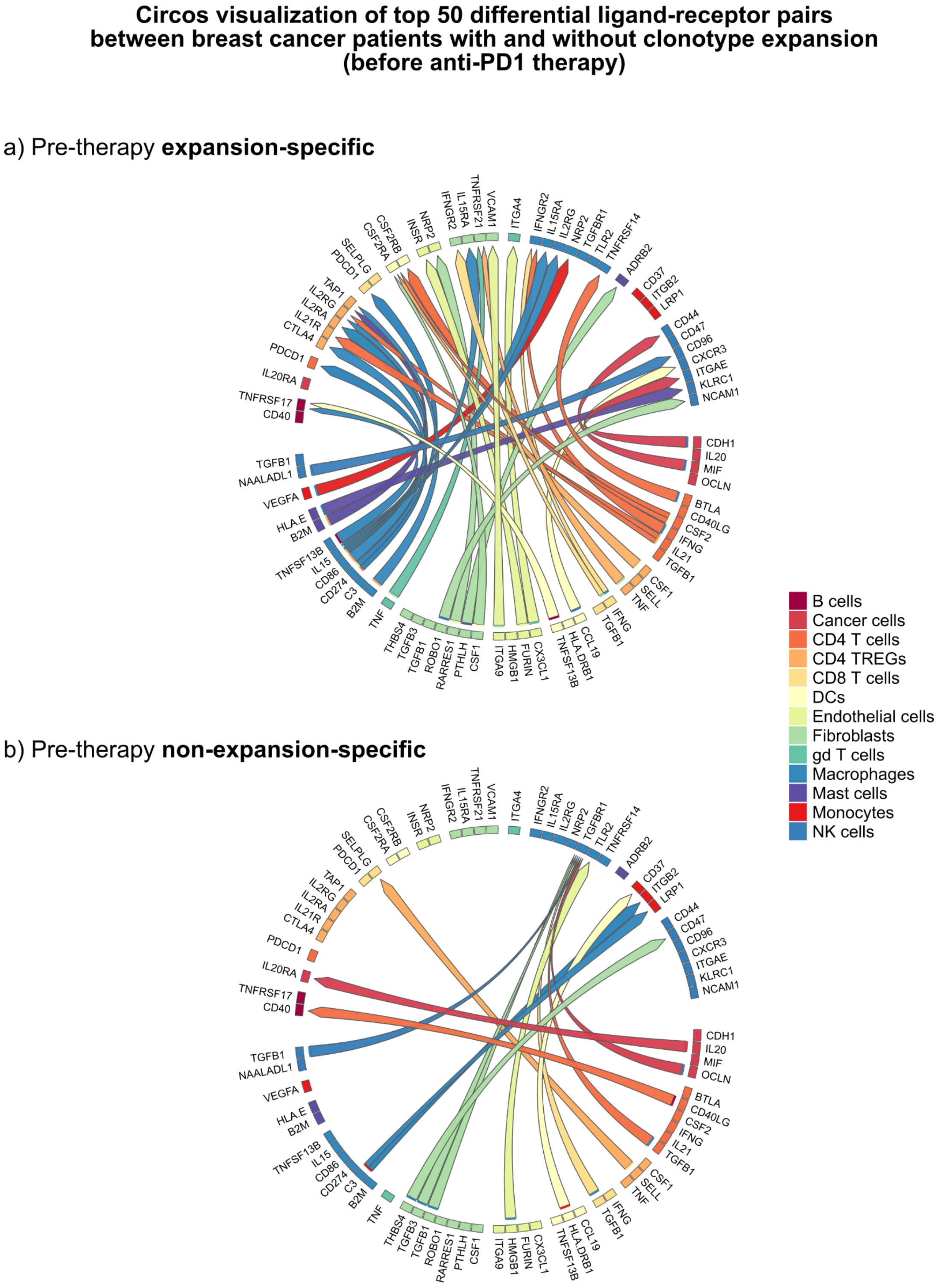


**Supplementary Notes Figure 3.1. | MultiNicheNet prioritizes differential cell-cell communication patterns between patients with and without T cell clonotype expansion.** MultiNicheNet was applied to scRNA-seq from Bassez et al. to compare pre-therapy cell-cell communication between expander and non-expander patients. The top 50 differential ligand-receptor pairs are depicted in chord diagrams, divided into the expander-specific pairs in a) and non-expander-specific pairs in b). The arrowhead indicates the direction from sender to receiver cell type, and the color of the arrow indicates the sender cell type that expresses the ligand.

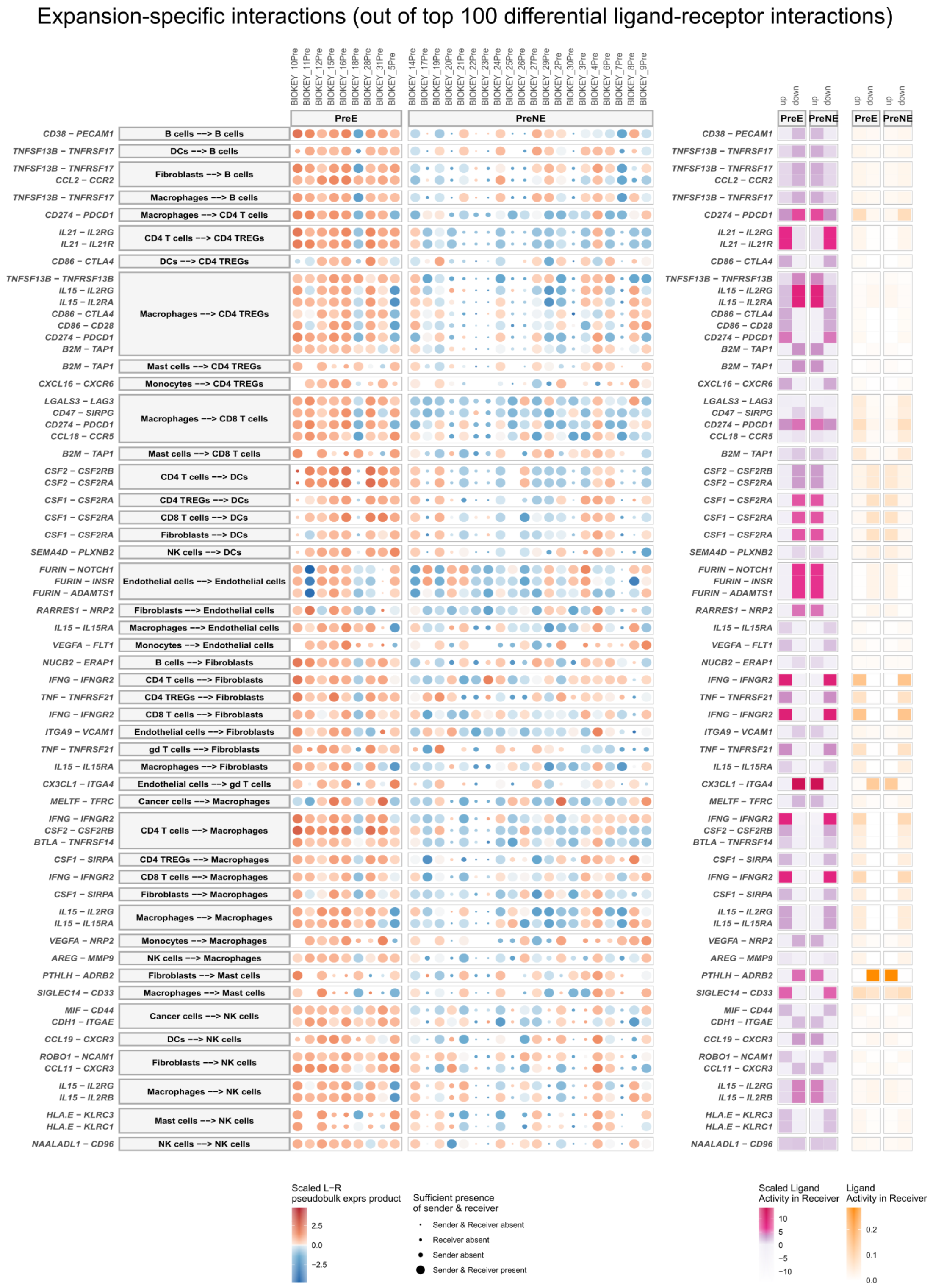


**Supplementary Notes Figure 3.2. | MultiNicheNet prioritizes cell-cell communication patterns specific for patients with clonotype expansion after anti-PD1 therapy.** MultiNicheNet was applied to scRNA-seq from Bassez et al. to compare pre-therapy cell-cell communication between expander (preE) and non-expander patients (preNE). The preE-specific interactions out of the total top 100 differential interactions are shown. For each interaction, ligand-receptor pseudobulk expression (product of logCPM values) and (scaled) ligand activity values are visualized. Ligand activity scores are the AUPRC scores in predicting the up- or downregulated genes in the preE or preNE group. Scaled ligand activities are z-score normalized ligand activity values, calculated per receiver cell type. The higher these values, the more enriched target genes of a specific ligand are compared to other ligands. The size of the dots indicates whether a sample had enough cells (>= 10) for a specific cell type to be considered for DE analysis. The absence of a dot means there were no cells at all. L-R: ligand-receptor.

Non-expansion-specific interactions (out of top 100 differential ligand-receptor interactions)

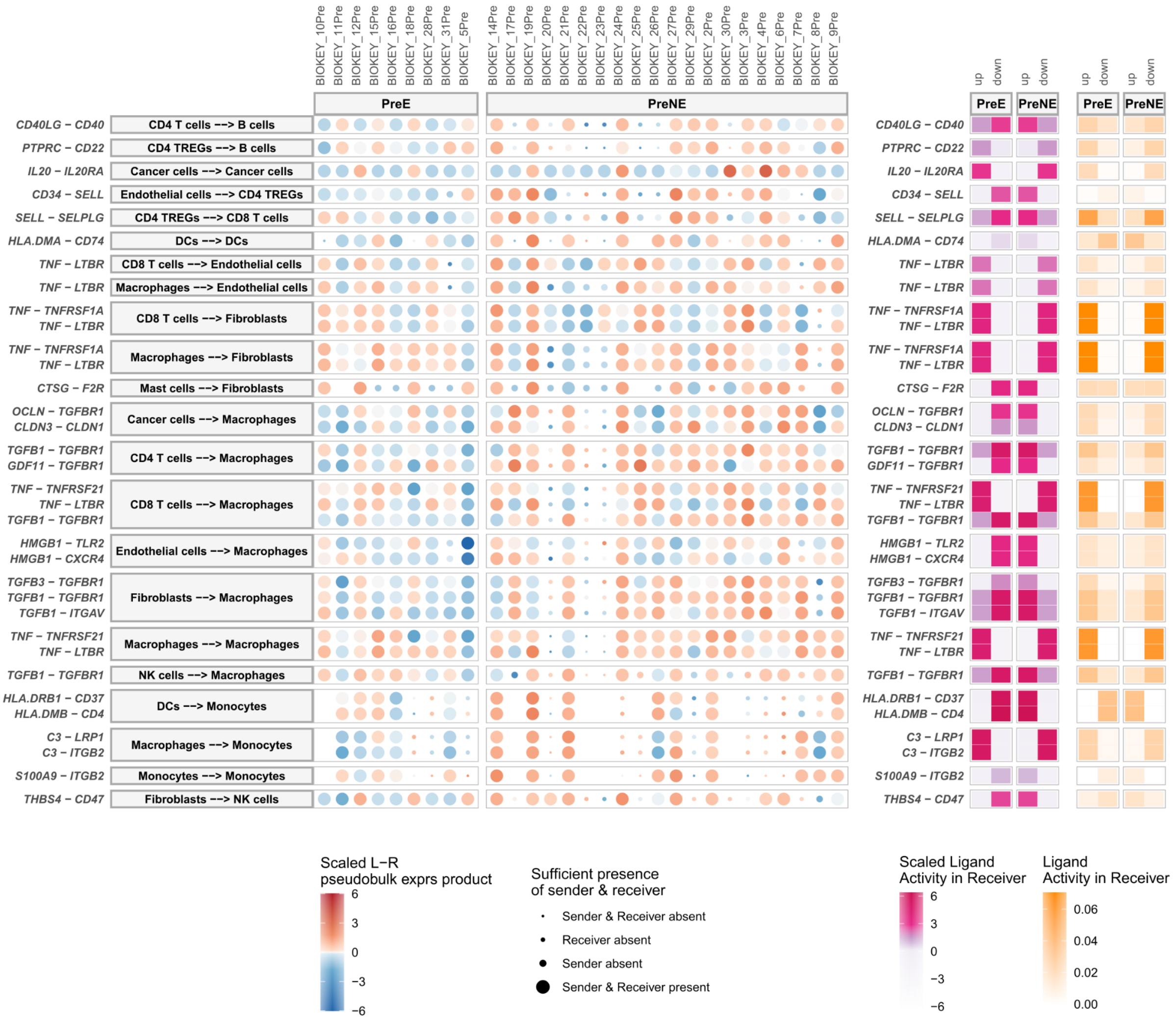


**Supplementary Notes Figure 3.3. | MultiNicheNet prioritizes cell-cell communication patterns specific for patients without clonotype expansion after anti-PD1 therapy.** MultiNicheNet was applied to scRNA-seq from Bassez et al. to compare pre-therapy cell-cell communication between expander (preE) and non-expander patients (preNE). The preNE-specific interactions out of the total top 100 differential interactions are shown. For each interaction, ligand-receptor pseudobulk expression (product of logCPM values) and (scaled) ligand activity values are visualized. Ligand activity scores are the AUPRC scores in predicting the up- or downregulated genes in the preE or preNE group. Scaled ligand activities are z-score normalized ligand activity values, calculated per receiver cell type. The higher these values, the more enriched target genes of a specific ligand are compared to other ligands. The size of the dots indicates whether a sample had enough cells (>= 10) for a specific cell type to be considered for DE analysis. The absence of a dot means there were no cells at all. L-R: ligand-receptor.

Top 50 expansion-specific ligand-receptor interactions between T cells and malignant cancer cells

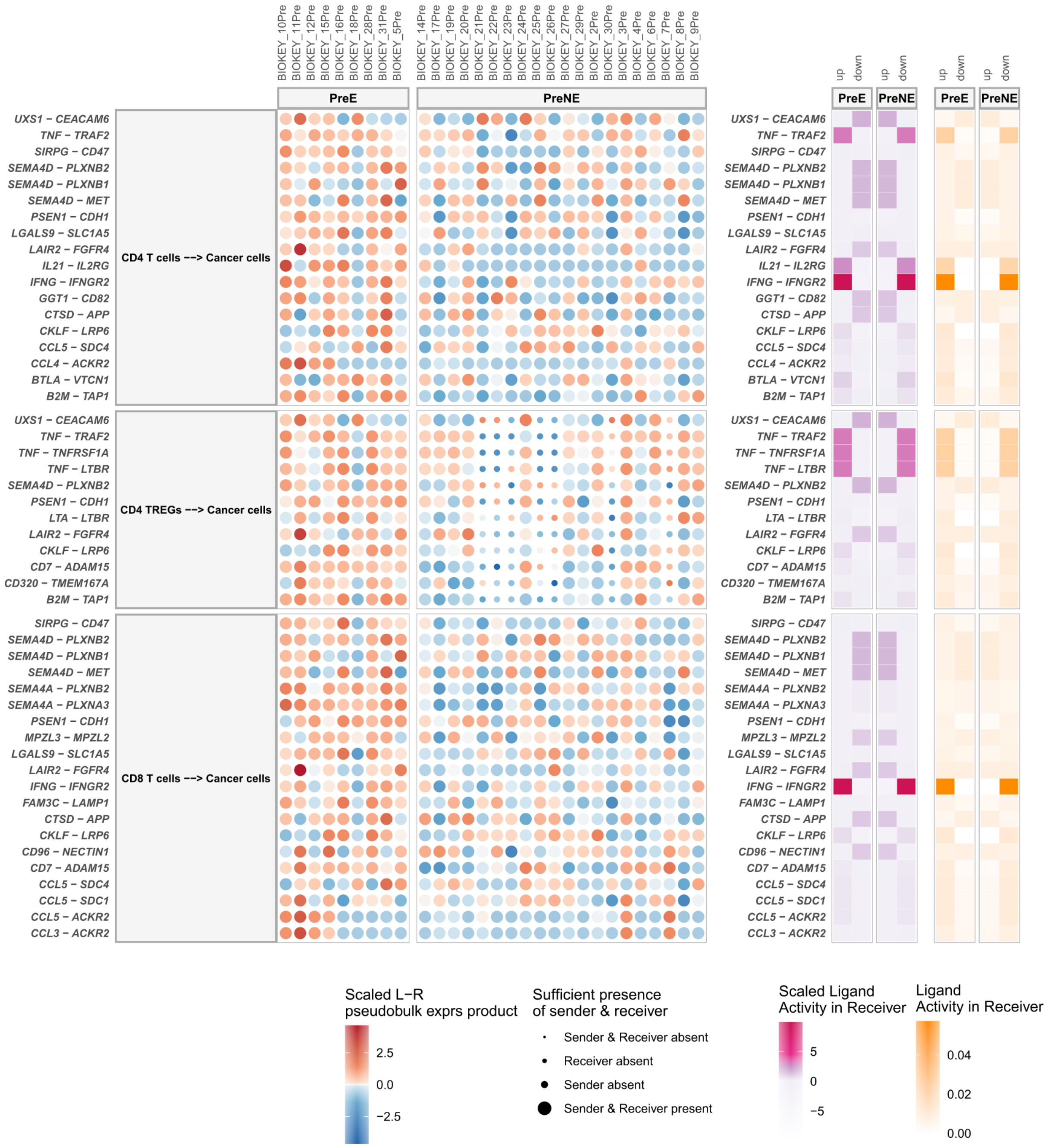


**Supplementary Notes Figure 3.4. | MultiNicheNet prioritizes cell-cell communication patterns between T cells and malignant cells that are specific for patients with clonotype expansion after anti-PD1 therapy.** MultiNicheNet was applied to scRNA-seq from Bassez et al. to compare pre-therapy cell-cell communication between expander (preE) and non-expander patients (preNE). The top 50 preE-specific interactions from T cells to malignant (cancer) cells are shown. For each interaction, ligand-receptor pseudobulk expression (product of logCPM values) and (scaled) ligand activity values are visualized. Ligand activity scores are the AUPRC scores in predicting the up- or downregulated genes in the preE or preNE group. Scaled ligand activities are z-score normalized ligand activity values, calculated per receiver cell type. The higher these values, the more enriched target genes of a specific ligand are compared to other ligands. The size of the dots indicates whether a sample had enough cells (>= 10) for a specific cell type to be considered for DE analysis. L-R: ligand-receptor.

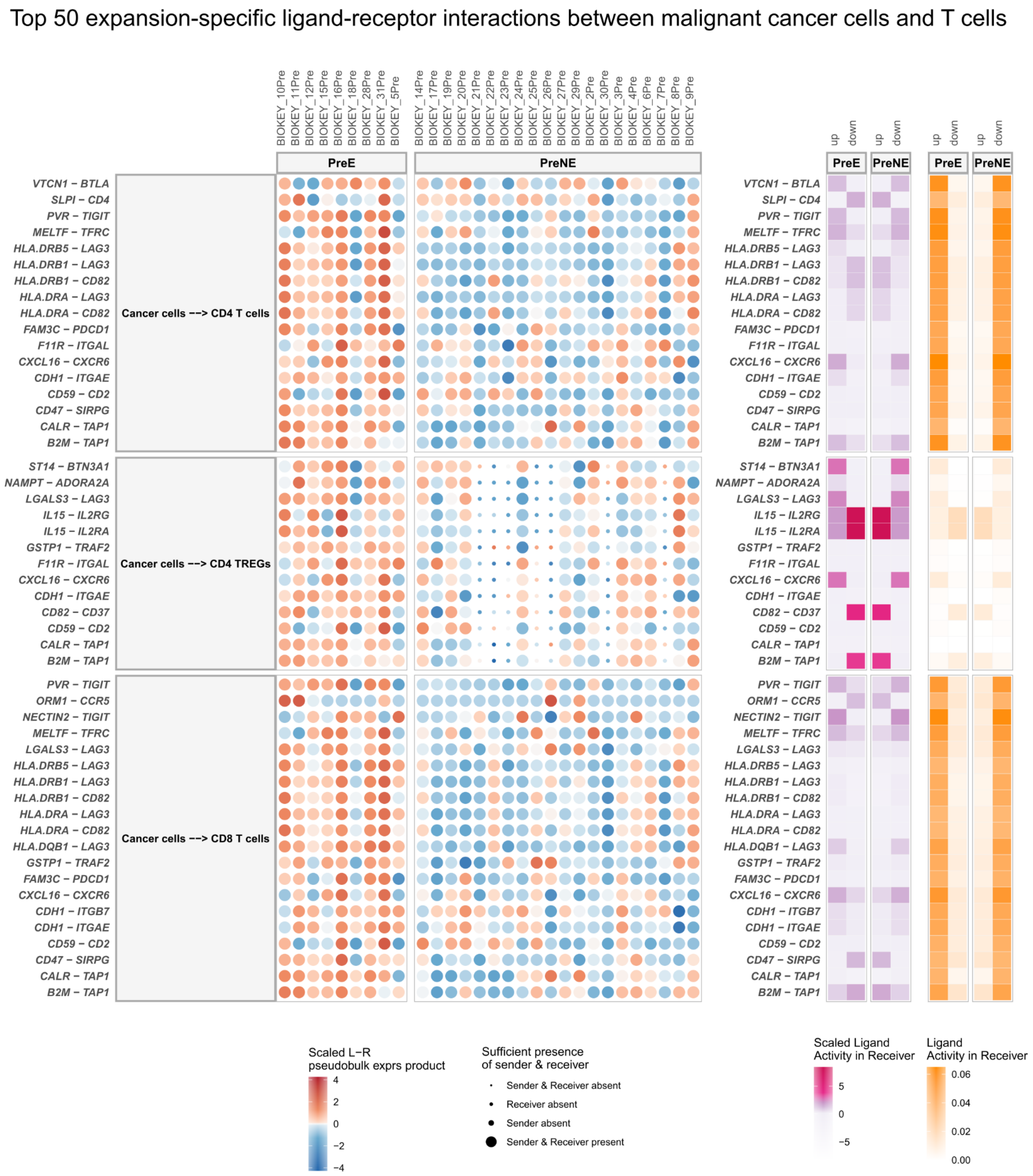


**Supplementary Notes Figure 3.5. | MultiNicheNet prioritizes cell-cell communication patterns between malignant cells and T cells that are specific for patients with clonotype expansion after anti-PD1 therapy.** MultiNicheNet was applied to scRNA-seq from Bassez et al. to compare pre-therapy cell-cell communication between expander (preE) and non-expander patients (preNE). The top 50 preE-specific interactions from malignant (cancer) cells to T cells are shown. For each interaction, ligand-receptor pseudobulk expression (product of logCPM values) and (scaled) ligand activity values are visualized. Ligand activity scores are the AUPRC scores in predicting the up- or downregulated genes in the preE or preNE group. Scaled ligand activities are z-score normalized ligand activity values, calculated per receiver cell type. The higher these values, the more enriched target genes of a specific ligand are compared to other ligands. The size of the dots indicates whether a sample had enough cells (>= 10) for a specific cell type to be considered for DE analysis. L-R: ligand-receptor.

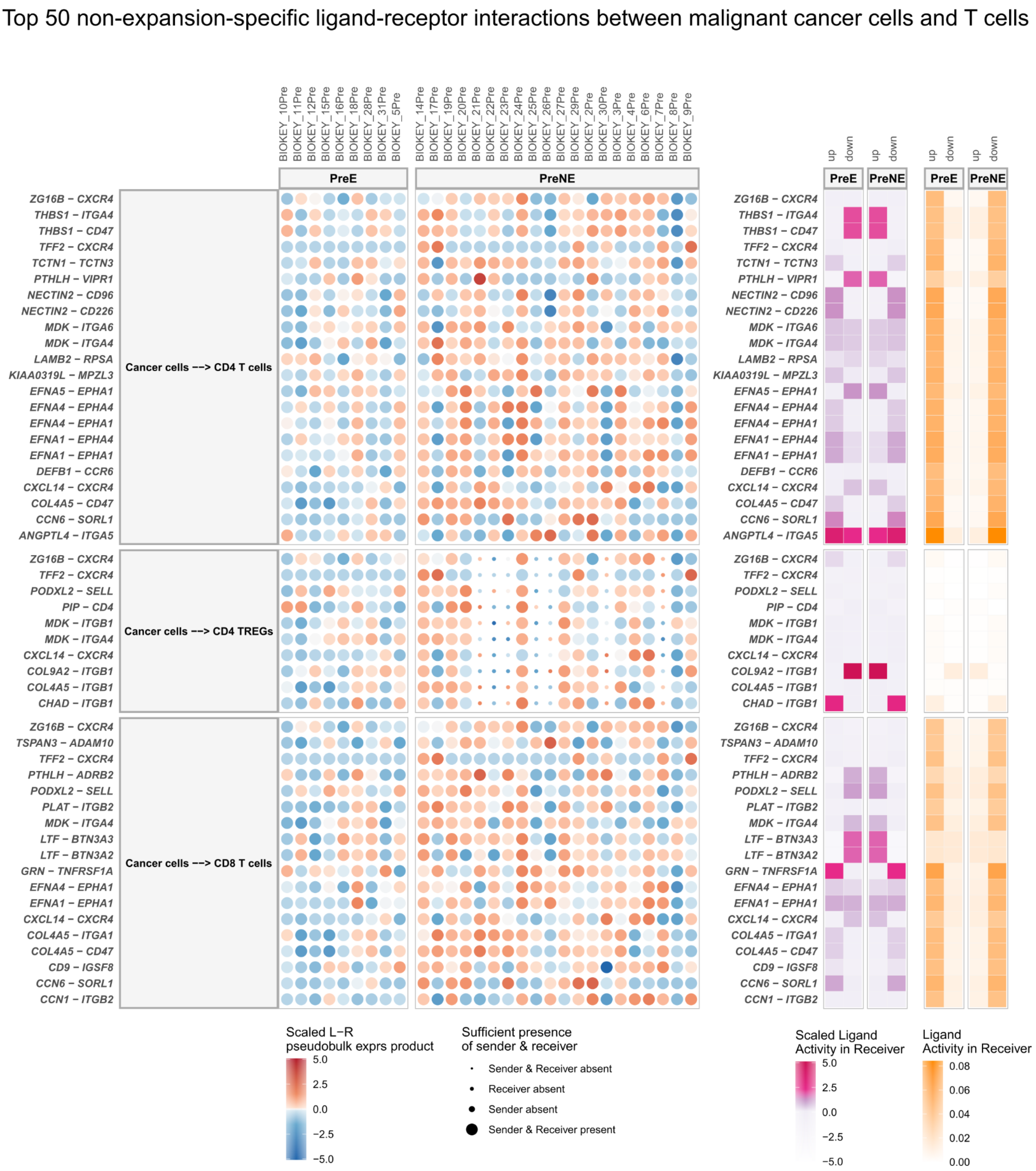


**Supplementary Notes Figure 3.6. | MultiNicheNet prioritizes cell-cell communication patterns between malignant cells and T cells that are specific for patients without clonotype expansion after anti-PD1 therapy.** MultiNicheNet was applied to scRNA-seq from Bassez et al. to compare pre-therapy cell-cell communication between expander (preE) and non-expander patients (preNE). The top 50 preNE-specific interactions from malignant (cancer) cells to T cells are shown. For each interaction, ligand-receptor pseudobulk expression (product of logCPM values) and (scaled) ligand activity values are visualized. Ligand activity scores are the AUPRC scores in predicting the up- or downregulated genes in the preE or preNE group. Scaled ligand activities are z-score normalized ligand activity values, calculated per receiver cell type. The higher these values, the more enriched target genes of a specific ligand are compared to other ligands. The size of the dots indicates whether a sample had enough cells (>= 10) for a specific cell type to be considered for DE analysis. L-R: ligand-receptor.

### Supplementary Note 4: Comparison between different methods

This supplementary note provides a comparative description of several recently developed tools that users can apply for studying differential cell-cell communication from multi-sample data. For a comparison between classic differential cell-cell communication tools, we refer the reader to the comparative table from Armingol et al.[26], the developers of Tensor-cell2cell[1*]. Except for Tensor-cell2cell, these tools are not optimized to run on multi-sample data. Moreover, none of the tools performs differential cell-cell communication by considering both the expression and activity of ligand-receptor pairs. Therefore, we will focus here only on very recently published tools that are not included in that table (except for Tensor-cell2cell itself). We selected tools that perform "multicellular" decomposition of multi-sample scRNA-seq data (DIALOGUE[27], scITD[28], and Tensor-cell2cell[26]), more classic differential cell-cell communication from multi-sample scRNA-seq data (MultiNicheNet and CINS[29]), and tools that enable differential cell-cell communication analysis at single-cell resolution (Scriabin[30] and NICHES[31]). We performed an extensive qualitative comparison of these tools in terms of method goal, use case, methodology, limitations, benefits, and whether prior knowledge of cell-cell communication is used (**Supplementary Notes Table 4.1.**). Our main conclusion is that these hypothesis-generating tools have often different limitations and benefits and can be considered complementary. The choice of which method to use depends on the type of data and research question, but the tools can often also be used in conjunction with each other.

For example, multi-sample methods like scITD, DIALOGUE, Tensor-cell2cell, and MultiNicheNet aggregate expression information per cell-type-sample combination, which is not ideal for rare cell types and subtypes. In contrast, Scriabin and NICHES were developed to overcome this issue. However, these tools provide less insight into inter-sample heterogeneity and are less suited to tackle multi-sample datasets with many samples.

Other differences can be appreciated between DIALOGUE, scITD, Tensor-cell2cell, and MultiNicheNet. The first three tools employ a decomposition strategy that does not require that samples are defined to belong to a certain condition. In contrast, MultiNicheNet performs DE analysis which requires pre-defined conditions of interest to compare. Whereas DIALOGUE and scITD are purely data-driven and uncover genome-wide multicellular programs, Tensor-cell2cell and MultiNicheNet specifically focus on ligand-receptor interactions, which might give more easily interpretable output.

Comparing Tensor-cell2cell to MultiNicheNet as final comparison shows that Tensor-cell2cell has a flexible ligand-receptor scoring scheme that can leverage information on receptor subunit architecture, whereas MultiNicheNet incorporates ligand activities and downstream target gene predictions of ligand-receptor pairs, and also addresses complex experimental designs more fluently.

---

[1*] Correction to that table: NicheNet can also be used for pairwise comparisons between contexts. The "communication score" is then the ligand activity calculated from DE genes in the receiver cell.

**Supplementary Notes Table 4.1. | Comparison between recently developed cell-cell communication tools that can be applied to study differential cell-cell communication from multi-sample data.**

| Method | Goal | Methodology | Prior knowledge of cell-cell communication | Limitations | Additional benefits |
|---|---|---|---|---|---|
| **DIALOGUE[27]** | **Identification of multicellular programs** (MCPs). These consist of gene sets over multiple cell types, that are correlated with each other across different samples/niches.<br><br>Identified MCPs can be associated with covariates of interest (disease states, clinical outcomes, …).<br><br>Required input: single-cell data with multiple samples. | Step 1: Multi-view Penalized Matrix Decomposition. This identifies multicellular latent features correlated across samples.<br><br>[Input for decomposition: feature-sample matrix containing feature values averaged per cell type.]<br><br>Step 2: multi-level hierarchical modeling to identify the genes comprising these latent features.<br><br>Step 3: association of MCPs to covariates of interest. | Not used to identify MCPs, which happens in a purely data-driven way.<br><br>A database of ligand-receptor interactions can be used for post-hoc interpretation of the MCPs: are some ligands/receptors part of the MCPs? | DIALOGUE can suffer from low numbers of cells per cell type, and low numbers of samples.<br><br>Ligand-receptor post-hoc analysis of MCPs is currently not demonstrated in the software tutorials. | Does not require pre-defined sample groups/conditions to compare.<br><br>Can also be used on spatially-resolved transcriptomics data with single-cell resolution.<br><br>Visualizations provide insights into inter-sample heterogeneity. |
| **scITD[28]** | **Identification of multicellular gene expression patterns** with co-varying expression **across** different biological **samples** (very similar to DIALOGUE's MCPs).<br><br>Identified patterns can be associated with covariates of interest ( disease states, clinical outcomes, batch effects, …) and used to stratify heterogeneous samples.<br><br>Required input: single-cell or bulk transcriptomics data with multiple samples. | Step 1: Tucker tensor decomposition. This identifies factors that consist of the multicellular gene expression patterns and their contribution to each sample.<br><br>[Input for decomposition: normalized pseudobulk gene expression values per cell-type-sample combination.]<br><br>Step 2: association of factors to covariates of interest.<br><br>Additional step: associate ligand expression in sender cell types with co-expression modules in receiver cell types (based on across-sample correlation). | A database of ligand-receptor interactions can be used for post-hoc interpretation of the multicellular patterns: are some ligand-receptor interactions mediating these patterns?<br><br>Use of the ligand-target prior knowledge model of NicheNet to assess whether some of scITD's ligand-module links are supported by prior knowledge. | scITD can suffer from low numbers of cells per cell type, and low numbers of samples.<br><br>Unclear whether it can handle cell types that are missing in one or more samples.<br><br>The inference of ligand-module links cannot be performed for autocrine communication patterns. | Does not require pre-defined sample groups/conditions to compare.<br><br>Appropriate handling of batch effects.<br><br>Identification of shifting cell subtype compositions.<br><br>Visualizations provide insights into inter-sample heterogeneity. |

| Method | Goal | Methodology | Prior knowledge | Limitations | Additional benefits |
|---|---|---|---|---|---|
| **Tensor-cell2cell**[26] | **Identification of communication patterns with co-varying expression across samples/contexts.** The prioritized communication patterns are unique combinations of **ligand-receptor** interactions **between sender and receiver cell types**.<br><br>Identified patterns can be associated with covariates (disease states, ... )<br><br>Required input: bulk or single-cell data with multiple contexts/samples. | Step 1: Tensor component analysis. This step identifies factors that account for the correlation structure across samples/contexts. A factor can be interpreted as a communication pattern that changes across contexts.<br><br>[Input for decomposition: 4D-tensor with the ligand-receptor pair communication scores (e.g., expression product) between each sender-receiver cell type pair, across all contexts.]<br><br>Step 2: interpretation of the factor loadings to infer the most important ligand-receptor pairs and sender-receiver pairs per factor.<br><br>Step 3: association of factors to covariates of interest. | Use of a ligand-receptor database (flexible, can handle multi-subunit architecture). | Tensor-cell2cell is not optimized to handle cell types and/or ligand-receptor pairs that are missing in one or more samples.<br><br>Unclear how robust Tensor-cell2cell is for low numbers of cells per cell type and low numbers of samples/contexts.<br><br>Current software implementation does not provide downstream signaling interpretation of ligand-receptor pairs (e.g., target gene predictions), nor prioritization based on ligand activity. | Does not require pre-defined groups/conditions to compare.<br><br>Can use the output (ligand-receptor communication score) of other existing cell-cell communication tools as input.<br><br>Visualizations provide insights into inter-sample heterogeneity. |
| **MultiNicheNet** | **Identification of communication patterns that differ between conditions**. The prioritized communication patterns are **ligand-receptor** interactions and their predicted **target genes, between sender and receiver cell types**.<br><br>Required input: single-cell transcriptomics data with multiple samples, with at least two pre-defined conditions | Multi-criteria prioritization framework that can consider the differential expression, cell-type specific expression, and target gene enrichment (ligand activity) to prioritize ligand-receptor pairs.<br><br>DE analysis is currently performed through pseudobulk aggregation followed by edgeR analysis | Use of a ligand-receptor database and a corresponding NicheNet ligand-target prior knowledge model | MultiNicheNet can suffer from low numbers of cells per cell type, and low numbers of samples.<br><br>MultiNicheNet cannot consider condition-specific cell types. | Appropriate handling of batch effects, which facilitates its use on integrated atlas data.<br><br>Enables addressing complex experimental designs and questions in a simple way.<br><br>Visualizations provide insights into inter-sample heterogeneity and cover all aspects of the predictions and multiple levels of cell-cell communication.<br><br>Prediction of intercellular regulatory networks is possible because prioritized ligand-receptor pairs are linked to targets. |

| Method | Goal | Methodology | Prior knowledge | Limitations | Additional benefits |
|---|---|---|---|---|---|
| **CINS**[29] | **Identification of interacting cell types** and **prioritization** of the **ligands** involved in the predicted interactions.<br>(both within and between conditions).<br><br>Required input: single-cell data with multiple samples, preferably case-control. | Step 1: Bayesian network analysis to predict interactions between cell types, assuming that cell types with co-varying abundances are interacting.<br><br>Step 2: ligand-target regression modeling to prioritize ligands by explaining changes in target genes as a function of changes in their activating ligands. | Use of the ligand-target prior knowledge model of NicheNet | Interactions between cells of the same type cannot be identified<br><br>CINS can miss important cell-cell interactions when there is a low number of cells and a low number of samples.<br><br>The ligand-target regression model does not consider the presence of the receptor(s) of the ligands in the receiver cell type.<br><br>Software provides only limited downstream visualizations. | |
| **NICHES**[31] | Recover ligand-receptor cell-cell communication patterns at **single-cell resolution.**<br><br>Required input: single-cell or spatial transcriptomics data with one or multiple samples. | Step 1: convert traditional single-cell transcriptomics atlases into single-cell signaling atlases. Per ligand-receptor pair, the expression product is calculated for each considered cell-cell pair (similar to Scriabin's cell-cell interaction matrix).<br><br>Step 2: apply downstream single-cell analysis tools to find ligand-receptor pairs between cell (sub)types and differences between conditions. | Use of a ligand-receptor database (flexible, can handle multi-subunit architecture). | Random subsampling of cell-cell pairs necessary could lead to loss of information.<br><br>No downstream signaling interpretation of ligand-receptor pairs.<br><br>Currently, not optimally adapted to multi-sample settings. The default differential analysis (as demonstrated in the software) pools cells from multiple samples from the same condition together. | Resolution at the single-cell level (with or without denoising) has the big advantage that intra-celltype heterogeneity in cell-cell communication can be picked up.<br><br>Allows users to explore changes in signaling due to the addition or loss of cell populations.<br><br>Integration with pseudotime inference. |

| Method | Goal | Methodology | Prior knowledge | Limitations | Additional benefits |
|---|---|---|---|---|---|
| **Scriabin**[30] | Recover biologically meaningful cell-cell communication patterns at **single-cell resolution.**<br><br>Required input: single-cell transcriptomics data with one or multiple samples. | *Main workflow (for small datasets): cell-cell interaction workflow:*<br>Step 1: convert traditional single-cell transcriptomics dataset into a cell-cell interaction matrix. Per ligand-receptor pair, the geometric expression product is calculated for each considered cell-cell pair. These interactions can be weighted by including NicheNet ligand activity predictions.<br>Step 2: apply downstream single-cell analysis tools to find ligand-receptor pairs between cell (sub)types and differences between conditions.<br><br>*Second workflow (for comparative analyses): summarized interaction graph workflow:* identification of cell-cell pairs with different total communicative potential between samples. This forms an intelligent sampling of cell pairs, which is necessary for a scalable analysis through the main workflow.<br><br>*Third workflow (for analyses with many samples [> 20]): interaction program discovery workflow*: identification of modules of co-expressed ligand-receptor pairs, followed by examination of sample differences. | Use of a ligand-receptor database and a corresponding NicheNet ligand-target prior knowledge model | Cell pair sampling through the summarized interaction graph workflow is not robust to situations where ligand-receptor pair mechanisms of cell-cell communication change between cell-cell pairs without changing the overall magnitude of cell-cell communication.<br><br>Hard threshold based on NicheNet ligand activities may limit the discovery of understudied interactions.<br><br>Currently, not optimally adapted to multi-sample settings. The interaction program discovery module is the only recommended module when there are more than 20 samples in the dataset. It is also not clear how it would handle batch effects in integrated atlas data. | Resolution at the single-cell level (with or without denoising) has the big advantage that intra-celltype heterogeneity in cell-cell communication can be picked up. |
